# Functional selection in a population of synthetic cells with a minimal metabolism

**DOI:** 10.1101/2025.08.25.672132

**Authors:** T. Di Meo, L. Bunel, G. Ragala, M. Van Tongeren, R. Sieskind, C. Danelon, Y. Rondelez

## Abstract

Various synthetic microcompartment systems have been developed to mimic key features of living cells. Here, we focus on artificial cells that capture their capacity to serve as vessels for Darwinian evolution. We assemble micro-compartmentalized In Vitro Transcription-Translation-Replication systems containing a minimal genome, a basic metabolic pathway, a reconstituted protein expression machinery, and a simple DNA replication module, wired in a positive feedback loop. The minimal genome encodes the enzyme deoxyribonucleoside kinase (DNK) whose expression, and then metabolic activity, is required for the genome’s replication. We show that these compartments act as minimal Darwinian elements by filtering out non-functional genotypes. We track individual replicators from a library of 42 genetic variants to reveal the system’s dynamics at both the population and the single replicator levels. At the population level, we extract the fitness function, which links a genome’s metabolic efficiency to its selective success, considering co-encapsulation and hitch-hiking effects. At the individual replicator level, we observe a bimodal distribution of replication yields and propose a mixed model with an inter-droplet heterogeneity with presence or absence of a metabolic feedback loop on the replicator. In addition, we leverage this autonomous self-selection loop to generate a high-resolution mutational map of the DNK enzyme.

## INTRODUCTION

Darwinian evolution operates via natural selection: replicators encoding beneficial phenotypes replicate more efficiently and gradually outcompete others in a mixed population. Despite the fundamental importance of this process from life’s origins to its current diversity, it has been historically challenging to reproduce and study evolution *in vitro*. As Gánti articulated, a minimalistic Darwinian system would still require three integrated components: (i) a closed compartment or boundary separating it from the environment, (ii) a source of building blocks and energy (a basic metabolism), and (iii) an information-carrying replicating molecule (Ganti, 2003). Moreover, the genetic molecule should be present in only a discrete, small number of molecular copies per compartment. This stochastic regime is a prerequisite for effective selection in a genetically diverse population (Ichihashi et al., 2013; Sakatani et al., 2018; Tawfik & Griffiths, 1998), higher ploidies leading to less efficient selection (Zadorin & Rondelez, 2019).

Tawfik and Griffiths pioneered the concept of Poissonian DNA partitioning in emulsions to obtain a genotype–phenotype linkage *in vitro* and demonstrated the possibility of establishing a feedback between the two (Tawfik & Griffiths, 1998). They encapsulated a linear DNA encoding a methyltransferase enzyme in water-in-oil droplets made from a cell extract. The expressed methyltransferase would methylate its own encoding gene inside the droplet. After breaking the emulsion, a selection step was applied in bulk by exposing the purified DNA to restriction enzymes: methylated genes being protected from digestion, only DNA from droplets containing active enzymes survived and were amplified by PCR. Through repeated cycles of such compartmentalized DNA self-modification and bulk selection, they successfully enriched the active methyltransferase genes out of a mixed population.

An important next step was achieving concurrent genome replication and expression inside the compartments. Based on Spiegelman’s seminal work with the Qβ RNA replicase (Mills et al., 1967), Ichihashi and colleagues encapsulated an RNA genome encoding the Qβ replicase in Protein production Using Recombinant Elements (PURE) system-containing microdroplets (Ichihashi et al., 2013). This system showed genotype-dependent replication from stochastically partitioned replicators, and therefore Darwinian dynamics. It, however, became rapidly dominated by parasitic short RNAs replicating faster than the full-length replicator. Nonetheless, this work provided the first observation of emergent evolutionary dynamics entirely *in vitro*, with functional and parasitic replicators co-evolving in a cell-free compartmentalized system (Furubayashi et al., 2020; Ichihashi et al., 2013, 2015; Mizuuchi & Ichihashi, 2018).

Mizuuchi and Ichihashi later tried to augment the Qβ replicase system with a minimal metabolic function (Mizuuchi & Ichihashi, 2018). They designed two co-replicating RNA genes in one compartment: one encoding the Qβ RNA polymerase and another encoding nucleoside diphosphate kinase (NDK) to regenerate NTPs, since the PURE system lacks that activity. This study revealed a limitation of RNA based systems: the stringent structural requirements of the RNA genomes (for replication) severely constrained their evolution and reduced modularity.

DNA-based replication systems offer a potential advantage over RNA replicators: they can more efficiently segregate the two functions of the genetic material - replication and gene expression (Takeuchi et al., 2011). Recently, minimally complex DNA self-replicators have been developed by combining viral DNA replication proteins with a PURE expression system (Abil et al., 2024; Sakatani et al., 2018, 2019; Van Nies et al., 2018). One successful approach uses the reconstituted bacteriophage φ29 DNA replication machinery composed of four protein elements (Blanco et al., 1994; Mencía et al., 2011). It includes a mesophilic DNA polymerase with strong strand-displacement activity, compatible with PURE’s operating conditions (Abil et al., 2022, 2024; Van Nies et al., 2018). To date, the only published examples of Darwinian evolution involving replication of a DNA genome in an artificial cell is the evolution of the φ29 DNA polymerase gene itself (Abil et al., 2024; Sakatani et al., 2018), achieved using liposomes for compartmentalization.

To our knowledge, a compartmentalized self-replicating DNA system enabling Darwinian evolution of a metabolic function *in vitro* has not yet been reported. In this work, we build a micro-compartmentalized system including three key elements: a minimal genome encoding for the deoxyribonucleoside monophosphate kinase (DNK) from phage T5 converting dNMP to dNDP in a synthetic dNTP metabolic pathway, a reconstituted protein expression system and a DNA replication machinery. These elements are wired in a positive feedback loop, whereby *dnk* expression and metabolic activity is required for *dnk* replication. We show that DNK activity-conditional replication with thousand-fold amplification can be observed from single-copy genetic elements encapsulated into picolitre droplets. To track selection dynamics, we follow the replicative fitness of a population containing 42 genetic variants with various catalytic activities and expression strengths. This reveals the fitness function that links the replicators’ metabolic efficiencies to their evolutionary success in this minimalist Darwinian system. Finally, we apply this approach to generate a high-resolution mutational map of the DNK enzyme.

## RESULTS

### Coupled genetic replication and gene expression

The objective of this study was to build a synthetic cell combining the full central dogma with a minimal metabolism. We thus needed a system that can support transcription and translation of a gene, as well as its replication. Among the various *in vitro* DNA replication systems described to date, the combination of the Protein production Using Recombinant Elements (PURE) system (Shimizu et al., 2001) with the φ29 bacteriophage replication machinery (Mencía et al., 2011) represents one of the few isothermal platform capable of exponential amplification of long linear DNA templates from sub-picomolar concentrations, while remaining compatible with simultaneous transcription and translation in a single reaction (Abil et al., 2024; Van Nies et al., 2018).

This In Vitro Transcription-Translation-Replication (IVTTR) system reconstitutes four essential components of the original φ29 replication machinery: the DNA polymerase (DNAP, encoded by gene *p2*), the terminal protein (TP, *p3*) acting as polymerization primer, the single-stranded DNA binding protein (SSB, *p5*), and the double-stranded DNA binding protein (DSB, *p6*) (Kamtekar et al., 2006; Salas et al., 2016).

The replication cycle is shown in Figure 1a. Replication is initiated when the Terminal Protein–DNA polymerase (TP–DNAP, encoded by genes p2 and p3 respectively) complex binds specific terminal sequences (OriL and OriR) of a linear DNA template. Unlike canonical DNA polymerases, φ29 DNAP does not require a DNA or RNA primer. Instead, it uses the hydroxyl group of Serine 232 on the TP as the priming site for covalent attachment of the first deoxynucleotide, dAMP, via a phosphodiester bond (Kamtekar et al., 2006). This yields a TP-capped DNA product, the natural form of the genome virus, which has been shown to enhance re-initiation by facilitating further TP–DNAP recruitment (de Vega et al., 1997). Interestingly, it has been shown that in absence of TP protein, a phosphorylated 5’ end was a prerequisite to allow significant replication (Blanco et al., 1994; Mencía et al., 2011). In the following, we will refer to linear DNA sequences flanked by phosphorylated or TP-capped OriL and OriR as replicators.

**Figure 1.**
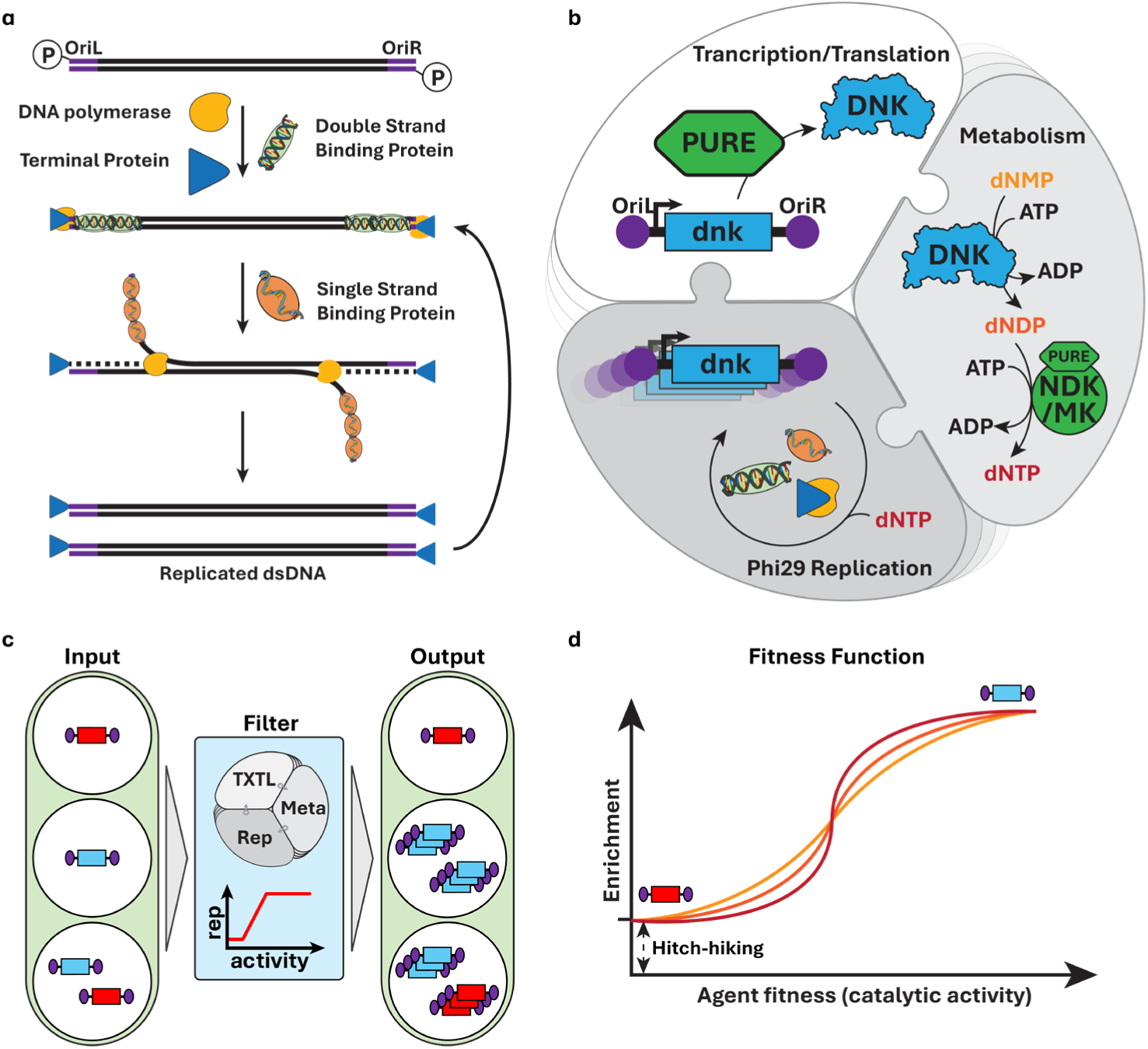
Minimal Darwinian compartments. **a**. φ29 replication cycle. Origins of replication flanking replicators are bound to DSB multimers, unwinding the DNA helix. DNAP-TP complex recognizes phosphorylated end of DNA. DNAP covalently attaches first adenine to Terminal Protein and starts copying while displacing DNA strand. Displaced DNA strand is coated by SSB until completion of the elongation step. TP-caped DNA allows more efficient re-binding of DNAP-TP complex to undergo another cycle. **b**. Schematic representation of a self-selecting IVTTR system inside a compartment. First lobe: DNK-encoding gene (dnk, blue) is flanked by origins of replications (OriL and OriR, purple dots). PURE Transcription/Translation system produces DNK enzyme. Second lobe: DNK enzyme produces dNDP from dNMP, then kinases NDK and MK from PURE system convert dNDP to dNTP. Third lobe: dNTP fuels the φ29 replication machinery composed of DNAP, TP, SSB and DSB proteins to replicate the dnk gene. If replication happens in a time frame where the PURE system is still active, the amplified gene can also be expressed by PURE system (back to the first lobe). **c**. Schematics of in vitro selection experiment. A mixed population of active (blue) or inactive (red) replicators is randomly partitioned in an emulsion. **d**. Darwinian selection applied to the population results in enrichment that depends on genotype performance and hitch-hiking effects induced by co-encapsulation, described as the fitness function. The shape of the curve (curve color) depends on the replication function at variant level and the composition of the population.

Two additional proteins facilitate the replication process. Unwinding of the origins by Double Stranded DNA binding protein (DSB) multimer binding is necessary for the polymerase to efficiently initiate strand-displacement synthesis (Alcorlo et al., 2024; Salas et al., 2016). The single DNA strands that are displaced by the polymerase are stabilized and protected from degradation *in vivo* by the Single Stranded DNA Binding protein (SSB), which has also been shown to increase replication efficiency *in vitro* (Blanco et al., 1994). Using this reconstituted system, replicators up to ∼20 kb have been amplified *in vitro* (Blanco et al., 1994; Mencía et al., 2011).

Danelon and colleagues demonstrated that φ29-based replicators can undergo simultaneous DNA replication and protein expression in PURE (Abil et al., 2022, 2024; Van Nies et al., 2018). More important, they showed that TP and DNAP could be expressed *in situ* and then catalyze the replication of their own encoding DNA, establishing the foundation of a self-sustained IVTTR system, which we expand upon in this work.

### Minimal metabolic pathway

The PURE system used in IVTTR contains the necessary components for gene expression: an energy source in the form of creatine phosphate, the building blocks for synthesis of messenger RNAs and proteins and the enzymes responsible for ATP regeneration, NTP homeostasis, and transfer RNA aminoacylation. However, it lacks a dNTP generation mechanism. This is the function we implemented in this study, via the introduction of a gene encoding DNK enzyme.

From a kinetic point of view, we expect the full system to work as follows: in the early phase, the PURE mixture produces the DNK enzyme encoded on the replicator (Figure 1b, white and light grey lobes), which triggers the buildup of dNTP and subsequent replication of DNA, creating a positive feedback loop (Figure 1b, dark grey lobe) as more DNA generates more enzyme. Eventually, IVTTR reaches a plateau when resources are exhausted, or the system’s efficiency drops.

### Compartmentalization

We encapsulated the reactions by creating monodisperse water-in-fluorinated oil emulsions. The use of fluorinated oil greatly limits diffusion of small molecules between droplets, thereby preserving genotype–phenotype linkage (Agresti et al., 2010; Holtze et al., 2008). We chose an appropriate dilution for our replicator DNA such that each droplet receives a discretized number of DNA molecules following a Poisson distribution. In a monodisperse population of microdroplets, having one, two, or three DNA molecules per compartment corresponds to 1×, 2×, or 3× a “quantum concentration”, which we define as the concentration corresponding to one molecule in one droplet. This quantum concentration is inversely proportional to the compartment size, which falls in the low-picomolar range in our study. Thus, compartment size directly influences the rate of production of DNK and the accumulation of metabolites. In addition, stochastic partitioning created the opportunity of hitch-hiking, where low fitness replicators can benefit from getting randomly co-encapsulated with variants supporting more active metabolism.

### Kinase activity is necessary for replication

Our first objective was to make DNA replication conditional on the metabolic cascade for dNTP production. The PURE system already contains three kinases: Creatine Kinase (CK), myokinase (MK) from rabbit muscle and nucleoside-diphosphate kinase (NDK) from *E. coli* (Shimizu et al., 2001). These enzymes maintain activated nucleotide levels by transferring phosphate from ATP to (d)NDPs, supporting (d)NTPs homeostasis. NDK and MK are already known to be able to transfer the phosphate from ATP to dNDP to form dNTP (Almaula et al., 1995; Barthel & Walker, 1999; Levit et al., 2002; Lu & Inouye, 1996), meaning we cannot use the replacement of dNTP by dNDP to create a selection pressure. However, their activity toward dNMP has not been reported.

We therefore selected an enzyme to realize this function. The deoxyribonucleoside monophosphate kinase from bacteriophage T5 can phosphorylate all four dNMP to dNDP (Mikoulinskaia et al., 2002), which should enable DNA replication in a PURE mixture where only dNMP, and no dNTP, have been supplied. It was possible, however, that PURE itself presents a low level of dNMP kinase activity from contaminant enzymes or promiscuous activities.

To confirm the dependence of dNMP-supported replication on DNK activity in PURE, we used a linear DNA construct carrying a GFP gene under a T7 promoter and flanked by φ29 origin sequences. In the presence of P2, P3, P5, P6 and dNTP, this construct is a replicator, noted <*gfp*>, where the chevrons represent OriL and OriR (Figure 2a). Its replication can be monitored in two ways: (1) qualitatively, by observing GFP fluorescence increase over time (Supplementary Figure 1), and (2) quantitatively, by measuring the fold-increase in <*gfp*> DNA via qPCR at the end of the reaction.

**Figure 2.**
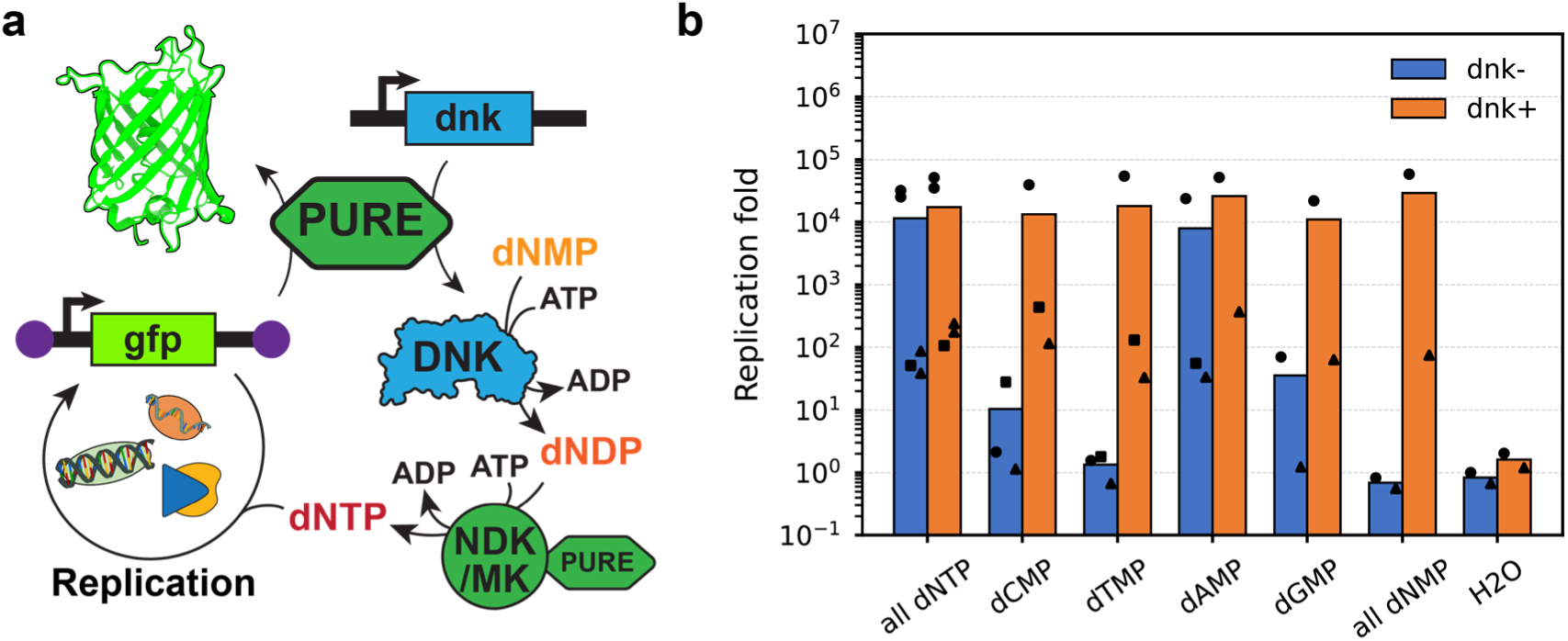
DNK-dependent <*gfp*> replication in bulk. **a**. Schematic of the <*gfp*> reporter (green) with φ29 origins of replication (purple) on both ends. Replication of <*gfp*> is allowed only if the four dNTP are present through expression of the *dnk* gene, conversion of dNMP to dNDP by DNK, and production of dNTP from dNDP by the NDK/MK enzymes present in PURE system. Replication of <*gfp*> leads to increased GFP production and, thus, an increase in fluorescence (Supplementary Figure 1). **b**. End-point quantification of <*gfp*> in bulk IVTTR by qPCR. A unique IVTTR mastermix was split in multiple aliquots and each aliquot was spiked with a different dNTP mixture, where one or all dNTP was replaced with its corresponding dNMP, in absence (-) or presence (+) of 25 pM of *dnk*. Symbols indicate tubes from the same mastermix. All reactions contained 100 pM of pUC_OriLR_p2p3 plasmid.

In this study, the φ29 replication machinery itself is decoupled from the replicating gene of interest: genes encoding DNAP and TP were added in the non-replicative form of a circular plasmid (pUC57_p2p3), while the proteins SSB and DSB were provided in purified form, because relatively high concentrations are necessary to support φ29-based replication (Blanco et al., 1994; Mencía et al., 2011).

We first verified that addition of DNK rescues <*gfp*> amplification in bulk reactions when some dNTP were omitted. To do so, we replaced either one or all four dNTP, with the corresponding dNMP, in presence or absence of *a non-replicating linear DNA containing the dnk gene*. A negative control was assembled without deoxyribonucleotides. Reactions were run at 33 °C for 20 hours and the final <*gfp*> DNA yield was quantified by qPCR (Fig 2b). As expected, we observed strong replication for the all-dNTP condition even in the absence of *dnk*, whereas no DNA replication occurred without dNTP, confirming that the commercial PURE system does not contain a significant contamination of all dNTP.

When dATP was substituted by dAMP, <*gfp*> replication was still detected, even without *dnk* gene. This suggests that PURE system contains either dATP (or dADP) contamination, or, most likely, that some of its enzymes (e.g. myokinase or NDK) can promiscuously convert dAMP to dATP. In contrast, when dTTP, dCTP, or dGTP was omitted and their monophosphate analog was added, replication of <*gfp*> was established only in the presence of *dnk*. This result demonstrates that DNA amplification can be connected to the activity of DNK, which phosphorylates dTMP, dCMP, and dGMP efficiently enough to support replication.

It is noteworthy that bulk IVTTR assays showed substantial run-to-run variability, even within a single experimental run (same master mix split in different tubes). For example, in different preparation batches, the positive control (all dNTP) sometimes yielded poor <*gfp*> amplification, while for other batches ∼100-fold or even ∼25,000-fold replication was reached. We have not fully identified the cause of this discrepancy. We suspect that heterogeneity in the added P5 and P6 proteins (e.g., P5 tends to jellify at low temperature), combined with challenges in pipetting very small volumes of viscous solutions, contribute significantly to the observed variability. In addition, minute differences in the initial conditions can be magnified due to the exponential nature of the reaction. Finally, the sporadic emergence of short “parasite” DNA replicons (which amplify faster than full-length <*gfp*>) is a known issue in isothermal DNA amplification and could lower <*gfp*> yields via resource competition. This issue is normally alleviated in compartmentalized systems which are less prone to catastrophic parasite invasion (Abil et al., 2024; Ichihashi et al., 2013; Manfred & Peter, 1977; Matsumura et al., 2016; Matsuura et al., 2002). Therefore, we went on to assay our IVTTR system in microfluidic-based water-in-oil droplets.

### Droplet IVTTR from single <*rfp-dnk*> replicator

In the bulk experiment described above, *dnk* was supplied as a separate non-replicating DNA, while the replicator encoded the reporter protein GFP. In a true metabolic self-replication scheme, the replicator must encode the metabolic activity itself, and an autocatalytic loop connecting this enzymatic activity to replication must be formed (Figure 1b). In addition, for efficient selection dynamics, the initial replicator concentration must be close to the quantum concentration, here at about 1 pM. Therefore, we set out to visualize replication in microfluidic droplets containing a single DNK-encoding replicator molecule on average as input DNA. We constructed <*rfp-dnk*>, a gene fusion encoding mCherry (red fluorescent protein) and DNK. Replication is expected only if DNK activity inside the droplet is sufficient and should be accompanied by an increase in mCherry fluorescence.

IVTTR reactions with or without dTTP were emulsified with an average ratio of <*rfp-dnk*> copies per droplet (the Poissonian parameter λ) of around 1.5. Upon imaging the emulsions, we observed a mixture of dark droplets (no mCherry fluorescence) and bright droplets (strong mCherry expression). The fluorescence intensity of the positive droplets spanned a broad range (Figure 3a), consistent with previous studies on the stochastic expression of GFP in microcompartments. Indeed an increase in phenotypical dynamic range was observed when moving from PURE-only to IVTTR (Abil et al., 2022). A broad positive peak has also been observed in stochastically encapsulated Replication-Cycle Reaction (RCR), using a DNA-intercalating dye (Ueno et al., 2021).

**Figure 3.**
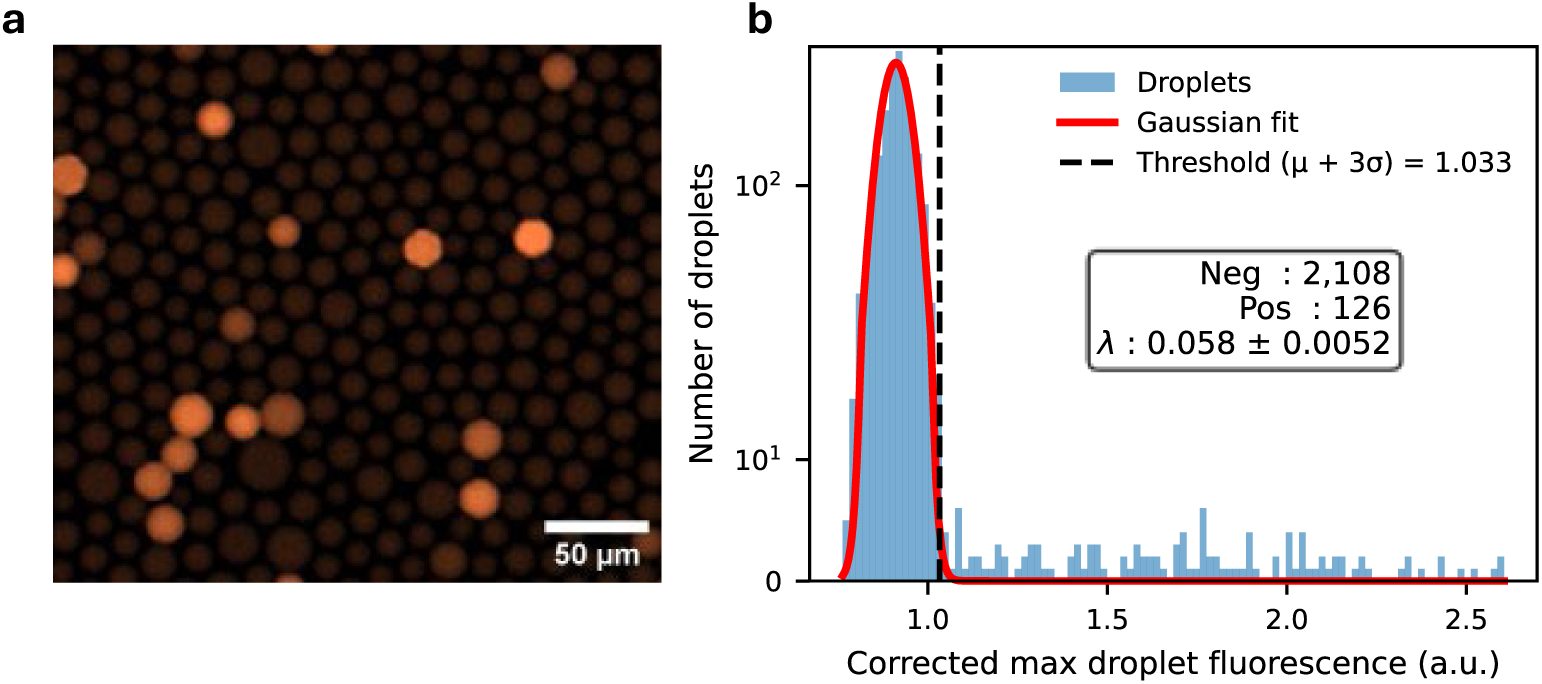
DNK-dependent single replicator self-ampliciaction in droplets. **a.** Fluorescence image of emulsion after incubation of IVTTR using <*rfp-dnk*> wild type in absence of dTTP, replaced by dTMP. Accumulation of mCherry indicates replication. **b**. Histogram of droplet fluorescence with a gaussian fitting around the median of negative droplets. Positive droplets corresponds to median + 3 SD (vertical dashed line).

We defined the threshold of positive droplets as median plus 3x standard deviation of empty droplets peak (Figure 3b), giving an observed positive droplet fraction of 5-6%. Since the expected value from poissonian partitioning is around 80%, we can estimate that ∼1/16^th^ of the compartmentalized DNA constructs have significantly replicated inside their droplet.

To supplement these observations, samples were extracted, and DNA was quantified by qPCR using primers targeting the *dnk* gene. We obtained a final replicator concentration of 680±90 pM, corresponding to 680-fold amplification. However, if we attribute this increase solely to the 5-6% droplets with significant fluorescence level, then each of those droplets experienced on the order of a 12,000-fold amplification of its DNA. This high value is consistent with some of the bulk observations (Figure 2b).

To further explore the behavior of <*rfp-dnk*> in this system, we characterized the selection dynamics in a mixed replicator population encoding various levels of metabolic activity. Specifically, our goal was to quantify the selective pressure experienced by the various genotypes and the relationship between total enzymatic activity, which we call genotype performance, and replication, taking into account replicator co-encapsulation.

### Selection dynamics in a population of genetically diverse replicators

To characterize the fitness function of our system, which we define as the relationship between enzymatic activity and DNA amplification within a given population, we aimed to explore how both specific catalytic activity and protein expression level contribute to selection outcomes. To do so, we took a combinatorial approach by constructing a “mini library” of genotypes. We constructed a small collection of variants that combinatorically varies both enzyme catalytic activity and expression level, so that we can infer the fitness function without needing an exact prior measurement of each variant’s activity (Figure 4a).

**Figure 4.**
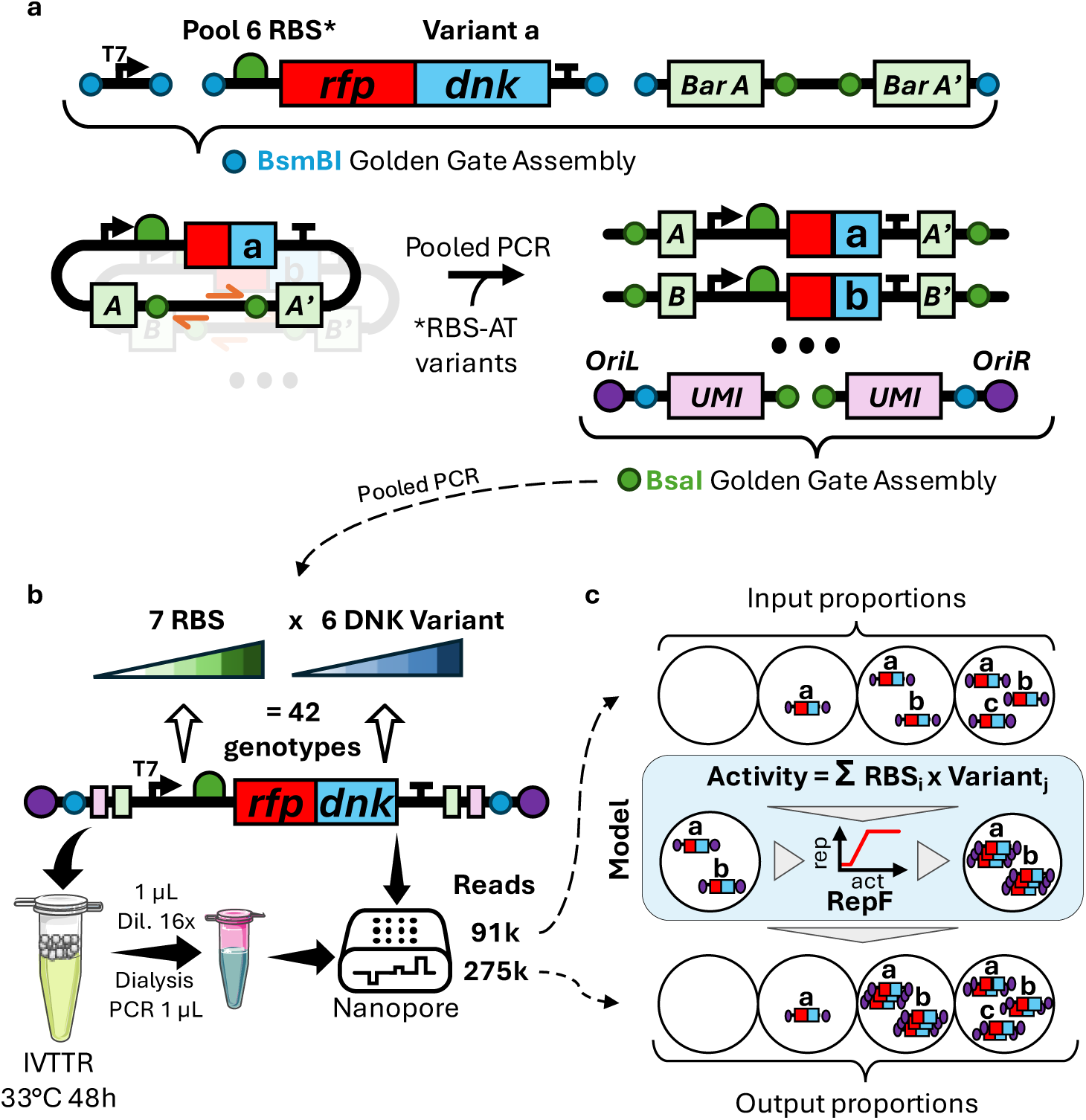
Tracking selection dynamics in a population of phenotipically diverse replicators. **a.** Construction of the combinatorial mini library by hierarchical Golden Gate Assembly (GGA). Each variant is assembled with an equal concentration of all RBS (except RBS-AT) and assigned a unique pair of barcodes. Golden Gate assembly products are pooled and PCR amplified. A second GGA is performed with another enzyme to flank every construct with a unique UMI and origins of replication. A final PCR is performed with 5’-phosphorylated primers to amplify the full construct. A BsmBI cutting site is present outside the UMI to allow addition of UMI by alternating BsaI and BsmBI GGA. **b**. IVTTR selection of the library is performed in water-in-oil emulsion. After incubation, 1 μL of emulsion is extracted with 15 μL water, dialyzed and 1 μL is PCR amplified, yielding the output libary. Input and output libraries were sequenced using nanopore. **c**. The read frequencies are then fitted by a mathematical model taking into account coencapsulation and the replication function (RepF) linking activity and replication.

The library comprised 42 constructs generated by combining six *dnk* variants harboring seven ribosome binding sites (RBS) of varied translation strength, and five variants of *dnk* corresponding to point mutations (R172I, D16N, T17S, K131E, and W150F) with a reported dTMP-to-dTDP conversion efficiency of 0, 16, 68, 86, and 117 % of the wild type, respectively (Mikoulinskaia et al., 2002).

RBS elements included five sequences named RBS-1 to RBS-5, with a wide span of predicted translation initiation rates (168, 3237, 37137, 63222 and 72085 arbitrary units, respectively) according to the RBS Calculator (Reis & Salis, 2020). Two additional RBS sequences commonly used in expression plasmids were also included: RBS-AT and RBS-C, which differ by a single point mutation and have predicted activities of 3309 and 3295 A.U., respectively. It must be noted that the previous experiment with wild-type <*rfp-dnk*> replicator was conducted with RBS-AT.

Each DNK variant was fused to mCherry in a <*rfp-dnk*> replicator, so that replication could still be monitored by RFP fluorescence. The library was emulsified in IVTTR lacking dTTP, with a replicator Poisson parameter *λ* of 1.3. We recovered DNA from the emulsion after incubation and analyzed the genotype frequencies by long-read sequencing. The input library was independently sequenced to obtain the original fraction of each genotype.

Input library composition is balanced, with a median genotype frequency of 2.3 ± 0.7 %, close to the expected 2.4% (1/42^nd^ of total), with minimum and maximum values of 1.1% and 3.9%, respectively (Figure 5a). RBS-3 is the least represented, around 9% of total and RBS-AT the most represented, around 18% of total. The output library composition is altered with median, minimum, and maximum values of 1.5 %, 0.2 % and 10 %, respectively (Figure 5b). Among all RBS backgrounds, RBS-2 is the least abundant and RBS-AT the most enriched with final proportions of 3% and 31%, respectively. These results show that replicators containing the least active RBS (e.g. RBS1 and RBS2) and/or the least active enzyme mutants (e.g. R172I) have been purified out, reflecting an efficient selection process.

**Figure 5.**
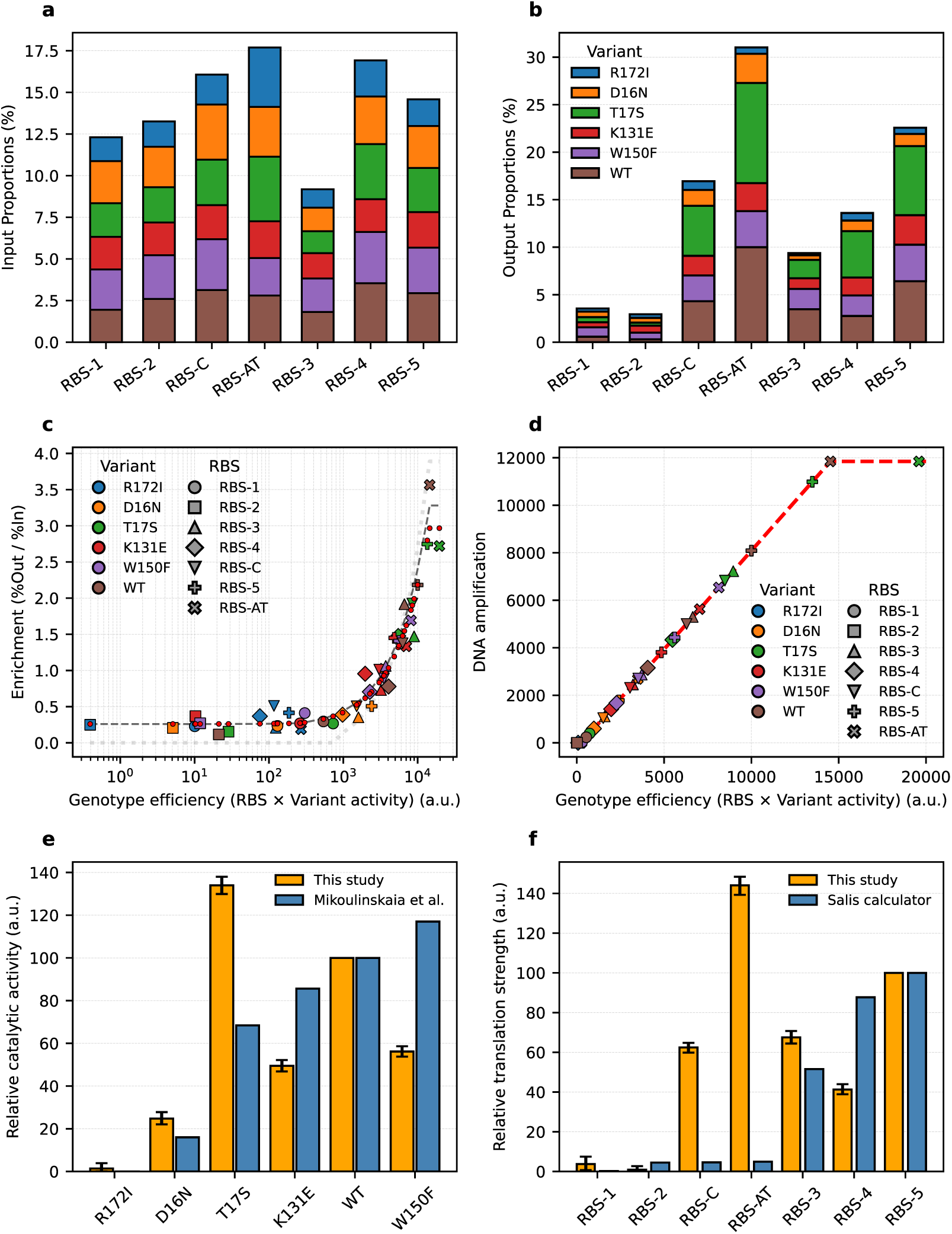
Selection dynamics of replicators with varying expression levels and DNK activity. **a, b.** Proportions of each variant and RBS in input (a) and output (b) libraries comprising variations in the RBS and DNK coding sequence. **c**. Observed and fitted enrichment value, according to the performance (RBS strength times DNK specific activity). Symbols show observed values, with color coding for DNK mutant and shape coding for RBS. The red dots are the fitted values for each variant situated on the same vertical line. Black dashed curve represents theoretical enrichment value according to the model in the hypothetical case of a flat distribution of genotypes in the input library. Dotted line represents enrichment without co-encapsulation. **d**. Replication function (RepF) representing amplification fold according to genotype activity. Genotypes are represented on the replication function according to their fitted efficiency. Genotypes with lowest replication value are superimposed at the value of 1 (no replication, only parental DNA) and RBS-AT_T17S is the best genotype with maximum replication values. **e**. Fitted parameters for variant activities, compared against reported activities in bulk measurements. **f**. Fitted parameters for RBS translational strength, compared against calculated activities using RBS calculator. Error bars indicate standard deviation computed by bootstrapping, shuffling the input and output library compositions according to the sequencing counts.

### Analysis and modelling of the mini library selection dynamics

We sought to explain the observed output library composition with a simple model, assuming that the genotype’s DNK catalytic activity and its RBS-driven expression level are independent factors and thus contribute multiplicatively to its performance (dTTP production rate). It is a reasonable hypothesis since the two processes of enzyme function and protein synthesis are distinct and the DNA concentration is very low. With this model we assume that exchanging one RBS for another should change the selective advantage of every DNK variant by the same factor. Similarly, a particularly active DNK mutation should increase the activity for all RBSs by the same factor.

Under this hypothesis, the 42 genotype data points can be described by just six variant-specific factors and seven RBS-specific factors (plus a few global parameters discussed below).

It is to be noted that kinase activity in the droplet increases as the gene is expressed, meaning the concentration of dNTP does not increase linearly. The IVTTR dynamics are quite complex: the overall protein synthesis rate declines over time in PURE system (Supplementary Figure 2), the initiation of replication of a single replicator molecule by P2/P3 is a stochastic event, and φ29 DNAP requires relatively high dNTP levels (31 μM) for optimal speed (Morin et al., 2015). Therefore, rather than attempting to derive a detailed kinetic model, we assume that the number of replicated DNA molecules is a monotonically increasing function of the activity. For simplicity, we use a linear piecewise function, RepF, which is set to 1 below a first activity threshold (only the parent DNA is carried over) and saturates at MaxRep above a second activity threshold and is thus fully specified with only 3 parameters. Note that here we average over many identical molecules of each type, which is expected to cancel the stochastic noise, and we neglect the possible effect of noncoding segments of the replicator, such as unique molecular identifiers (UMI).

We also included co-encapsulation effects in the model. Given our experimental λ (∼1.2–1.3), a minority of droplets contains more than one replicator molecule (e.g. ∼22% doubles, ∼9% triples, ∼3% quadruples by Poisson statistics). We explicitly modelled outcomes for droplets with up to three different genotypes (higher-order encapsulations were neglected). In the model, if multiple genotypes share a droplet, we assume their total dTTP production are additive, essentially treating their enzyme activities as cumulative, and sharing the corresponding replication budget (Zadorin & Rondelez, 2019). This assumes DNA concentration is low enough in the yield-determining phase so that the two genotypes do not appreciably compete for translation resources.

After fitting, the predicted and observed compositions of the output library are in good agreement (Supplementary Figure 3a). The fit allows us to extract the local fitness function, which relates each replicator to its specific amplification given the initial population distribution and the lambda parameter (Figure 5c), the replication function RepF (Figure 5d), and the variant activities and RBS strengths (Figure 5e and f). In Figure 5c, we also used the model to predict expected enrichment ratios for two hypothetical conditions: a model where all genotype frequencies were exactly 1/42 (dashed line) and a model without any co-encapsulation (dotted line).

The model-fit effective λ was 0.7, notably lower than the nominal 1.2 from initial droplet loading calculations, but higher than the 1/16^th^ value observed in previous experiment. In practical terms, a λ value of 0.7 corresponds to ∼15% of droplets with more than one replicator. This implies a non-negligible number of possible hitch-hiking replicators and thus justify considering co-encapsulation in our model.

Knowing all the genotypes efficiencies and lambda parameter, we calculated the Selection Quality Index (SQI, see material and methods) (Dramé-Maigné et al., 2020). It is a value from zero to one, one being the best selection attainable by a selection method for a randomly encapsulated library, and corrected for the initial fraction. Here we found a SQI of 0.92, a high value compared to other reported works (Dramé-Maigné et al., 2020). The difference between the relatively small observed enrichment values (up to 3.5, Figure 5c) and the good selection quality index is related to the high proportion of co-encapsulation, together with the high fraction of active variants in the input library.

### RBS and variant parameters and replication function

In Figure 5d, all the R271I genotypes (dead variant), as well as variants with RBS-1 or RBS-2, are superimposed at the bottom left of the replication function. The presence of replicated copies of these genotypes in the selected population is thus entirely attributed to hitch-hiking on the genotypes of high performance. The non-zero ramp is populated by several different combinations of variants and RBS. We classified the genotypes as strong (RBS-AT_WT, RBS-5_T17S and RBS-AT_T17S), intermediate (efficiency between RBS-1_T17S and RBS-AT_WT excluded) and inactive (efficiency below or equal to RBS-1_T17S included).

We observed a positive correlation between variant and RBS parameters and to their theoretical values (Spearman rank correlation of 0.6 and 0.8, respectively). We excluded the classic RBS-AT and RBS-C from this calculation, since they were not generated by the Salis calculator and appear to be poorly predicted. For variant strengths, the model also qualitatively confirms the activity profile of the variants, with R172I being completely inactive, D16N only displaying residual levels of activity and the four others being closer to wildtype. The observed differences may be due to the different assay conditions used in the previous study.

### Tracking single replicator fates via UMI analysis

Each DNA strand of the library carries a random stretch of 11 bases (UMI) next to each origin of replication, giving 10^12^ possible different combinations by concatenating the two UMI sequences. This number is much larger than the theoretical number of droplets collected for the analysis (3.5×10^5^ droplets, Supplementary Figure 4a). UMI clustering was performed inside each population of variants by grouping fused pairs of UMI of Levenshtein distance below 6, giving less than 0.01% of false clustering (Supplementary Figure 5) but an unknown amount of rejection of sequences in true clusters. For this reason, clusters of size 1 are artificially inflated and were excluded from analysis.

The distribution of clusters size is shown in the histogram Figure 6a, for all, strong-only or intermediate-only genotypes. Size distribution of UMI clusters across all genotypes (green) shows a bimodal population, separated by a gap around 40 counts (black dotted vertical line). A cluster of size 40 corresponds to an amplification fold of 5200 (Supplementary note 3). The median size of clusters of above 40 for all, intermediate and strong genotypes is 150, 146 and 190, respectively, giving a replication fold of 2.6×10^4^, 2.6×10^4^ and 3.3×10^4^, respectively.

**Figure 6.**
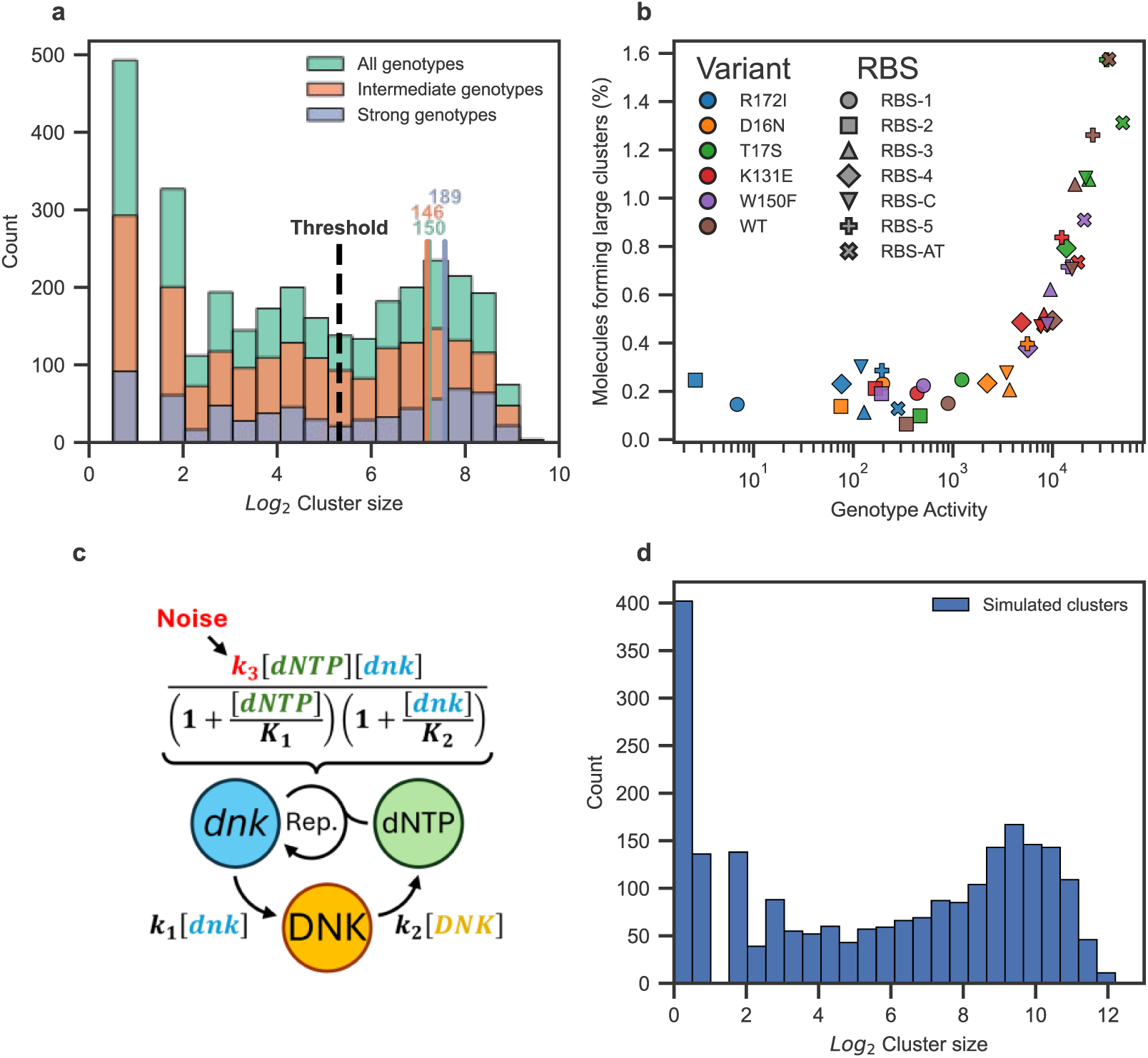
Fate of single replicator molecules. **a.** Histogram of distribution of cluster size across all genotypes (green), strongest (RBS-AT_WT and RBS-AT_T17S, blue) or intermediate genotypes (efficiency between RBS-1_T17S and RBS-AT_WT, orange). Colored vertical bar indicate median cluster size above the threshold. **b**. Relation between the number of clusters larger than 40 divided by the input proportion of each genotype in function of calculated genotype performance. A similar trend than using whole counts (Figure 5c) is observed. **c**. Scheme of the minimal kinetic model to explain observed sizes distribution. Normally distributed noise was applied to factor *k3* prior simulation. **d**. Cluster size distribution of the minimal kinetic model, reproducing the bimodal distribution of observed data.

We counted the number of clusters larger than the threshold value of 40 for each genotype and normalized this number by the number of starting DNA molecules per genotype (Figure 6b). Firstly, the maximum value of 1.6% shows that even for an active genotype, only a small fraction of DNA lead to formation of a large cluster. Secondly, we observe a similar curve as in Figure 5c, suggesting that the overall replication efficiency of a given genotype is driven by its ability to stochastically trigger replication from a single encapsulated molecule, rather than by the total number of replicators produced by such events. In other words, in compartmentalized conditions, metabolic efficiency controls initiation, rather than replication yield.

Still, the proportion of clusters under the 40 threshold is not negligible and cannot be explained only by non-replicated parental DNA molecules. Therefore, there seems to exist an intermediate regime, where replication does happen but remains limited and cannot reach the final growth stage leading to the saturated plateau. To better understand this observation, we built a minimal kinetic model of metabolic self-replication (Figure 6c). The model only assumes first order production of DNK from the gene, subsequent first order dNTP accumulation, and a gene replication process depending on dNTP concentration, with saturated kinetics. Integration of this model typically produced gene accumulation curve showing a long lag phase before entering an explosive growth regime (see Supplementary Figure 6). We then simulated droplet-to-droplet variability by drawing the replication rate *k_3_* from a normal distribution with 25% variation, all other parameters being kept constant. Multiple integration could reproduce the bimodal shape in the distribution of end-point DNA amount (Figure 6d). This supports the view that variation in individual replicator fate is due to small sources of noise, acting on very nonlinear kinetics with a long phase of slow product accumulation, and allowing them to reach, or not, the replication plateau before the replication potential of the system dies out (see Supplementary note 1 and Supplementary Figure 6). The decrease of the protein production and replication power of IVTTR system over time due to the burden of *p2p3* gene expression imposes a limit on the time window in which the dNTP accumulated may lead to replication, resulting in higher variability (see Supplementary note 2 and Supplementary Figure 15).

### Mutational scan of the DNK enzyme

*In vivo* complementation assay, where an exogeneous gene is necessary to restore the growth of an organism (e.g., a knockout placed in selective conditions), provide a convenient way to score large number of genetic variants and, in combination with deep sequencing, establish detailed mutational landscapes of a given protein (Fowler et al., 2014). Here we wished to test if our fully *in vitro* Darwinian compartment could also be used to extract the deep mutational scans of its essential enzymes.

To this purpose, we chose an RBS in which the WT variant was at the middle of the replication function: RBS-C (brown inverted triangle in Figure 5d), and we attached it to a randomized library of the <*dnk*> replicator. Here, we did not fuse the *rfp* gene, and the library was obtained by error-prone PCR, with an average of around 4 mutations per gene. Note that the introduction of 4 random mutations per gene is expected to obliterate the catalytic activity of a large fraction of the variants (Chen et al., 2023; Lundin et al., 2018; Sarkisyan et al., 2016). A bottleneck targeting 2×10^5^ molecules was applied during PCR to limit the diversity of the input library and allow for the determination of frequencies by Illumina sequencing (Figure 7a). This step also converted a UMI into a variant barcode, which was then used to facilitate tracking and sequencing This naive library was submitted to 3 rounds of selection by subjecting it to cycles of encapsulation, incubation, and PCR recovery. Each round was performed at a theoretical lambda of 0.5 in 2 pL droplets. Ten microliter of emulsion was used, giving theoretically about 3×10^6^ DNA molecules per round. No additional mutagenesis was performed in between rounds. We then used deep sequencing to track both the molecular diversity and the evolution of the mutant frequencies. To obtain both positive and negative selection signals, we also sequenced the initial library (H. Song et al., 2021).

**Figure 7.**
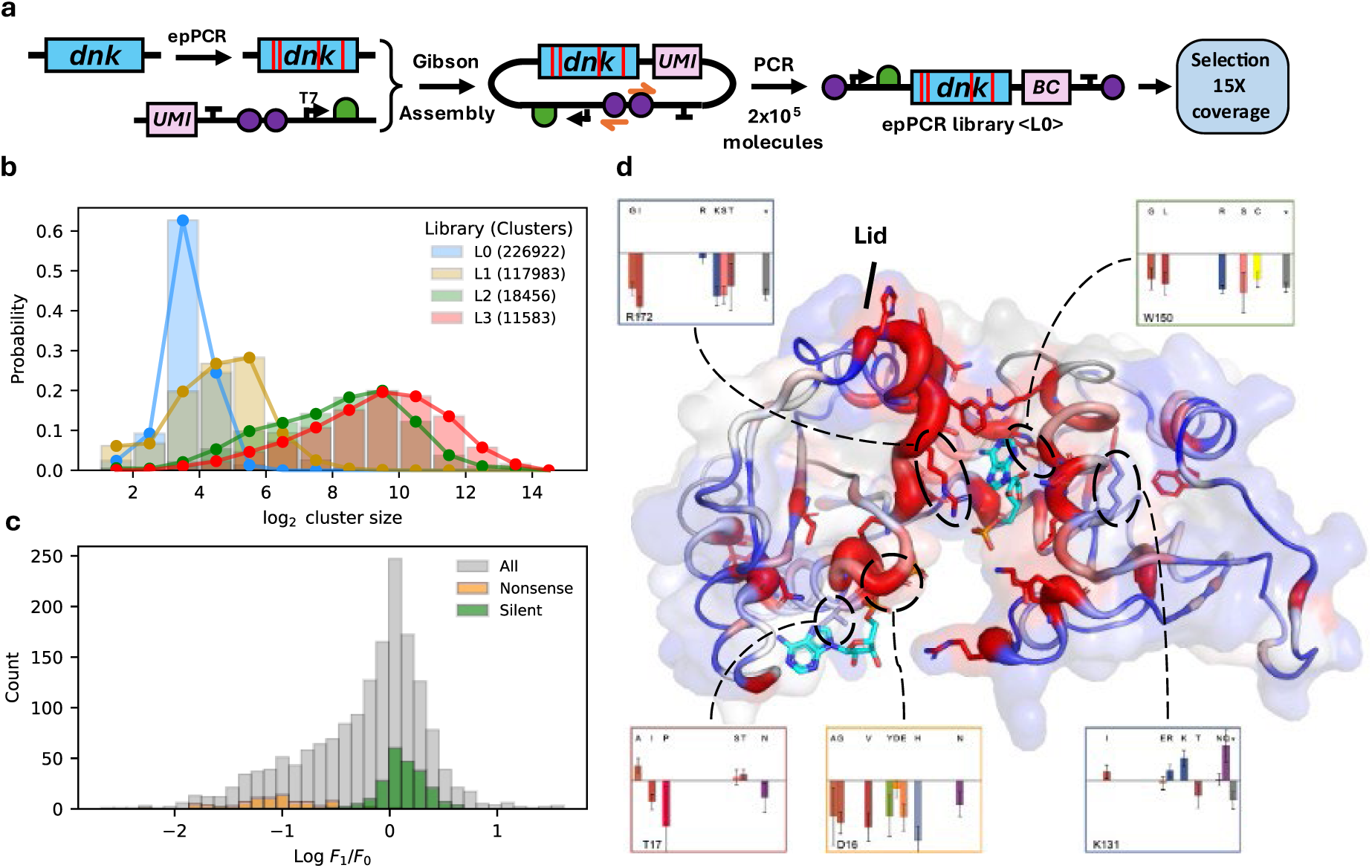
DNK mutational landscape. **a.** Creation of barcoded <DNK> replicator library using error prone PCR and Gibson assembly, restricting diversity to 2×10^5^ at the PCR step. UMI becomes barcode (BC) after PCR. **b**. Distribution of variant counts in the naïve library L0 and three successive rounds of selection L1, L2 and L3. **c**. Distribution of log-jump in frequency of individual mutation, grouped by type, between L0 and L1. **d**. AlphaFold3 model of T5 DNK including AMP and ATP with open lid, in PyMOL putty cartoon representation with tube radius and color change according to the relative entropy of the residue (blue: 0; red: 0.02). The model aligns well with the crystal structure of T4 DNK in complex with dGMP and AMP (PDB 1DEL), the AMP in the model taking the place of the dGMP acceptor of the crystal structure. Observed fitness contributions extracted with the randomized library are shown for the residues targeted in the mini library.

The UMI diversity clearly decreased during rounds of selection, with most reads partitioning into gradually less numerous but larger clusters (Figure 7b). The number of different UMI decreased by 49%, 92% and 95% after 1, 2 or 3 rounds, respectively, indicating an efficient selection. We then computed the frequency of each mutation in libraries L0 and L1, as well as the fitness contribution, computed as population-averaged log-jump in frequency, Log[*f_1_/f_0_*] (Figure 7c,d and Supplementary Figure 8-9). The total frequency decreased from 4.2 to 3.5 mutations per gene, with nonsense mutations typically rejected and silent mutations experiencing little selection pressure. Other amino-acid replacements had diverse fitness contributions, generally biased toward the negative side.To evaluate the position-wise sensitivity to mutations, we also computed the relative entropy between frequencies in L0 and L1. A high value for this metric indicates a position where the mutational spectrum strongly changes upon selection, whereas a low value indicates an insensitive position, where mutations are mostly neutrally carried over (Romero et al., 2015).

Reported on the Alphafold3-predicted structure (in presence of AMP, ATP and no magnesium ion), this analysis yields a functionally coherent pattern, where high sensitivity is associated with residues lining the active site and mutational tolerance is associated to solvent-exposed residues. One special case is the lid residues H208 and S210 which are solvent-exposed yet very conserved. By adding magnesium ion for Alphafold3 prediction we could observe that these residues cover the acceptor molecule in the active site (Supplementary Figure 7). Indeed, closing of the lid domain is required to shield the active site from solvent (Schulz et al., 1990). The individual mutational contributions obtained for the positions that were mutated in the mini library are plotted below the structure and are consistent with the previously characterized mutations and the observations in Fig. 5e. Overall, this single experiment allowed us to draw a mutational map of the DNK enzyme for 1660 mutations, covering 35% of all possible amino acid replacements (see Supplementary Figure 9 to 11).

## Discussion

In this study we successfully linked the replication of a DNA to the activity of the metabolic enzyme it encodes, forming a basic Darwinian entity in a cell free system. Upon stochastic compartmentalization in microdroplets, this fully *in vitro* system autonomously selects DNA molecules encoding active complementing phenotypes. We demonstrated the concept using a deoxynucleotide kinase gene, whose encoded protein participates in the production of the nucleoside triphosphates necessary for its replication. This system is in principle applicable to other enzymes, given that a link can be established with DNA replication. An important point is that the metabolic positive feedback should be able to ignite from the very low initial DNA concentration corresponding to one (or a discrete number of) DNA molecules in the compartment. In our case, it appeared that the molecular amplification stages involved in cascaded transcription/translation of DNK activity were strong enough to trigger replication from picomolar initial replicator concentrations, compatible with picolitre compartments. Smaller compartments may be required for designs with lower sensitivity.

Early molecular replicators may have emerged independently of the defined boundaries of a cell, subjected to cycles of mixing and transient compartmentalization (Blokhuis et al., 2020; Matsumura et al., 2016; Szathmáry & Demeter, 1987; Szathmáry & Smith, 1995). Compartmentalization may have provided a linkage between genotype and phenotype, but also limited the accumulation of fast replicating parasitic species. This scenario blurs the definition of a genome, as it usually refers to the genetic material of an organism because here, the organism forms when encapsulation occurs. Thus, the “genome” of a compartment can be considered as the ensemble (one or more) of replicators that have been co-encapsulated. Nevertheless, the evolving entities are the individual replicators, interacting with each other similarly to Wilson’s trait-group model (Wilson, 1975).

A fluorescent gene inserted in the replicator genome allowed to visually track the replication of single replicators in droplets. Even in the case of a clonal population of replicators, we observed a large droplet-to-droplet variation of RFP fluorescence and a lower-than-expected fraction of active droplets (about 1/16^th^). These findings were confirmed by individual replicator tracking using UMI and deep sequencing, where for strong genotypes, only about 1.6% of molecules were able to form large clusters. These observations are reminiscent of previous findings that showed increases in variation for replication-expression systems, compared to PURE-only experiment (Abil et al., 2022), although this reported observation was made in much smaller compartments.

Compartmentalized self-replication systems autonomously select enzymes with particular activities from large genetic libraries. However, it is generally difficult to precisely characterize the selection dynamics at play in these approaches. Indeed, most artificial selection systems are assayed using a simple binary library. A known mixture of wild-type enzyme and inactive variant is created, passed through the process, and the induced change in active/inactive ratio is used as a proxy for the purifying efficiency of the system. Here, by using a barcoded mini library containing a wide range of enzymatic activity and expression levels, we managed to characterize more precisely the selection process.

Firstly, it allowed us to characterize the effect of co-encapsulation on the survival of inactive genotypes. This phenomenon can be understood as a form of hitch-hiking permitted by random partitioning of active species. Similar hitch-hiking effect has already been observed by Ichihashi and colleagues (Furubayashi et al., 2020; Ichihashi et al., 2013; Mizuuchi & Ichihashi, 2018). In our study, parasitic genomes did not appear spontaneously, neither did they obtain a replicative advantage by reducing their genome size.

Secondly, the mini library experiment gave us more insight into the inner parameters of the system, especially the shape of its replication function. Here, the data was accurately fit with a simple piecewise linear relationship between activity and replication. This smooth behavior at the population level needs to be reconciliated with a large variability at the droplet, hence individual replicator levels. Our initial hypothesis was that the probability for replication to initiate is low. This could be due to multiple nonexclusive mechanisms. First, P2 and P3 are produced *in situ*, implying a low concentration in the early phase, and they need to associate prior binding to DNA ends in order to initiate replication. Single encapsulated DNA target concentration is only 1.5 pM, limiting the chances of association. Second, it has also been described that the phosphorylated origins of replication, which are used here, are suboptimal compared to TP-capped template (Mencía et al., 2011; Van Nies et al., 2018).

Deep sequencing of the paired UMIs equipping each replicator molecule in the mini library, provided a finer tracking of their individual fate. When grouped by UMI copy number, this revealed two sizes of clusters, regardless of the genotype performance, indicating two possible fates for a given genotype. We have shown that this is compatible with a simple kinetic model of self-sustained IVTTR. According to our model, the fact that DNA replication depends on the catalytic activity of a product of the replicating DNA itself induces a lag phase followed by explosive growth. This kinetic regime appears very sensitive to small fluctuations and may explain the variable fate of replicators with identical payload. Importantly, the model is able to reproduce the bimodal distribution of UMI copy number.

Overall, our compartmentalized self-replication scheme does efficiently select active *dnk* genes over inactive ones, but not all identical replicators experience the same outcome. Our observations do not support the all-or-nothing replication scenario, but rather more subtle scenario where many functional replicators only undergo a background level of DNK-dependent amplification. Several factors may contribute to droplet-to-droplet heterogeneity in replication efficiency and future work will be needed to pinpoint the dominant causes and decrease fate variability.

Beyond fundamental insights into self-selection processes, this *in vitro* minimal Darwinian compartment provides a scalable platform for directed evolution of enzymatic functions, circumventing limitations faced with living systems, including cytotoxicity and transformation inefficiencies (Dramé-Maigné et al., 2024). The high-throughput and quantitative nature of our technology provides a valuable tool for rapid protein engineering, as demonstrated with the establishment of a mutational map of DNK, submitted to dTMP phosphorylation pressure. It would be interesting for example to map the response to other nucleotide replacements, to better understand the basis for nucleotide promiscuity in this family of enzymes. We have also shown that multiple selection rounds can be performed with minimal efforts, opening the way to challenging directed evolution campaigns. In addition, the high efficiency of the φ29 replication machinery enables the amplification of long replicators containing multiples genes (Restrepo Sierra et al., 2025). As such this tool may prove valuable to study the build-up of complexity in minimal evolutionary systems. For example, by inserting 2 copies of the *dnk* gene on a same replicator, we could track the evolution of genes following duplication, and monitor the occurrence of inactivation or specialization (Ohno, 1970).

## Material and Methods

### Reagents

Reagent suppliers and their products were as follows. GeneFrontier: PUREfrex 1.0, 2.0 and 2.1. Thermo Fisher: Dextran Alexa647 10000 Da Anionic (ref. D22914) and Dextran Alexa Fluor 680 3000 Da Anionic (ref. D34681), diluted in milliQ water and stored at -20 °C. Biotium: EvaGreen Dye 20X (ref. 31000). Sigma: dGMP, dTMP, dCMP and dAMP, references D9500-100MG, T7004-100MG, D7625-100MG, and D6375-100MG respectively, diluted in milliQ water and stored at -20 °C. MF-Millipore: Membrane Filter, 0.025 µm pore (ref. VSWP02500).

### Molecular cloning

All plasmids used in this study are listed in Supplementary Table 3. Plasmid pUC_OriLR_p2p3 was given by Dr Christophe Danelon. Plasmids pAcc-pXfus-dnk and pUC_OriLR_gfp were obtained from pUC_OriLR_p2p3 using standard molecular biology techniques and the sequence verified using Sanger sequencing. Plasmids pEX-A128-pAnobar pEX-A125-pBnobar containing φ29 origins of replication sequences OriL and OriR in an inverted orientation were ordered as clonal plasmid from Eurofins Genomics. X_T7promoter was ordered as a dsDNA GeneStrand (Eurofins). Polymerase Chain Reactions (PCR) were conducted using One Taq Master Mix (ref. M0484S) or Q5 Hot Start High Fidelity Polymerase (ref. M0493S, used in High Fidelity buffer) from New England Biolabs, with dNTP and primer concentrations as indicated by the manufacturer. Other enzymatic reagents were obtained from New England Biolabs: DpnI (ref. R0176S), NEBuilder Master Mix (ref. E2621S), Exonuclease V (ref. M0345S), Blunt/TA Ligase Master Mix (ref. M0367S), Golden Gate Assembly (GGA) Kits (BsaI-HFv2, ref. E1601S and BsmBI-v2, ref. E1602S). RNAse A (ref. EN0531) was obtained from Thermo Fisher and RNAse One (ref. M4261) from Promega. All bacterial transformations were done using *E.coli* NEB 5-alpha competent cells High Efficiency from New England Biolabs (ref. C2987H). Luria Broth (LB) cultures or agar plate were supplemented with ampicillin at a final concentration of 100 μg/mL.

### Purification of SSB DSB proteins

Single-stranded DNA binding protein and DSB were expressed and purified as described in (Soengas et al., 1995) and (Mencía et al., 2011), respectively.

### dnk point-mutation plasmids

pAcc_pXFus_dnk containing the dnk wildtype sequence from T5 fused in C-terminal of mCherry by a flexible linker (GGGGS LVPRGS GGGGS) under the control of RBS-AT was used as a starting point to introduce point mutations in the dnk sequence (Mikoulinskaia et al., 2013). 1.5 amol/μL of plasmid was amplified by PCR using Q5 polymerase using primers T242/T243, T244/T245, T246/T247, T248/T249 and T250/251 for mutations R172I, D16N, T17S, K131E, W150F respectively using program **PCR 5** with the following annealing temperatures: 64 °C for R172I, D16N, T17S and W150F; 59 °C for K131E. Crude PCR product was then circularized using 1/10^th^ of PCR in KLD mix and incubated 10 min at 25 °C. NEB 5-alpha competent cells were transformed with crude KLD product (7% of total volume) following a quick version of manufacturer’s protocol, spread on LB agar with ampicillin and grown overnight at 37 °C. Individual colonies were picked and grown overnight in 4 mL of LB with ampicillin, and plasmid was extracted using Miniprep kit (Macherey-Nagel, ref. 740588.250) following manufacturer’s protocol, eluting in 50 µL MilliQ water. Purified plasmids were Sanger sequenced using primers iR120, iR184 and iR186 to confirm desired sequences.

### GFP reporter experiment

A typical IVTTR reaction mastermix was assembled with PUREfrex 1.0 (Supplementary Figure 2) or PUREfrex 2.0 (Figure 2) solutions diluted as in manufacturer protocol (GeneFrontier), 20 mM Ammonium sulfate, 0.375 mg/mL P5, 0.105 mg/mL P6, 100 nM of plasmid pUC_OriLR_p2p3, 25 nM of wild type sequence of dnk gene gblock (IDT) and 12 pM of the replicator <*gfp*>. 4 μL of mastermix was aliquoted in an optical PCR microtube (white, low profile, BioRad) and 1 μL of dNTP mixture of different composition was added to a final concentration of 0.3 mM. Reactions were incubated in real-time thermocycler (CFX Touch BioRad) at 33 °C for 20 h while monitoring fluorescence in all channels (examples in Supplementary Figure 1).

### qPCR quantification

IVTTR samples were diluted to 1/1000^th^ prior to qPCR quantification in order to avoid PCR inhibition by PURE components. A standard 10 μL reaction contained OneTaq HotStart 2X MasterMix to 1X final concentration with 3% (v/v) Evagreen 20X (Biotium), 200 nM of each primer and 2 μL of diluted DNA sample. Primers were T133/T134 for dnk and T129/T130 for *gfp*. 10-fold serial dilutions of a standard DNA from 10^-10^ M to 10^-16^ M of were used for precise quantification. Standard DNA were gBlock T7_RBS-AT_dnk_wt for dnk quantification, its concentration determined by manufacturer and pUC_OriLR_gfp miniprep for *gfp* quantification, its concentration determined by UV spectrometer (Clariostar, BMG labtech).

### <*rfp-dnk*> selection

The selection of <*rfp-dnk*> wild type was conducted by assembling a typical IVTTR reaction with PUREfrex 2.0 solution (GeneFrontier), 20 mM Ammonium sulfate, 0.375 mg/mL P5, 0.105 mg/mL P6, 200 nM accessory plasmid pUC_OriLR_p2p3 and the replicator to the final concentration of 1 pM yielding a theoretical lambda of 1.5 in 2.5 pL droplets. The solution was emulsified in HFE Novec 7500 fluorinated oil (3M) supplemented with 4% Fluosurf-C (Emulseo) and filtrated, using a Millipede device with oil and aqueous phases pushed at approximately 200 mbar pressure using Microfluidic Flow Control System (Fluigent) through PTFE tubing. The droplets were gathered in a 1 mL pipet tip at the outlet of the millipede device and collected in multiple PCR tubes for incubation in thermocycler overnight at 33 °C. Emulsion was imaged using Nikon Ti fluorescence microscope (see below). The emulsion layer was transferred to fresh HFE Novec 7500 and supplemented with 20% (v/v) perfluorooctanol aided by pulses using Zerostat 3 Anti-Static Gun (Milty) until a clear drop was observed. The solution was flicked and spun down on a tabletop centrifuge and 2 µL of supernatant was cascade-diluted up to 10000-fold in milliQ water prior to qPCR quantification.

### Library design strategy

We designed 5 RBS of different translation activities using Salis lab RBS calculator in control mode (Cetnar & Salis, 2021; Reis & Salis, 2020; Salis et al., 2009). We also included two variant of the commonly used RBS of T7 phage gene 10 (Olins and Rangwala, 1989) in the context between RBS and start codon: ”tatacAT” found in pET family plasmids, and “tatacC” found in cell free expression plasmid pIVEX2.4d for example, these variations will be referred as RBS-AT and RBS-C, which translation rate were predicted as only 3558 and 3309, respectively. The library was assembled in a semi-batch fashion using hierarchical Golden Gate assembly (Figure 4a). Each variant construct was flanked by a unique pair of 8-bp DNA barcodes, allowing unambiguous genotype identification via nanopore sequencing. The use of paired barcodes and nanopore long-read sequencing also enabled the detection of chimeric sequences arising during PCR amplification. Such sequences were then removed from analysis. Following ligation of different barcodes and RBS, all constructs were pooled and flanked by φ29-compatible replication origins (OriL and OriR). Both replication origins contained unique molecular identifiers of (UMI) to enable single molecule tracking of amplification dynamics and bias. Parts were obtained as follows (all sequences are in Supplementary Table 3)

### Barcoded X_OriLR1-12 parts

Plasmid pEX-A128-pAnobar (0.12 amol/μL) was amplified using Q5 polymerase and primers M76/M77 using program **PCR 1** (table below). After amplification, PCR products were purified (Zymo Research), eluted with milliQ water. The obtained fragment of 619 bp formed a single band on agarose gel. The fragment was then PCR amplified with 12 different pairs of primers bearing different barcodes. Pairs of primer are as follows: Bar1: M111/M112; Bar2: M113/M114; Bar3: M115/M116; Bar4: M117/M118; Bar5: M119/M120; Bar6: M121/M122; Bar7: M123/M124; Bar8: M125/M126 Bar9: M127/M128; Bar10: M129/M130; Bar11: M131/M132; Bar12: M133/M134. All PCR parameters were the same except for primers used: 0.24 amol/μL of plasmid, 1 unit of Q5 polymerase. Samples were incubated in a thermocycler using program **PCR 2.** After amplification, PCR products were purified (Zymo Research), eluted with 25 µL milliQ water, yielding 12 barcoded OriLR parts.

### UMI-bearing Y_OriLumi and Y_OriRumi parts

Plasmid pEX-A125-Bnobar (0.12 amol/μL) was PCR amplified using Q5 polymerase with primers M173 and M174 bearing stretch of degenerated bases NNWNNNNNWNN. Samples were incubated in thermocycler using program **PCR 3** After amplification, PCR products were purified (Zymo Research), eluted with 20 µL milliQ water, yielding a product called YoriLRumi. A second PCR was performed on this product using outer primers. OriL and OriR were amplified separately from the same initial product using primers T36/M190 and T37/M191, respectively, to avoid correlation bias between left and right UMI. It should be noted that barcode diversity being 1.10^6^, thus 100 amol represents multiple times (∼60X) the diversity. In a PCR microtube were mixed around 100 amol of YoriLRumi, 2 units of Q5 polymerase, 0.2 mM of each dNTP, 1% of Evagreen 20X (Biotium) and 500 nM forward and reverse primers in a total volume of 100 µL. Samples were incubated in a thermocycler using program **PCR 4** and the product was purified (Zymo Research) and eluted with 25 µL MilliQ water. DNA concentration was determined using Clariostar plate reader (BMG labtech) yielding 95 nM and 99 nM for Y_OriLumi and Y_OriRumi respectively.

### X_RBS_mCherry_dnk_T7term parts

42 different RBS_mCherry_dnk_T7term parts representing a combination of seven RBS and six variants were obtained by PCR amplification of each mutant plasmids pAcc_pXFus_dnk, with each forward primer bearing different RBS and a common reverse primer. Forward primers are as follows: RBS-1: M139; RBS-2: M140; RBS-3: M143; RBS-4: M141; RBS-5: M142; RBS-AT: MT01; RBS-C: M137, reverse primer is M84. All PCR parameters were as follows: In a PCR microtube were mixed 0.1 fmol of plasmid, 1 unit of Q5 polymerase, 0.2 mM of each dNTP, 2% Evagreen 20X (Biotium) and 500 nM forward and reverse primers in a total volume of 50 µL. Samples were incubated in a thermocycler using program **PCR 6.**

After amplification, 10 µL of crude PCR products of RBS 1 to 5 and RBS-C of each variant were pooled and purified (Zymo Research), eluted with 15 µL milliQ water. 50 µL of PCR products of each variant of RBS-AT were purified separately. All 12 purified RBS-variant parts were quantified using clariostar and adjusted at 25 nM with milliQ water. Golden Gate Assemblies (GGA) were performed using barcoded XOriLR part 1 to 6 for RBS-pooled variants R172I, D16N, T17S, K131E, W150F and WT, respectively, and XOriLR part 7 to 12 for individually purified variants R172I, D16N, T17S, K131E, W150F of RBS-AT, respectively. Assemblies were performed as follows: In a PCR tube, 2.5 nM of XoriLR part, Xprom part and RBS-variant part were assembled to 1X ligase buffer and 1/20^th^ of BsmBI GGA mix in a total of 10 µL. Reaction mixtures were incubated in a thermocycler (ProFlex PCR System, Applied Biosystems) for 30 cycles alternating 42 °C for 1 min and 16 °C for 1 min, followed by an inactivation step of 5 min at 60 °C.

To obtain an equimolar distribution of each variant-RBS combination, 6 µL of each GGA 1 to 6 were pooled with 1 µL of each GGA 7 to 12. Assembled products were amplified by PCR using primers GR25 and GR26 that truncate both origins of replication, using the following protocol: In an optical PCR microtube (white, low profile, BioRad) were mixed 1/50^th^ of pooled GGA products, 2 units of Q5 polymerase, 0.2 mM of each dNTP, 1% Evagreen 20X (Biotium) and 500 nM of primers GR25 and GR26 in a total volume of 100 µL. Reaction mixtures were incubated in qPCR machine (Miniopticon, BioRad), using the following program: Samples were incubated in thermocycler using program **PCR 7.** The amplification product was purified (Zymo Research) and eluted in 10 µL MilliQ water. Concentration was measured using Clariostar plate reader (BMG Labtech) yielding 7.6 ng/µL of product. Analysis of 2 µL of product on 1% agarose gel showed a single band at higher mass than expected (>3000 bp instead of 1980 bp) but we assumed a gel artifact.

### Addition of UMI

Golden Gate Assembly was carried out between the purified PCR product and UMI-bearing Y_OriLumi and Y_OriRumi parts. In a PCR tube, 2.5 nM of Y_OriLumi part, Y_OriRumi part and PCR product were assembled to 1X ligase buffer and 1/20^th^ of BsmBI GGA mix in a total of 10 µL. Reaction mixtures were incubated in a thermocycler (ProFlex PCR System, Applied Biosystems) for 1 h at 37 °C, followed by an inactivation step of 5 min at 65 °C. 6 μL of crude GGA product was used for a PCR using 5’-phorphorylated primers. In an optical PCR microtube (white, low profile, BioRad) were mixed 6 μL of pooled GGA products, 6 units of Q5 polymerase, 0.2 mM of each dNTP, 1% Evagreen 20X (Biotium) and 500 nM of primers T22 and T23 in a total volume of 300. Reaction mixtures were incubated in qPCR machine (Miniopticon, BioRad), using the following program: Samples were incubated in thermocycler using program **PCR 8.** The amplification product was purified (Zymo Research) and eluted in 10 µL MilliQ water. Agarose gel analysis revealed a band at the expected size and a lower band at 1000 bp. Gel extraction was performed (Macherey-Nagel) and DNA was eluted in 50 μL milliQ water. DNA was quantified with Qubit dsDNA Quantification Assay (ThermoFisher) at 3.7 nM. A single band was seen on agarose gel.

### Mini library selection

A typical IVTTR reaction mastermix was assembled with PUREfrex 2.1 solution (GeneFrontier), omitting reducting agent, 20 mM Ammonium sulfate, 0.375 mg/mL P5, 0.105 mg/mL P6, 200 nM of plasmid pUC_OriLR_p2p3, 25 nM of wild type sequence of *dnk* gene gblock (IDT) and 1 pM of the mini library. The mastermix was split in three aliquot; in the first aliquot 0.3 mM of each dNTP and 4 mM GSH were added, in the second aliquot 0.3 mM of each dATP, dCTP, dGTP, dTMP and 2 mM DTT were added, in the third aliquot, 0.3 mM of each dATP, dCTP, dGTP, dTMP and 4 mM GSH were added (see Supplementary Figure 16). dTMP conditions contained 2 μM of Alexa Fluor 680 3000 Da Anionic was added as a marker. Sample partitioning was performed by a step-emulsification process using a one-inlet disperse phase in 2 pL droplets, yielding an expected Poisson *λ* parameter of 1.3 for the replicator. The continuous phase consisted of a fluorinated oil (HFE Novec 7500, 3 M) supplemented with 4% w/w Fluosurf-C (Emulseo). Emulsions were gathered in a 200 μL PCR microtube and incubated at 33 °C for 48 h.

### MinION sequencing and analysis of Mini library

DNA libraries were purified (Zymo Research) according to manufacturer’s protocol and dialyzed against 50 mL of milliQ water for one hour at room temperature using Membrane Filter, 0.025 µm pore (MF-Millipore VSWP02500) prior to sequencing. The sequencing was performed using 1D ligation chemistry (SQK-LSK110), sequenced in a FLO-FLG001 (R9.4.1) flow cells and basecalled using Guppy software with dna_r9.4.1_450bps_sup model, Oxford Nanopore. Basecalling gave 1.84×10^5^ and 4.90×10^5^ passed reads for input and output, respectively. After size filtering for expected length ±200 bp, sequences were aligned against theoretical sequence RBS-C_WT as reference using minimap2. In house R script was then used to analyze each CIGAR codes of highest quality alignments (56 and above) and retrieve positions and sequences of UMI, tags and RBS. The Levenshtein distance was measured against tag references and the closest was assigned if measured distance was inferior or equal to 2 with no ambiguity, otherwise the tag was marked as unassigned. Similarly, RBS were compared with references and assigned if Levenshtein distance was inferior or equal to 5 with no ambiguity, except for RBS-AT and RBS-C which were pooled together and sorted out using their unique tags. Only fully attributed reads with matching left and right barcodes were kept for analysis. Information is summarized in Supplementary Table 1.

### Barcoded acceptor vector for Gibson Assembly pGibs1

pGibs1 is a Gibson assembly acceptor DNA that bears the origin of replication OriR and OriL flanking T7 promoter and T7 terminator, also containing an UMI upstream the terminator. We first circularized the 34 ng <*gfp*> replicator using 10 μL KLD mix and incubating 10 min at room temperature. A PCR reaction using Q5 polymerase was carried out on 1 μL of KLD product, with forward primer T181 binding upstream of T7 terminator and carrying a tail containing secondary primer binding sequence and a stretch of 30 degenerated nucleotides “N” and reverse primer iR397. Samples were incubated in a thermocycler using program **PCR 9.** The reaction product was purified (Macherey-Nagel) following supplier instructions, and eluting using 50 μL of 10-fold dilute AE buffer. It is expected that the PCR product should contain 50% of mismatched barcodes. Barcodes were completed by a second PCR using Q5 polymerase and 400 pmol of the first PCR in 100 μL reaction, with primers T182 and iR176 that bind at both extremities outside of the degenerate region. Samples were incubated in a thermocycler using program **PCR 10.** The product of reaction was purified (Macherey-Nagel) following supplier instructions, and eluting using 50 μL of AE buffer diluted 1/10^th^.

### Barcoded acceptor vector for Gibson Assembly pGibs2

pGibs2 plasmid was created to accept second round of selection of error prone library. Two PCR were used, one for adding UMI using degenerated primers and a second PCR of two cycles to complement the mismatching UMI. For the first PCR, in 200 μL reaction were added 5 Units of OneTaq HotStart, 0.4 mM of dNTP mix, 0.2 μM of T274/T304, 72 μg of plasmid pUC_OriLR_gfp, 2% of Evagreen 20X. The sample was incubated in a thermocycler using program **PCR 11.** To remove parental DNA, 40 U of DpnI was added, and solution was incubated at 37 °C for 1 h. 10 μL of unpurified PCR product was used for the completion round. In 200 μL reaction were added 4 Units of Q5 polymerase, 0.2 mM of dNTP mix and 0.5 μM of primers T275/T305. The sample was incubated in a thermocycler using program **PCR 12**. A single band was observed on agarose gel, so the linear plasmid was purified (Macherey-Nagel) and eluted in 20 μL milliQ water.

### DNK random mutational libraries creation

Plasmid pUC_OriLR_dnk was amplified by PCR using Genemorph II mutagenesis kit (Qiagen) to obtain DNA library. A volume of 50 μL reaction containing 0.13 ng/μL of plasmid starting concentration was PCR amplified using Q5 Polymerase and iR251 forward primer and 5’-biotinylated reverse primer T227. Samples were incubated in a thermocycler using program **PCR 13.** Parental DNA was digested by adding 1 μL of DpnI restriction enzyme followed by 15 min incubation at 37 °C. DNA was purified (Zymo Research) and eluted with twice 10 μL elution buffer yielding 99 ng/μL (134 nM or 8.10^10^ molecules/μL) of DNA measured using ClarioStar (BMG Labtech).

A second PCR reaction of 100 μL using Q5 polymerase on 1.10^11^ DNA molecules was performed with nested primers T167/T232 to increase the DNA quantity without introducing mutation and prepare the DNA ends for Gibson Assembly. As the primers were binding on the 9 first and 7 last amino acids of the coding sequence, these amino acids are not mutated in the final library. Samples were incubated in a thermocycler using program **PCR 14**. To remove the biotinylated parental DNA, a capture step was performed using Dynabeads MyOne Streptavidin C1 (Thermo Fisher Scientific) with the following protocol. 30 μL of beads were conditioned by a wash with 500 μL NaCl at 1.25 M, then diluted again in 250 μL NaCl at 2 M. 100 μL of the bead solution was added to 100 μL of PCR reaction and incubated 15 minutes at room temperature. Beads were peletted using a magnet and the supernatant was kept. Supernatant was purified (Zymo research) and eluted with twice 10 μL of elution buffer, yielding 20 μL at 110 ng/μL (236 nM or 1.4×10^11^ molecules/μL).

The DNA fragment was then inserted into the barcoded acceptor vector pGibs1 using Gibson assembly. In 20 μL total reaction, 10^11^ molecules of vector and 2×10^11^ molecules of *dnk* library were mixed with Gibson Assembly MasterMix and incubated at 50 °C for 30 min in a thermocycler. All linear fragments were then digested by Exonuclease V (RecBCD, NEB). 20 μL of the assembly product was supplemented with NEB buffer 4, ATP to 1 mM final, 10 U Exonuclease V and milliQ water to a final volume of 50 μL. The mixture was incubated at 37 °C for 30 min and heat inactivated at 70 °C for 30 min. Considering 5% of reaction efficiency, a PCR was performed using Q5 polymerase directly on the Gibson assembly solution diluted down to 200000 starting molecules, using phosphorylated primers T22/T23 giving replicative library <L0>. Samples were incubated in a thermocycler using program **PCR 15.** The product of reaction was purified (Zymo research) following manufacturer’s instructions and eluted using twice 10 μL of elution buffer, yielding 20 μL of input library measured at 21 nM using ClarioStar.

### Random mutational library selection

A scheme indicating the workflow of random mutational library selection is shown Supplementary Figure 12. The first round of selection was conducted by assembling a typical IVTTR reaction with PUREfrex 1.0 solution (GeneFrontier), 20 mM Ammonium sulfate, 0.375 mg/mL P5, 0.105 mg/mL P6, 200 nM accessory plasmid pUC_OriLR_p2p3 and the replicator to the final concentration of 0.5 pM, yielding a theoretical lambda of 0.6 in 2 pL droplets. The solution was emulsified in HFE Novec 7500 fluorinated oil (3M) supplemented with 4% Fluosurf-C (Emulseo) and filtrated, using a Millipede device with oil and aqueous phases pushed at approximately 200 mbar pressure using Microfluidic Flow Control System (Fluigent) through PTFE tubing. The droplets were gathered in a 1 mL pipet tip at the outlet of the millipede device and collected in multiple PCR tubes for incubation in thermocycler overnight at 33 °C. The emulsion layer was transferred to fresh HFE Novec 7500 and 10 μL of emulsion (around 3×10^6^ droplets) was transferred to 90 μL ultra-pure water. Emulsion was destabilized using Zerostat 3 Anti-Static Gun (Milty) (Karbaschi et al., 2017).

The whole construction was amplified using Q5 polymerase and primers T22 and T23 in 100 μL reaction, using 0.5 μL of dilution. Samples were incubated in a thermocycler using program **PCR 16.** Reaction product was gel purified (Macherey-Nagel) as manufacturer’s protocol and eluted using twice 10 μL NE buffer diluted 1/10^th^ yielding 20 μL of purified replicative library <L1> at 35 nM measured using ClarioStar).

A second round of selection was performed directly with the amplified library <L1> using the same protocol of the first round, except pUC_OriLR_p2p3 plasmid concentration was fixed at 100 nM, emulsifying 30 μL of IVTTR solution. After selection, 10 μL of emulsion was broken and 1 μL was used to amplify the gene using primers T84/T22 and Q5 polymerase. Samples were incubated in a thermocycler using program **PCR 17.** A single band was seen on agarose gel. DNA was purified (Macherey-Nagel) and DNA was eluted in 50 μL milliQ water. To add Gibson Assembly overhangs, a second PCR was performed using primers T276/T306 and Q5 polymerase. Sample was incubated in a thermocycler using program **PCR 18**. Gibson assembly was performed using 51 ng of insert and 87 ng of acceptor plasmid pGibs2, in 20 μL reaction of NEBuilder. Gibson assembly was incubated at 50 °C for 40 min and 1 μL of the crude Gibson assembly product was used for a PCR with phosphorylated primers as follows: in 20 μL were added 0.4 Units of Q5 Polymerase, 0.2 mM dNTP mix, 0.5 μM of T22/T23 primers and 5 μL Evagreen 20X. Sample was incubated in a qPCR thermocycler (Miniopticon, BioRad) using program **PCR 19**. As a secondary band of lower molecular size was observed on gel, the PCR product was gel purified (Macherey-Nagel) and eluted with 20 μL milliQ water, giving the replicative library <L2>.

The third round of selection was performed using the same protocol of the second round, emulsifying 30 μL of IVTTR solution. After selection, 10 μL of emulsion was broken and amplified for illumina sequencing with UMI flip (below).

### Illumina sequencing libraries L0 and L1

Samples from libraries <L0> and a 10-fold dilution of broken emulsion after selection of <L0> (called L1) were amplified using Q5 polymerase and two sets of primers, T313/T314 and T315/T316 to amplify 500 base pairs fragments of N-terminal or C-terminal coding sequence, respectively. In 50 μL reaction were mixed 1 Unit of Q5 Polymerase, 0.2 mM dNTP mix, 0.5 μM of each primer and 140 pM of library <L0> or 1 μL of diluted library L1. As T314 and T315 partially overlap, mutations on amino acids Q143 to R148 were not sequenced. A 10 μL aliquot was taken and supplemented with 5% Evagreen 20X to follow the reaction in a real-time PCR machine (CFX Touch, BioRad) and determine the optimal cycle number. Samples were incubated in thermocycler using program **PCR 20** using an annealing temperature of 56 °C for T313/T314 and 69 °C for T315/T316, and a number of cycles N = 19, 19, 15, 16 for L0 Nter, L0 Cter, L1 Nter, L1 Cter, respectively. The reaction products were purified (Zymo research) following manufacturer’s instructions and eluted using 20 μL of elution buffer, yielding 20 μL of DNA quantified with ClarioStar at: L0 Nter: 117 ng/μL, L0 Cter: 115 ng/μL, L1 Nter: 103 ng/μL, L1 Cter: 115 ng/μL. Samples were visualized on agarose and a single band was observed. Samples Nter and Cter were pooled and sent for Illumina sequencing analysis using Genewiz Amplicon-EZ platform.

### Illumina sequencing with UMI flip libraries L1 L2 L3

In order to link N-terminal and C-terminal sequences using 500 base-pair sequencing, as the UMI is positioned downstream the *dnk* sequence, the libraries L1, L2 and L3 were subjected to a first PCR of the whole DNK construct, followed by a circularization by blunt ligation, followed by a second PCR of the N-terminal sequence with the barcode flipped in 5’. Crude broken emulsion was 10-fold diluted in milliQ water and 1.2 μL was used in the following PCR: in 60 μL was mixed 1.2 Units of Q5 polymerase, 0.2 mM dNTP mix, 0.5 μM of phosphorylated primers T393/T394 and 1% Evagreen 20X. Sample was incubated in a real-time thermocycler (Miniopticon, BioRad) using program **PCR 21**. Amplified DNA was purified (Zymo Research), eluted with 30 µL milliQ water. Circularization was performed by mixing 10 nM of purified DNA and Blunt/TA ligase cloning mastermix in a total of 10 μL and incubating 15 min at 25 °C. Crude ligation product was diluted 10-fold and 1 μL was used for PCR amplification in 100 μL using 2 Units of Q5 polymerase, 0.2 mM dNTP mix, 0.5 μM of primers T395/T396 and 1X Evagreen 20X. Sample was incubated in a real-time thermocycler (Miniopticon, BioRad) using program **PCR 22**. Amplified DNA was purified (Zymo Research), eluted with 15 µL milliQ water, yielding the N-terminal sequences. The C-terminal sequences were retrieved by PCR amplification of 1 μL of the 10-fold diluted broken emulsion as follows: In 100 μL were mixed 2 Units of Q5 polymerase, 0.2 mM dNTP mix, 0.5 μM of primers T397/T398 and 1X Evagreen 20X. Sample was incubated in a qPCR thermocycler (Miniopticon, BioRad) using program **PCR 23**. Amplified DNA was purified (Zymo Research), eluted with 30 µL milliQ water, yielding the C-terminal sequences. Samples Nter and Cter were pooled and sent for Illumina sequencing analysis using Genewiz Amplicon-EZ platform.

#### AlphaFold model

DNK structure was determined using the AlphaFold3 webserver, in presence of AMP and magnesium, AMP and ATP without magnesium, or AMP and ATP and magnesium. The model aligns well with the crystal structure of T4 DNK in complex with dGMP and AMP (PDB 1DEL), the AMP in the T5 model taking the place of the dGMP acceptor of the T4 crystal structure Supplementary Figure 13.

### Millipede device

We adapted the millipede microfluidic design introduced by Amstad et al. (Lab Chip, 2016), which enables the high-speed production of monodisperse droplets ranging from 20 to 200 µm in diameter. First, the original design was uniformly scaled down by maintaining all length ratios and angles, reducing the characteristic dimension ‘a’ to 10 µm (Supplementary Figure 14). Second, given the strong dependence of droplet formation on the width-to-height (w/h) ratio, the height of the lower channel layer was reduced to 3.5 µm, yielding a w/h ratio of approximately 14.3, below the threshold of 19 specified in the original work. The primary PDMS layer directly to a glass substrate, eliminating the need for secondary PDMS support. As glass wettability can disrupt droplet uniformity, the channels were rendered hydrophobic by infusing a silane-based treatment prior to each experiment, ensuring consistent surface chemistry and reliable droplet generation.

### Microfluidic device production

Device master moulds were prepared using standard soft-lithography techniques according to (Gines et al., 2024). Briefly, a photoresist (SU-8, MicroChem) was spin-coated to the desired height on a 4-inch dehydrated silicon wafer, followed by ultraviolet exposure through the nozzle and filter mask phototraced on a transparent film (Selba). Following development in propylene glycol methyl ether acetate, the operation (spin-coating, exposure, development) was repeated to include the rest of the channels. The mold was aligned with the mask using an MJB4 instrument (SUSS MicroTec). Devices were replicated from the master mould using polydimethylsiloxane 10:1 base/curing agent (Sylgard 184, Dow Corning). After baking 1 h at 70 °C, the polydimethylsiloxane slab was peeled off, the inlet and outlet were drilled using a 1.5 mm biopsy puncher (Integra Miltex). The PDMS layer was then bonded to a 1 mm-thick glass microscope slide after oxygen plasma activation, followed by thermal bonding at 200 °C for 5 hours. For the millipede device, channels were rendered hydrophobic by infusing the device with an electronic-grade fluorinated coating (Novec 1720, 3M), which was subsequently removed before a final bake at 90 °C for 1 hour.

### Microfluidic droplet generation

Sample partitioning was performed by a step-emulsification process using either a one-inlet disperse phase device or a millipede device. The continuous phase consisted of a fluorinated oil (HFE Novec 7500, 3 M) supplemented with 1% w/w (one-inlet) or 4% w/w (millipede) Fluosurf-C (Emulseo). Samples and oil were connected to the chip using PTFE tubing (inner diameter, 200 µm; Cluzeau Info Labo) and injected using a pressure-based flow controller (MFCS EZ pump, Fluigent). Droplets were collected in a pipette tip plugged into the outlet.

### End-point imaging

Following incubation, emulsions were imaged in end-point by microscopy as previously described (Gines et al., 2020, 2024; Menezes et al., 2020). In brief, a glass slide and coverslip were hydrophobized with Novec 1720 (3M). Polystyrene beads of a diameter close to the droplet size were placed at the four corners of the coverslip and used as spacer to control the thickness of the droplet monolayer. The chamber was sealed using an epoxy glue (Sader) and imaged with an epifluorescence microscope (Nikon Eclipse Ti) equipped with a motorized stage, a Nikon DS-Qi2 camera, a CoolLed pE-4000 illumination source, a filter set (Semrock, DyLight405-C, FITC-3540C, Cy3-4040C, mCherry-C and LF635/LP-B) and a 20× apochromatic objective (numerical aperture, 0.75; working distance, 1.0 mm) for wild type *dnk* experiment or an epifluorescence microscope (Nikon Eclipse Ti2) equipped with a motorized stage, a Hamamatsu ORCA-Flash4.0 camera a Lumencor MultiLaser Sectra/Aura light source, a LED-DA/FI/TR/Cy5-A filter cube (DAPI / FITC / TRITC / Cy5 - Full Multiband Quad) and a 10x Plan Apo λ objective (numerical aperture, 0.45) for the mini library experiment. False-color images were generated using ImageJ open-source software.

### Droplet analysis

Droplet images were analyzed using image J and custom python script. A threshold was used either on the far red (when Alexa 680 fluorophore was present in all droplets) or Brightfield channel was used to segment the droplets. Droplets were detected using image J particle module and ROIs were reduced by 50% to only measure gray value of the center of the droplet. Mean fluorescence was plotted and a gaussian was fitted on the empty droplet peaks excluding outliers; the positive droplet threshold was set at three times the standard deviation from the gaussian mean value.

### Mini library model

Model parameters were fitted using a homemade python script. Briefly, the function to optimize computed all possible droplet compositions and their probability up to three co-encapsulations, determined the global DNK activity in the droplet and the resulting DNA composition after replication, considering an equal sharing of the resources. The function’s parameters are the input library genotype proportions, an activity factor for each variant and RBS, the Poisson factor λ, and the three parameters of the replication function RepF. RepF is a linear relationship between two minimal and maximal plateau values of k (number of co-encapsulated genotypes) and *MaxRep*, respectively. The linear segment has two parameters: a slope *a*, and an intercept *b*. This correspondence between RBS, variant factors and DNA replication translates into the following equations:

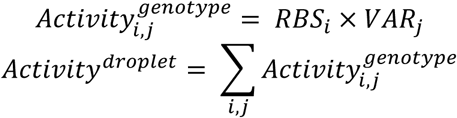

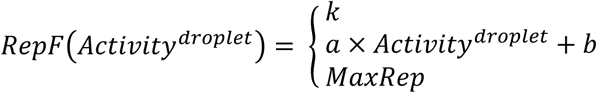

The wild-type DNK and the RBS-5 parameters were fixed to 100 to anchor the model. This left 16 free parameters to fit: 5 variant activity coefficients (relative to WT), 6 RBS strength coefficients (relative to RBS-5), the effective λ (loading factor), and the RepF parameters *a, b* and *MaxRep*. The parameters were fitted by minimizing the error between observed and model-predicted output composition. Parameter uncertainties were estimated via bootstrap resampling of the sequencing counts (input library ∼9.1×10^4^ reads, output ∼2.75×10^5^ reads).

### SQI calculation

As SQI is normally calculated from mock library experiments, we had to consider two populations of “active” and “inactive” genotypes. For the inactive group we considered all genotypes with efficiency of RBS-1_T17S and lower, and for the active group we selected RBS-5_T17S, RBS-AT_T17S and RBS-AT_WT. Proportion of active variants before and after replication is of 0.21 and 0.74, respectively. SQI was calculated according to (Dramé-Maigné et al., 2020), using the equation for selection.

## Data accessibility

NGS data are available on the European Nucleotide Archive, accession number PRJEB96338.

## Conflict-of-interest

The authors declare no conflict-of-interest.

## Funding

This research was supported by the European Research Council [ERC-CoG-2014 ProFF ID: 647275]. The microfluidics was realized at the Pierre-Gilles de Gennes Institute (IPGG) with the support of the Equipex/Idex ANR program (Equipex ANR: 10-EQPX-0034).

## Aknowledgments

Proteins P5 and P6 were kindly provided by the late Margarita Salas laboratory. We thank Taro Furubayashi for fruitful discussions and Rocío Espada for valuable assistance with next-generation sequencing.

## PCR program table

|  |  |  |
| --- | --- | --- |
| PCR 1 | 98 °C 30 s; (98 °C 10 s; 60 °C 10 s; 72 °C 45 s) x3; (98 °C 10 s; 72 °C 45 s) x17; 72 °C 2 min | Barcoded X_OriLR1-12 parts |
| PCR 2 | 98 °C 30 s; (98 °C 10 s; 72 °C 25 s) x20; 72 °C 2 min | Barcoded X_OriLR1-12 parts |
| PCR 3 | 98 °C 30 s; (98 °C 10 s; 60 °C 10 s; 72 °C 30 s) x21; 72 °C 2 min | UMI-bearing Y_OriLumi and Y_OriRumi parts |
| PCR 4 | 98 °C 30 s; (98 °C 10 s; 50 °C 10 s; 72 °C 20 s) x18; 72 °C 2 min | UMI-bearing Y_OriLumi and Y_OriRumi parts |
| PCR 5 | 98 °C 30 s; (98 °C 10 s; X °C 10 s; 72 °C 2 min) x20; 72 °C 2 min | dnk point-mutation plasmids |
| PCR 6 | 98 °C 30 s; (98 °C 10 s; 56 °C 10 s; 72 °C 45 s) x2; (98 °C 10 s; 72 °C 55 s) x20; 72 °C 2 min | X_RBS_mCherry_dnk_T7term parts |
| PCR 7 | 98 °C 30 s; (98 °C 10 s; 68 °C 10 s; 72 °C 72 s) x9; 72 °C 2 min | X_RBS_mCherry_dnk_T7term parts |
| PCR 8 | 98 °C 30 s; (98 °C 10 s; 69 °C 10 s; 72 °C 1 min) x14; 72 °C 2 min | Addition of UMI |
| PCR 9 | 98 °C 30 s; (98 °C 10 s; 70 °C 10 s; 72 °C 20 s) x25; 72 °C 2 min | Barcoded acceptor vector for Gibson Assembly pGibs1 |
| PCR 10 | 98 °C 30 s; (98 °C 10 s; 61 °C 10 s; 72 °C 20 s) x2; 72 °C 2 min | Barcoded acceptor vector for Gibson |
| PCR 11 | 94 °C 30 s; (94 °C 30 s; 62 °C 30 s; 68 °C 4 min) x30; 68 °C 5 min | Barcoded acceptor vector for Gibson Assembly pGibs2 |
| PCR 12 | 98 °C 30 s; (98 °C 10 s; 64 °C 20 s; 72 °C 1 min 39 s) x2; 72 °C 2 min | Barcoded acceptor vector for Gibson Assembly pGibs2 |
| PCR 13 | 98 °C 2 min; (98 °C 30 s; 66 °C 30 s; 72 °C 1 min 18) x30; 72 °C 10 min | DNK random mutational libraries creation |
| PCR 14 | 98 °C 30 s; (98 °C 10 s; 64 °C 15 s; 72 °C 25 s) x2; (98 °C 10 s; 72 °C 40 s) x28; 72 °C 2 min | DNK random mutational libraries creation |
| PCR 15 | 98 °C 30 s; (98 °C 10 s; 69 °C 15 s; 72 °C 40 s) x20; 72 °C 2 min | DNK random mutational libraries creation |
| PCR 16 | 98 °C 30 s; (98 °C 15 s; 69 °C 20 s; 72 °C 40 s) x30; 72 °C 2 min | Error prone library selection |
| PCR 17 | 98 °C 30 s; (98 °C 15 s; 69 °C 10 s; 72 °C 30 s) x20; 72 °C 2 min | Error prone library selection |
| PCR 18 | 98 °C 30 s; (98 °C 10 s; 69 °C 10 s; 72 °C 30 s) x20; 72 °C 2 min | Error prone library selection |
| PCR 19 | 98 °C 30 s; (98 °C 10 s; 60 °C 10 s; 72 °C 30 s) x12; 72 °C 2 min | Error prone library selection |
| PCR 20 | 98 °C 30 s; (98 °C 10 s; (56 or 69) °C 20 s; 72 °C 30 s) x4; (98 °C 10 s; 72 °C 50 s) xN; 72 °C 2 min | Illumina sequencing libraries L0 and L1 |
| PCR 21 | 98 °C 30 s; (98 °C 10 s; 58 °C 10 s; 72 °C 22 s) x12; 72 °C 2 min | Illumina sequencing with UMI flip libraries L1 L2 L3 |
| PCR 22 | 98 °C 30 s; (98 °C 10 s; 59 °C 10 s; 72 °C 5 s) x19; 72 °C 2 min | Illumina sequencing with UMI flip libraries L1 L2 L3 |
| PCR 23 | 98 °C 30 s; (98 °C 10 s; 58 °C 10 s; 72 °C 5 s) x20; 72 °C 2 min | Illumina sequencing with UMI flip libraries L1 L2 L3 |
| PCR 24 | 98 °C 30 s; (98 °C 10 s; 69 °C 10 s; 72 °C 1 min) x14; 72 °C 2 min | Amplification of <gfp> |

## Notes

https://www.ebi.ac.uk/ena/browser/view/PRJEB96338

### Competing Interest Statement

The authors have declared no competing interest.

