## Supplementary Material for "Functional selection in a population of synthetic cells with a minimal metabolism"

### SUPPLEMENTARY MATERIAL
### For
### Functional selection in a population of synthetic cells with a minimal metabolism

T. Di Meo, L. Bunel, G. Ragala, M. Van Tongeren, R. Sieskind, C. Danelon, Y. Rondelez*

| | Input library | Output library |
|---|---|---|
| **Passed Bonito sequences** | 184000 | 490000 |
| **+/-200 bp reference size** | 120000 | 381000 |
| **Aligned to reference** | 120000 | 381000 |
| **Alignments score >55** | 119005 (100%) | 363105 (100%) |
| **Matching tags** | 97193 (82%) | 290863 (80%) |
| **Unmatching tags** | 6639 (5,6%) | 25812 (7,1%) |
| **At least one unassigned tag** | 15173 (13%) | 46430 (13%) |
| **Unassigned RBS** | 7785 (6,5%) | 20150 (5,5%) |
| **Matching tags and assigned RBS** | 91223 (77%) | 275441 (76%) |

**Supplementary Table 1.** Summary of nanopore sequencing and analysis in input and output selection of mini library. Chimera formation is a known pitfall of amplification of homologous DNA (Boers et al., 2015; Karst et al., 2021; Lahr & and Katz, 2009; Meyerhans et al., 1990; Shao et al., 2011). Only the sequences with assigned and matching left and right barcode were used for analysis.

| Input | RBS-1 | RBS-2 | RBS-3 | RBS-4 | RBS-C | RBS-5 | RBS-AT |
|---|---|---|---|---|---|---|---|
| **R172I** | 1303 | 1387 | 1007 | 1978 | 1636 | 1461 | 3255 |
| **D16N** | 2305 | 2213 | 1287 | 2603 | 3025 | 2298 | 2731 |
| **T17S** | 1842 | 1932 | 1199 | 3020 | 2492 | 2416 | 3534 |
| **K131E** | 1783 | 1793 | 1392 | 1800 | 1868 | 1952 | 2018 |
| **W150F** | 2214 | 2396 | 1834 | 2802 | 2783 | 2495 | 2048 |
| **WT** | 1774 | 2368 | 1654 | 3232 | 2853 | 2682 | 2558 |

| Output | RBS-1 | RBS-2 | RBS-3 | RBS-4 | RBS-C | RBS-5 | RBS-AT |
|---|---|---|---|---|---|---|---|
| **R172I** | 898 | 1042 | 631 | 2194 | 2521 | 1829 | 1851 |
| **D16N** | 1625 | 1372 | 1369 | 3054 | 4610 | 3508 | 8438 |
| **T17S** | 1483 | 898 | 5325 | 13466 | 14481 | 20043 | 29032 |
| **K131E** | 1448 | 1996 | 3066 | 5187 | 5706 | 8547 | 8125 |
| **W150F** | 2746 | 1949 | 5881 | 5956 | 7473 | 10596 | 10466 |
| **WT** | 1581 | 823 | 9569 | 7597 | 11861 | 17665 | 27533 |

**Supplementary Table 2.** Counts of each genotype in input and output libraries, using reads with assigned and matching left and right barcode.

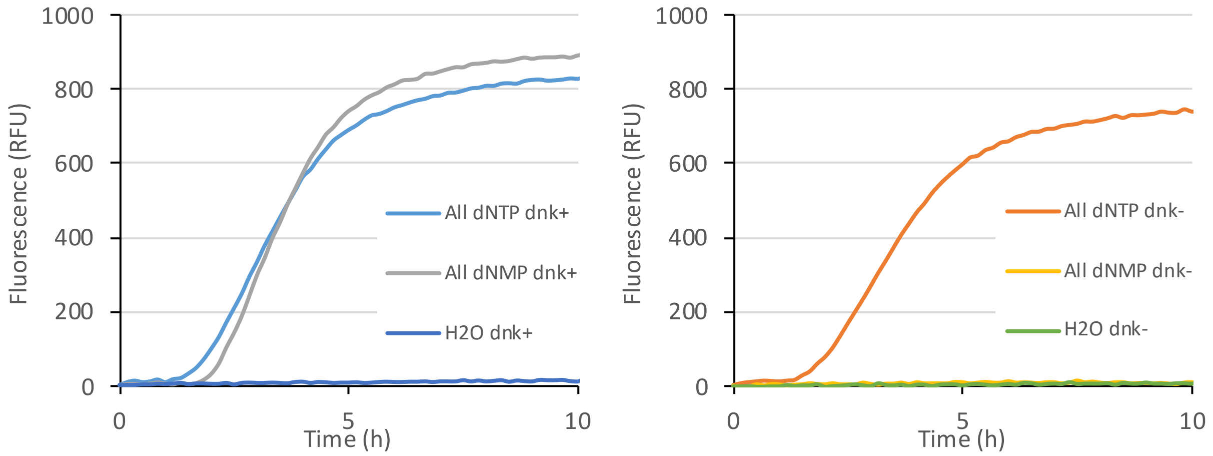


**Supplementary Figure 1.** Baseline-corrected GFP fluorescence of <*rfp*> replicator in IVTTR conditions in different dNTP conditions with or without 25 pM *dnk* gene. Raw fluorescence in channel Cal Orange 560 is shown, as fluorescence on FAM channel was saturating.

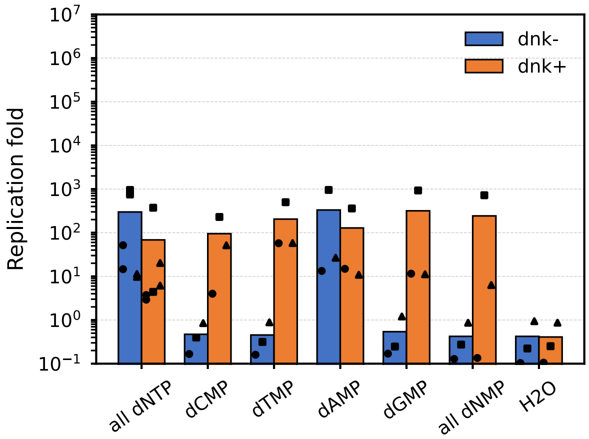


**Supplementary Figure 2.** qPCR quantification of <*gfp*> replicator fragment after IVTTR experiment in bulk. In each aliquot of a same IVTTR mastermix a different dNTP mixture was added where one or all of the dNTP was replaced with the corresponding dNMP, in absence or presence of 25 pM of *dnk*. Data point shape indicate same mastermix.

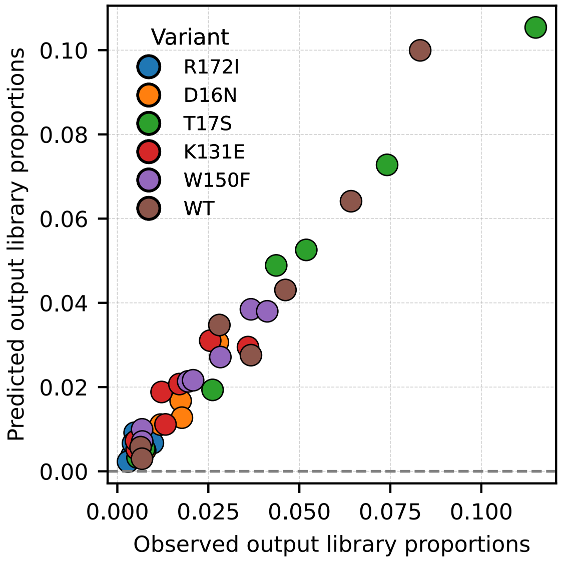


**Supplementary Figure 3.** Quality of fit of the model. Output library proportions of each genotype plotted as observed vs predicted by the model. RMSD of fit: 0.0009.

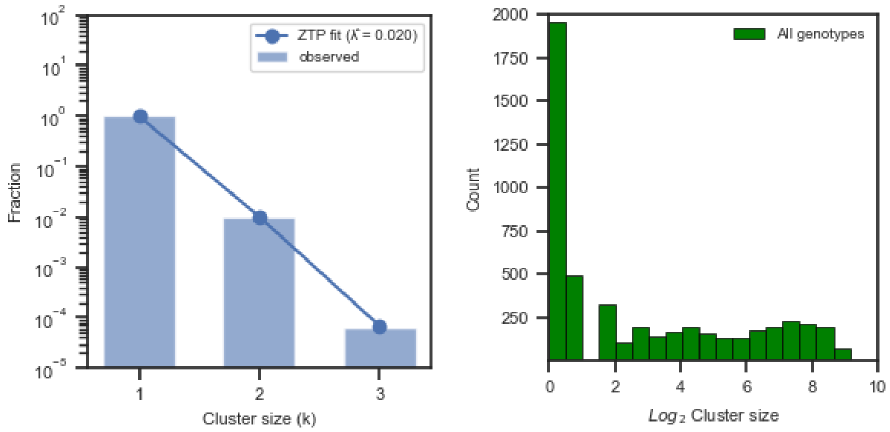


**Supplementary Figure 4. Left.** Distribution of cluster size in the input library sequencing for one genotype (15546 reads) (blue bars). Fitted curve with a zero-truncated Poisson distribution at lambda 0.018 (blue line). Estimated UM diversity is 8x10$^5$. Same result is observed across all genotypes. **Right.** Distribution of cluster size in $Log_2$ scale in the output library: 5133 different clusters were found in total.

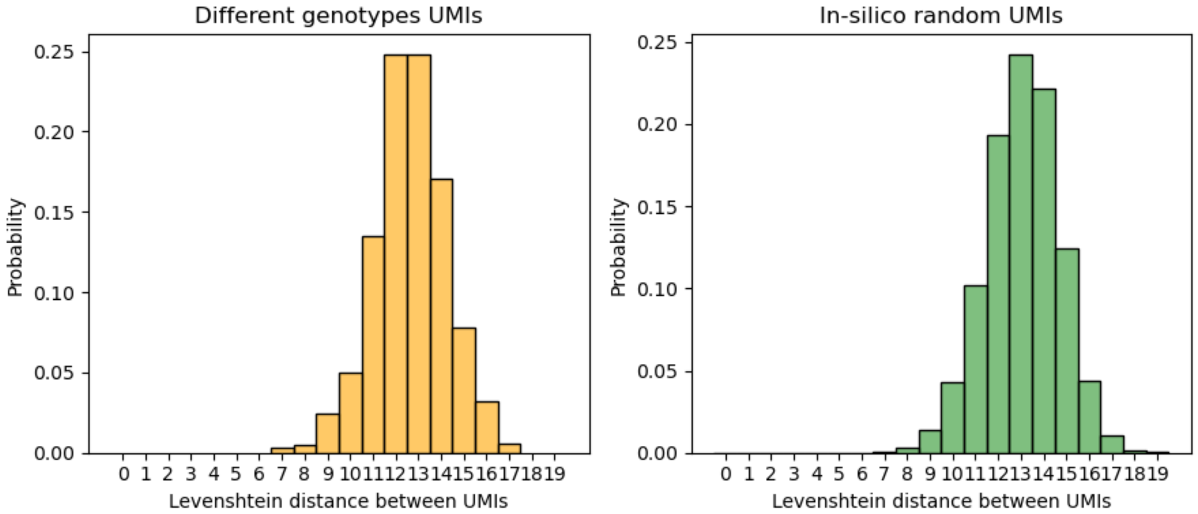


**Supplementary Figure 5.** Analysis of UMI Levenshtein distance between randomly picked UMI of input library (blue) versus randomly generated UMI (yellow). Truly orthogonal UMI were taken inside input library by sampling pairs using two different genotypes (RBS-AT_WT and RBS-C_T17S). Cross-Levenshtein distances of 10000 generated UMI matching pattern NNWNNNNNWNNNNWNNNNNWNN were computed, showing a mean Levenshtein distance of 13. Using clustering cutoff distance of 6 gives 0.01% false-positive rate.

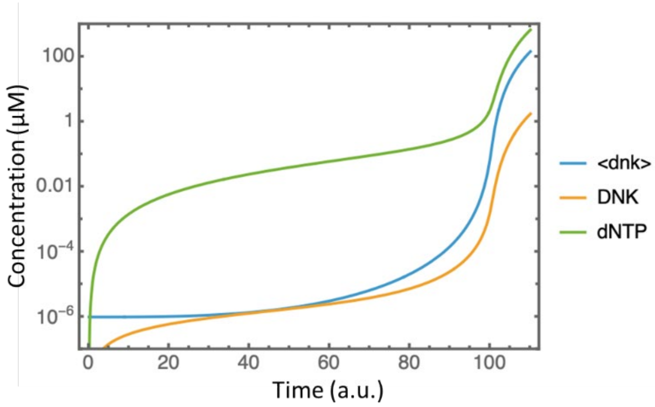


**Supplementary Figure 6.** Simulated time-trace (log scale) of <*dnk*>, DNK and dNTP concentration. The model shown in Figure 6c was integrated over a hundred time steps. The parameters were set to $k_1 = 0.03$, $k_2 = 1000$, $k_3 = 1$, $K_1 = 10$ µM, $K_2 = 1$ µM; At $t$=0, <*dnk*>, DNK and dNTP were set to 1 pM, 0 and 0, respectively. The upper and lower dashed curves and the shaded area show the <*dnk*> evolution when $k_3$ is replaced by 0.9, or 1.1, respectively.

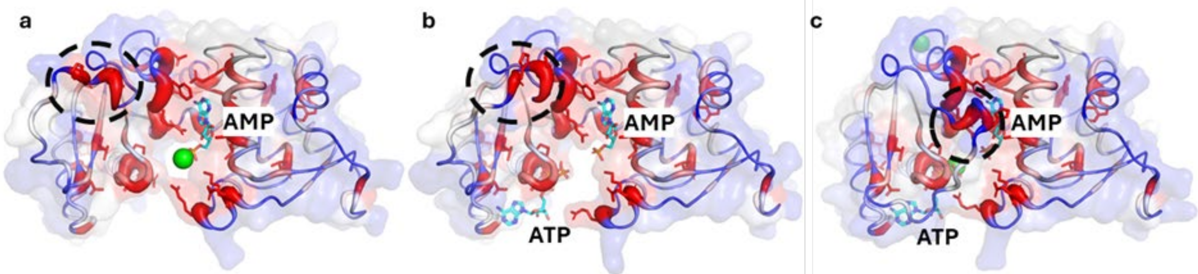


**Supplementary Figure 7.** AlphaFold3 model of T5 DNK including AMP and magnesium ions (a) AMP and ATP without magnesium (b), and AMP and ATP and two magnesium ions (c). These structures align well with the reported structure of T4 DNK complexed with dGMP and AMP (PDB 1DEL) where AMP and ATP of the model take the place of dGMP and AMP of the crystal structure, respectively. One can observe closed lid (black dashed circle)

only in presence of ATP, AMP and magnesium, as it has been reported in silico with adenylate kinase (Nam et al., 2024).

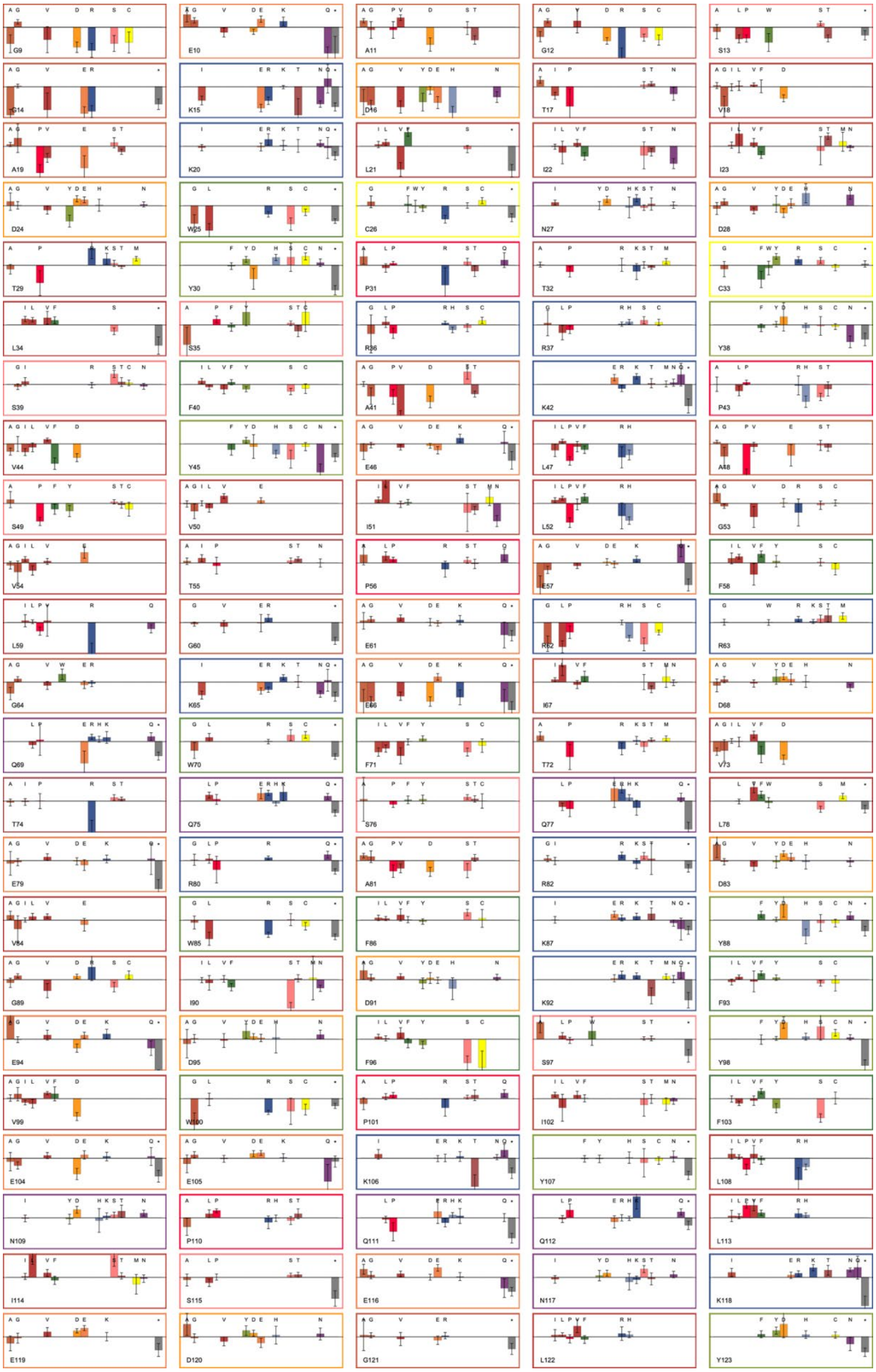

**Supplementary Figure 8.** log-jump in frequency, Log[$f_1/f_0$] computed for all scored positions. The error bars are evaluated from the counts of each mutation in L0 and L1, assuming Poissonian distribution.

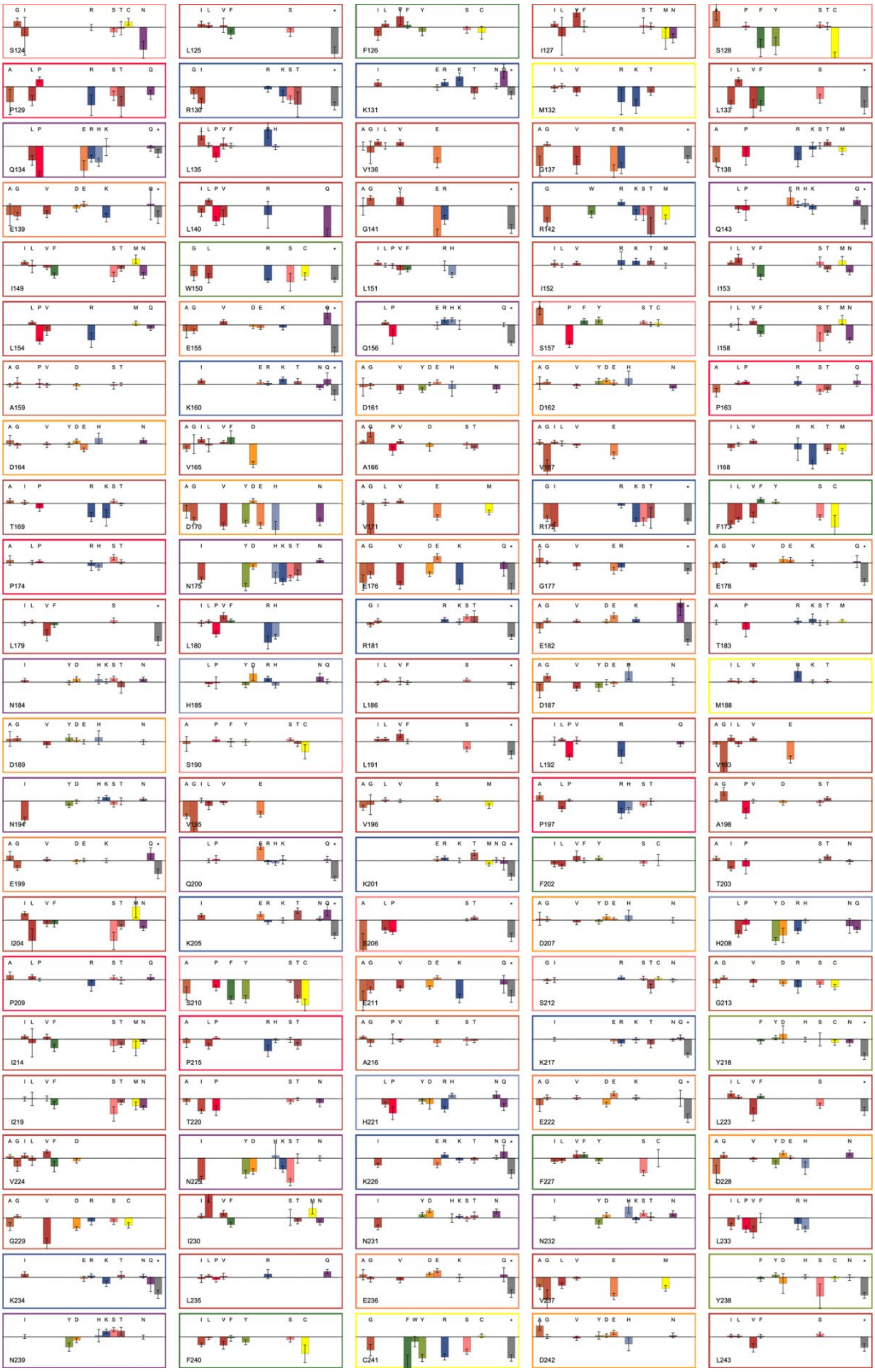

**Supplementary Figure 9.** Same as Supplementary Figure 8.

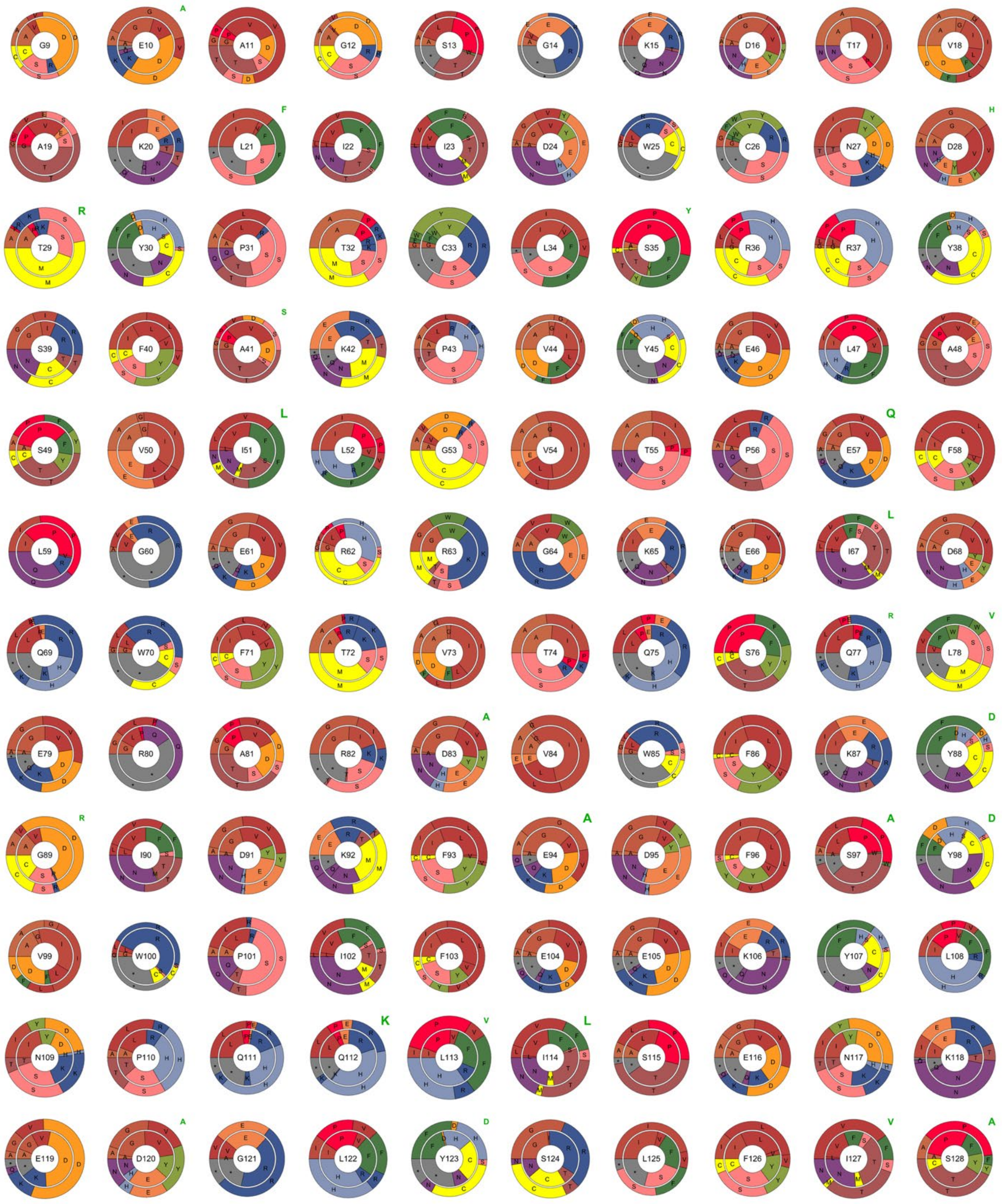

**Supplementary Figure 10.** Representation of the evolution of mutation frequencies between the naive and the selected library (first round) for all scored positions. Each sector chart gives the relative observed frequency of the indicated mutations for naive (inner) and selected (outer) library. The width of inner and outer sector indicates the total frequency rate. Therefore, positions that strongly reject mutations have a thin outer sector, while positions that accept most mutations have a wider one. Mutations with a fitness contribution Log[$f_1/f_0$] greater than Log(2) are highlighted in green, with font size related to the selection strength.

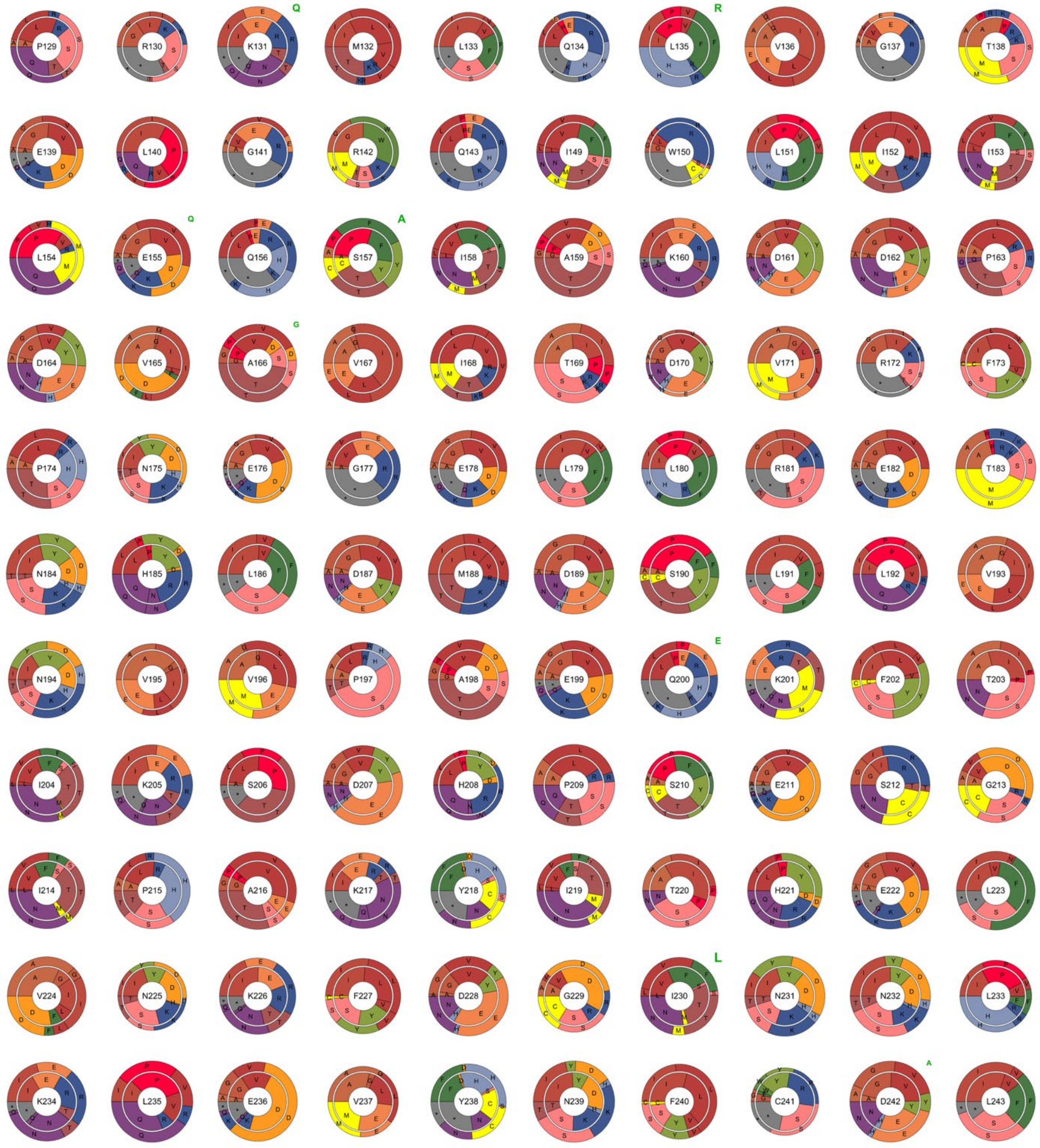


**Supplementary Figure 11.** Same as Supplementary Figure 10.

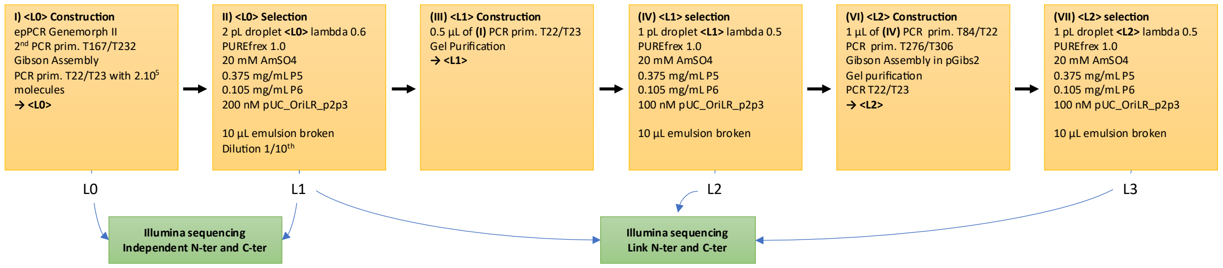


**Supplementary Figure 12.** Workflow for the creation and selection of error prone libraries.

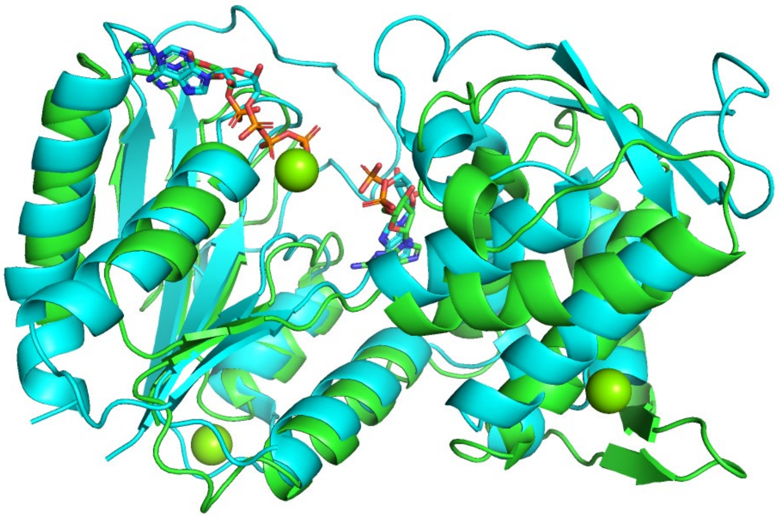

**Supplementary Figure 13.** Alphafold3 model of DNK from T5 phage (blue) in presence of ATP (left), AMP (center) and two magnesium ions (blue sphere) aligned with crystal structure of DNK from phage T4 (green, PDB 1del) in complex with AMP (left) and dGMP (center). Left domain align with an RMSD of 2.5 Å.

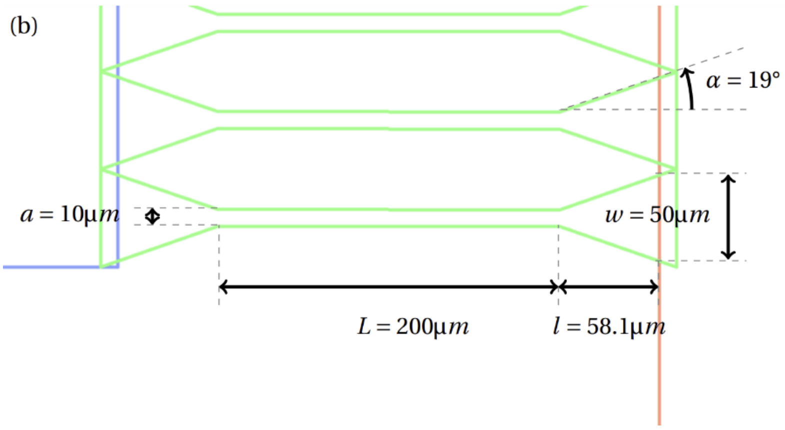


**Supplementary Figure 14.** Millipede design. The green pattern represents the lower layer. The blue pattern is linked to the inlet and the red one to the outlet. Both are parts of the upper layer.

**Supplementary Table 3. Primers and dsDNA sequences used.**

| Primers names | Sequence |
|---|---|
| GR25 | GAACGTCGCAGTCCCTCTAG |
| GR26 | GAAGCAGGCACGGTAAGACA |
| iR120 | CCTCAAGACCCGTTTAGAG |
| iR184 | TTGCTGGCCTTTTGCTCAC |
| iR186 | CACATTTCCCCGAAAAGTGCC |
| M111-Barcode1-R172I-F | ATCGTCTCGTTACAAGGCGAAATGCGAAGAGCTGCCGGAGACCGTCTG |
| M112-Barcode1-R172I-R | TACGTCTCGTCTGTCTCCGAACATCGAAGAGCTTGCGGAGACCGCTGA |
| M113-Barcode2-D16N-F | ATCGTCTCGTTACTGCGACAAATGCGAAGAGCTGCCGGAGACCGTCTG |
| M114-Barcode2-D16N-R | TACGTCTCGTCTGCCACACAACATCGAAGAGCTTGCGGAGACCGCTGA |
| M115-Barcode3-T17S-F | ATCGTCTCGTTACCTTGCCAAATGCGAAGAGCTGCCGGAGACCGTCTG |
| M116-Barcode3-T17S-R | TACGTCTCGTCTGAGTGTACCCATCGAAGAGCTTGCGGAGACCGCTGA |
| M117-Barcode4-K131I-F | ATCGTCTCGTTACGCAGCTAAATGCGAAGAGCTGCCGGAGACCGTCTG |
| M118-Barcode4-K131I-R | TACGTCTCGTCTGCCTTAGGACATCGAAGAGCTTGCGGAGACCGCTGA |
| M119-Barcode5-W150F-F | ATCGTCTCGTTACGCAATGGAATGCGAAGAGCTGCCGGAGACCGTCTG |
| M120-Barcode5-W150F-R | TACGTCTCGTCTGTTCGTGGACATCGAAGAGCTTGCGGAGACCGCTGA |
| M121-Barcode6-WT-F | ATCGTCTCGTTACCACAGTGAATGCGAAGAGCTGCCGGAGACCGTCTG |
| M122-Barcode6-WT-R | TACGTCTCGTCTGAACCTCCACATCGAAGAGCTTGCGGAGACCGCTGA |
| M123-Barcode7-rbsAT-T172I-F | ATCGTCTCGTTACATCCGGTAATGCGAAGAGCTGCCGGAGACCGTCTG |
| M124-Barcode7-rbsAT-T172I-R | TACGTCTCGTCTGCGAGGTTACATCGAAGAGCTTGCGGAGACCGCTGA |
| M125-Barcode8-rbsAT-D16N-F | ATCGTCTCGTTACTAGAGAGGATGCGAAGAGCTGCCGGAGACCGTCTG |
| M126-Barcode8-rbsAT-D16N-R | TACGTCTCGTCTGAGACTTGGCATCGAAGAGCTTGCGGAGACCGCTGA |
| M127-Barcode9-rbsAT-T17S-F | ATCGTCTCGTTACTGTAAGCGATGCGAAGAGCTGCCGGAGACCGTCTG |
| M128-Barcode9-rbsAT-T17S-R | TACGTCTCGTCTGCTAACTCGCATCGAAGAGCTTGCGGAGACCGCTGA |
| M129-Barcode10-rbsAT-K131I-F | ATCGTCTCGTTACACCATCTGATGCGAAGAGCTGCCGGAGACCGTCTG |
| M130-Barcode10-rbsAT-K131I-R | TACGTCTCGTCTGCTGTTCTGCATCGAAGAGCTTGCGGAGACCGCTGA |
| M131-Barcode11-rbsAT-W150F-F | ATCGTCTCGTTACTGGTGTTGATGCGAAGAGCTGCCGGAGACCGTCTG |
| M132-Barcode11-rbsAT-W150F-R | TACGTCTCGTCTGTATGGCACCATCGAAGAGCTTGCGGAGACCGCTGA |
| M133-Barcode12-rbsAT-WT-F | ATCGTCTCGTTACGGCATTACATGCGAAGAGCTGCCGGAGACCGTCTG |
| M134-Barcode12-rbsAT-WT-R | TACGTCTCGTCTGGATAGTCCCATCGAAGAGCTTGCGGAGACCGCTGA |
| M139-RBS-1 | TCGTCTCGAACTTAGGGCCCGTTAGGTTCCGATGgttagtaaaggtgaag |

| | |
|---|---|
| M140-RBS-2 | TCGTCTCGAACTAGAGTTAATTAGGTACTAAATGgttagtaaaggtgaag |
| M143-RBS-3 | TCGTCTCGAACTTTTGTTTAACTAATTTAAGGAGGTTTTGTAATGgttagtaaaggtgaag |
| M141-RBS-4 | TCGTCTCGAACTTTTTTAAGGAGGTAGCGATATGgttagtaaaggtgaag |
| M142-RBS-5 | TCGTCTCGAACTTTGTAAGGAGGTTTTTTTAATGgttagtaaaggtgaag |
| M137-RBS-C | ATCGTCTCGAACTTTAAGAAGgagatataccATGgttagtaaaggtgaagaggataac |
| MT01-RBS-AT | GGCGTCTCGAACTTTAAGAAGGAGATATACATATG |
| M84 | CCATCGTCTCGGTAACATGATTACTACAGTACAAAAAACCCCTCAAGAC |
| M173-YUMI-For | TCGGTCTCCTGCCNNWNNNNNWNNTGCCCGAGACGGAACATCCACACTTTAGTGA |
| M174-YUMI-rev | TCGGTCTCCTTGCNNWNNNNNWNNTTGCCGAGACGGCGAAGCATATAAAAGAAG C |
| T36 | AAAGTAAGCCCCCACCCTC |
| M190 | TCGGTCTCCTTGC |
| T37 | AAAGTAGGGTACAGCGACAAC |
| M191 | TCGGTCTCCTGCC |
| M76 | TGCGTCTCGTGCCGGAGACCGTCTGATCAACCTTCACACAGATC |
| M77 | AGCGTCTCGTTGCGGAGACCGCTGAACTGAATTCTAAGGC |
| M78 | TCGGTCTCCTGCCCGAGACGGAACATCCACACTTTAGTGA |
| M79 | TCGGTCTCCTTGCCGAGACGGCGAAGCATATAAAAGAAGC |
| T84 | CAAAAAACCCCTCAAGACCCGTTTAGAGG |
| MT01 | GGCGTCTCGAACTTTAAGAAGGAGATATACATATG |
| T22 | 5'P-AAAGTAAGCCCCCACCCTCACATG |
| T23 | 5'P-AAAGTAGGGTACAGCGACAACATACAC |
| T242-R172I-0-for | ATTttccccaatgaaggagagttac |
| T243-R172I-0-rev | cacatctgttattacggcgac |
| T244-D16N-16-for | AACaccgttgcaaaattaattatcg |
| T245-D16N-16-rev | ttttcccgaaccagcttcac |
| T246-T17S-68-for | TCGgttgcaaaattaattatcgattgg |
| T247-T17S-68-rev | atcttttcccgaaccagcttc |
| T248-K131E-86-for | GAAatgttacagctcgtaggaacg |
| T249-K131E-86-rev | tctcggagaaataaataaactataaag |
| T250-W150F-117-for | TTCctcataattctggagcaatcc |
| T251-W150F-117-rev | aatgcgttcatgtaccagctg |
| T133-qPCR-dnk-F | CCCGTTTATGAACTTGCATCCG |

| | |
|---|---|
| **T134-qPCR-dnk-R** | **TCGGCCAAACGTACGAGAAG** |
| **T129-qPCR-gfp-F** | **GGAGCTTTTCACTGGCGTTGTTC** |
| **T130-qPCR-gfp-R** | **GGTGGTGCAGATGAACTTCAGG** |
| **T274-primer-vector-2** | **TGTCACAACTCGACGATCTCHVBDHVBDHVBDHVBDHVBDHVBDHVBDHVBDHVB CCTATGACACCAACGTTGGGGCTTCTAATAGCATAACCCC** |
| **T304-primer-vector-2rev** | **ggcaatgtctaacatgacttacagHVBDHVBDHVBDHVBDHVBDHVBDHVBDHVBDHVBT GCGACACAGACGAAGCGCTAAAACG** |
| **T275** | **TGTCACAACTCGACGATCTC** |
| **T305** | **ggcaatgtctaacatgacttacag** |
| **T393** | **5'P-AACTTTAAGAAGGAGATATACCATG** |
| **T394** | **5'P-GACAGATCCTAACTAGTCA** |
| **T395** | **AAATCTACTGGGACTCAAAAT** |
| **T396** | **cgagctgtaacatttttctcg** |
| **T397** | **cggactttatagtttatttatttctc** |
| **T398** | **GCGACAGATCCTAACTAGTC** |

| **dsDNAb names** | **Sequence** |
|---|---|
| pAcc_pXFus_dnk | cacatttccccgaaaagtgccacctgacgtctaagaaaccattattatcatgacattaacctataaaaataggcgtatcacgaggcccttAAAGTAAGCCC<br>CCACCCTCACATGATACCATTCTCCTAATATCGACATAATCCGTCGATCCTCGGCATACCATGATCAGGGAGGGAAACTAC<br>TACTTAATATATCAATCTATAGACCTACTAGATAGGTTTGTCAATGAACAACATAAAACGACACAGAATCCCACGTTTTAGC<br>GCTTCGTCTGTGTCGCAGAAGCAGGCACGGTAAGACACGGTCTAGCTGACTGTCTATCGGCTAGGTCAAATAGGGAGCTT<br>TGATATCTGCGTGTCCAGCCTTAGAATTCAGTTCAGCGGTCTCCGCAAGCTCTTCGATGTAGTTACTACCTCTAAACAGATT<br>GTAATACGACTCACTATAGGGATAATACAACGGTTTCCCTCTAgaaataattttgtttaactttaagaaggagatatacatATGgttagtaa<br>aggtgaagaggataacatggccatcatcaaggagttcatgcgcttcaaggtgcacatggagggctccgtgaacggccacgagttcgagatcgagggcgaggg<br>cgagggccgcccctacgagggcacccagaccgccaagctgaaggtgaccaagggtggccccctgcccttcgcctgggacatcctgtcccctcagttcatgtacg<br>gctccaaggcctacgtgaagcaccccgccgacatccccgactacttgaagctgtccttccccgagggcttcaagtgggagcgcgtgatgaacttcgaggacggc<br>ggcgtggtgaccgtgacccaggactcctccctgcaggacggcgagttcatctacaaggtgaagctgcgcggcaccaacttcccctccgacggccccgtaatgca<br>gaagaagaccatgggctgggaggcctcctccgagcggatgtaccccgaggacggcgccctgaagggcgagatcaagcagaggctgaagctgaaggacggc<br>ggccactacgacgctgaggtcaagaccacctacaaggccaagaagcccgtgcagctgcccggcgcctacaacgtcaacatcaagttggacatcacctcccaca<br>acgaggactacaccatcgtggaacagtacgaacgcgccgagggccgccactccaccggcggcatggacgagctgtacaagGGTGGCGGTGGCAGTT<br>TAGTGCCACGTGGTTCAGGCGGTGGCGGTTCTATGTCAGTATTAGTCGGTTTACATGGTGAAGCTGGTTCGGGAAAAGAT<br>ACCGTTGCAAAATTAATTATCGATTGGTGTAATGACACGTATCCAACGTGTTTATCCCGTCGTTACAGTTTTGCTAAGCCCG<br>TTTATGAACTTGCATCCGTAATCCTCGGCGTAACTCCAGAGTTTCTAGGAGAGCGCAGGGGGAAAGAGATTGACCAATGG<br>TTTACGGTCACACAATCTCAATTGGAGCGAGCTCGTGACGTATGGTTCAAATACGGTATCGATAAGTTTGAGGACTTCTCG<br>TACGTTTGGCCGATTTTTGAGGAAAAATATCTTAATCCTCAACAACTTATCTCAGAGAATAAAGAAGACGGACTTTATAGT<br>TTATTTATTTCTCCGAGAAAAATGTTACAGCTCGTAGGAACGGAGCTAGGAAGGCAGCTGGTACATGAACGCATTTGGCT<br>CATAATTCTGGAGCAATCCATCGCTAAAGACGACCCAGATGTCGCCGTAATAACAGATGTGAGATTCCCCAATGAAGGAG<br>AGTTACTCCGTGAAACGAACCATTTAGATATGGATTCTTTACTAGTAAATGTAGTGCCCGCTGAGCAGAAGTTCACTATTA<br>AATCAGATCATCCATCTGAAAGTGGTATTCCTGCAAAATATATCACTCACGAATTAGTTAATAAATTCGATGGTATCAATAA<br>CCTTAAACTAGAAGTGTATAACTTCTGTGACTTAGAGCTAGAACCACTAGTTGGATAAATAGTCAatcttcggaaatcccATGCC<br>TCTAATACTACACACTTAGCATAACCCCTTGGGGCCTCTAAACGGGTCTTGAGGGGTTTTTTGTACTGTAGTAATCATGTTA<br>CAGATGTTCTGTATTCCAATGCGAAGAGCTGCCGGAGACCGTCTGATCAACCTTCACACAGATCTAGAATCAAAAGCAGT |

| | |
|---|---|
| | GATCTCCCGGGTGCGAAATAAAAATACTAGGTAACTAGAGGGACTGCGACGTTCCCTCCTATGATTGGTTGTCTTATTACC<br>TTACTTCTATTATAGTATAACATGTTAAACGATAGTTTGTCTACCCTTTTCGACAAATTGATGATAATAAATAGTATAGGTAT<br>ATAGTCGTGATTTAGTTGTTAGATTCTTGTCGAAGATAGTCGGTCAATGGGGAAATGGTGTATGTTGTCGCTGTACCCTAC<br>TTTtgcgctcggtcgttcggctgcggcgagcggtatcagctcactcaaaggcggtaatacggttatccacagaatcaggggataacgcaggaaagaacatgtg<br>agcaaaaggccagcaaaaggccaggaaccgtaaaaaggccgcgttgctggcgtttttccataggctccgcccccctgacgagcatcacaaaaatcgacgctc<br>aagtcagaggtggcgaaacccgacaggactataaagataccaggcgtttccccctggaagctccctcgtgcgctctcctgttccgaccctgccgcttaccggata<br>cctgtccgcctttctcccttcgggaagcgtggcgctttctcatagctcacgctgtaggtatctcagttcggtgtaggtcgttcgctccaagctgggctgtgtgcacga<br>accccccgttcagcccgaccgctgcgccttatccggtaactatcgtcttgagtccaacccggtaagacacgacttatcgccactggcagcagccactggtaacag<br>gattagcagagcgaggtatgtaggcggtgctacagagttcttgaagtggtggcctaactacggctacactagaagaacagtatttggtatctgcgctctgctgaa<br>gccagttaccttcggaaaaagagttggtagctcttgatccggcaaacaaaccaccgctggtagcggtggtttttttgtttgcaagcagcagattacgcgcagaaa<br>aaaaggatctcaagaagatcctttgatcttttctacggggtctgacgctcagtggaacgaaaactcacgttaagggattttggtcatgagattatcaaaaaggat<br>cttcacctagatccttttaaattaaaaatgaagttttaaatcaatctaaagtatatatgagtaaacttggtctgacagttaccaatgcttaatcagtgaggcacctа<br>tctcagcgatctgtctatttcgttcatccatagttgcctgactccccgtcgtgtagataactacgatacgggagggcttaccatctggccccagtgctgcaatgata<br>ccgcgactcccacgctcaccggctccagatttatcagcaataaaccagccagccggaagggccgagcgcagaagtggtcctgcaactttatccgcctccatcca<br>gtctattaattgttgccgggaagctagagtaagtagttcgccagttaatagtttgcgcaacgttgttgccattgctacaggcatcgtggtgtcacgctcgtcgtttg<br>gtatggcttcattcagctccggttcccaacgatcaaggcgagttacatgatcccccatgttgtgcaaaaaagcggttagctccttcggtcctccgatcgttgtcag<br>aagtaagttggccgcagtgttatcactcatggttatggcagcactgcataattctcttactgtcatgccatccgtaagatgcttttctgtgactggtgagtactcaac<br>caagtcattctgagaatagtgtatgcggcgaccgagttgctcttgcccggcgtcaatacgggataataccgcgccacatagcagaactttaaaagtgctcatcat<br>tggaaaacgttcttcggggcgaaaactctcaaggatcttaccgctgttgagatccagttcgatgtaacccactcgtgcacccaactgatcttcagcatcttttactt<br>tcaccagcgtttctgggtgagcaaaaacaggaaggcaaaatgccgcaaaaaagggaataagggcgacacggaaatgttgaatactcatactcttcctttttca<br>atattattgaagcatttatcagggttattgtctcatgagcggatacatatttgaatgtatttagaaaaataaacaaataggggttccgcg |
| pUC_<br>OriLR_<br>p2p3 | ACCGCATCAGGCGCCATTCGCCATTCAGGCTGCGCAACTGTTGGGAAGGGCGATCGGTGCGGGCCTCTTCGCTATTACGC<br>CAGCTGGCGAAAGGGGGATGTGCTGCAAGGCGATTAAGTTGGGTAACGCCAGGGTTTTCCCAGTCACGACGTTGTAAAA<br>CGACGGCCAGTGAATTCGAGCTCGGTACCTCGCGAATGCATCTAGATCCAATAAAGTAAGCCCCCACCCTCACATGATACC<br>ATTCTCCTAATATCGACATAATCCGTCGATCCTCGGCATACCATGATCAGGGAGGGAAACTACTACTTAATATATCAATCTA<br>TAGACCTACTAGATAGGTTTGTCAATGAACAACATAAAACGACACAGAATCCCACGTTTTAGCGCTTCGTCTGTGTCGCAT<br>GTGAAATTAATACGACTCACTATAGGGAGACCACAACGGTTTCCCTCTAGAAATAATTTTGTTTAACTTTAAGAAGGAGAT<br>ATACATATGCCGCGTAAAATGTACAGCTGCGATTTTGAAACGACGACGAAAGTTGAAGATTGCCGTGTCTGGGCCTATGG<br>TTATATGAACATCGAAGACCATTCAGAATATAAAATTGGCAACTCGCTGGATGAATTTATGGCGTGGGTGCTGAAAGTTC<br>AGGCCGACCTGTACTTCCACAATCTGAAATTTGATGGTGCGTTCATTATCAACTGGCTGGAACGTAATGGCTTTAAATGGA<br>GCGCCGATGGTCTGCCGAACACCTATAATACGATTATCTCTCGTATGGGCCAATGGTATATGATTGATATCTGCCTGGGCT<br>ACAAAGGTAAACGCAAAATTCATACCGTGATCTATGACAGCCTGAAAAAACTGCCGTTTCCGGTGAAGAAAATTGCGAAA<br>GATTTCAAACTGACCGTCCTGAAAGGCGATATTGACTATCACAAAGAACGTCCGGTTGGTTACAAAATCACGCCGGAAGA<br>ATATGCGTACATTAAAAACGATATCCAGATTATCGCAGAAGCTCTGCTGATTCAGTTTAAACAAGGCCTGGATCGCATGAC<br>CGCCGGCAGTGACTCCCTGAAAGGTTTCAAAGATATCATCACCACGAAAAAATTTAAGAAAGTGTTCCCGACCCTGAGCC<br>TGGGTCTGGATAAAGAAGTTCGTTATGCATACCGCGGCGGTTTTACGTGGCTGAACGACCGTTTCAAAGAAAAAGAAATT<br>GGCGAGGGTATGGTCTTTGATGTGAATAGTCTGTATCCGGCTCAGATGTACTCCCGCCTGCTGCCGTATGGCGAACCGAT<br>CGTTTTCGAGGGTAAATATGTCTGGGATGAAGACTACCCGCTGCATATTCAGCACATCCGTTGTGAATTTGAACTGAAAGA<br>AGGCTATATTCCGACCATTCAAATCAAACGTAGCCGCTTCTATAAGGGTAACGAATACCTGAAAAGCTCTGGCGGTGAAA<br>TCGCAGACCTGTGGCTGAGTAACGTCGATCTGGAACTGATGAAAGAACATTACGATCTGTACAACGTTGAATACATCTCC<br>GGCCTGAAATTTAAAGCCACCACGGGTCTGTTTAAAGATTTCATTGACAAATGGACCTACATCAAAACCACGTCTGAAGGT<br>GCAATCAAACAGCTGGCTAAACTGATGCTGAACAGCCTGTATGGCAAATTTGCATCTAATCCGGATGTTACCGGTAAAGT<br>CCCGTACCTGAAAGAAAATGGCGCTCTGGGTTTTCGCCTGGGCGAAGAAGAAACCAAAGATCCGGTGTATACGCCGATG<br>GGTGTTTTCATTACCGCGTGGGCACGTTACACCACCATCACCGCCGCACAAGCGTGCTATGACCGCATTATCTACTGTGAT<br>ACCGACTCAATTCATCTGACCGGCACGGAAATCCCGGATGTGATTAAAGATATCGTTGACCCGAAAAAACTGGGTTATTG<br>GGCACACGAATCGACCTTTAAACGTGCTAAATACCTGCGCCAGAAAACGTACATCCAAGACATCTACATGAAAGAAGTCG<br>ATGGCAAACTGGTGGAAGGTTCACCGGATGACTATACCGACATTAAATTTTCGGTGAAATGCGCCGGCATGACCGATAAA<br>ATTAAGAAAGAAGTGACGTTCGAAAATTTCAAAGTGGGTTTCAGTCGCAAAATGAAACCGAAACCGGTCCAAGTTCCGG<br>GCGGCGTTGTGCTGGTCGATGACACCTTCACGATCAAATAAGAATTGTACTAGAGTATCTGTTAGTTTTTTTCTACTAGAG<br>TACTAGAGTATCTGTTAGTTTTTTTCATCGGATCCCGGGCCCGTCGACTGCTAATACGACTCACTATAGGGAGACCACAAC<br>GGTTTCCCTCTAGAAATAATTTTGTTTAACTTTAAGAAGGAGATATACATATGGCACGCAGCCCGCGCATCCGCATCAAAG<br>ATAACGACAAAGCCGAATACGCCCGCCTGGTGAAAAATACGAAAGCTAAAATCGCACGTACCAAGAAAAAATATGGCGT<br>GGATCTGACGGCTGAAATTGACATCCCGGATCTGGACTCATTTGAAACCCGCGCGCAGTTCAATAAATGGAAAGAACAAG<br>CTAGCTCTTTTACGAACCGTGCGAATATGCGCTATCAGTTCGAGAAAAACGCCTACGGTGTGGTTGCATCGAAAGCTAAA<br>ATTGCGGAAATCGAACGTAACACCAAAGAAGTTCAACGCCTGGTCGATGAAAAAATCAAAGCCATGAAAGACAAAGAAT<br>ATTACGCAGGCGGTAAACCGCAGGGCACGATTGAACAACGTATCGCCATGACCTCACCGGCACATGTGACGGGTATCAA<br>CCGTCCGCACGATTTTGACTTCAGTAAAGTTCGCAGTTACTCCCGTCTGCGCACCCTGGAAGAATCCATGGAAATGCGCAC<br>GGATCCGCAGTATTACGAAAAGAAAATGATTCAGCTGCAACTGAATTTTATCAAAAGCGTCGAAGGCTCATTTAACTCGTT<br>CGATGCGGCCGACGAACTGATTGAAGAACTGAAGAAAATTCCGCCGGATGACTTTTATGAACTGTTCCTGCGTATTAGCG<br>AAATCTCTTTTGAAGAATTCGATTCTGAAGGCAACACCGTCGAAAATGTGGAGGGTAACGTTTACAAAATTCTGTCGTATC<br>TGGAACAATATCGTCGTGGTGATTTTGATCTGTCGCTGAAAGGCTTCTAATAGCATAACCCCTTGGGGCCTCTAAACGGGT<br>CTTGAGGGGTTTTTTGCCTCCTATGATTGGTTGTCTTATTACCTTACTTCTATTATAGTATAACATGTTAAACGATAGTTTGT<br>CTACCCTTTTCGACAAATTGATGATAATAAATAGTATAGGTATATAGTCGTGATTTAGTTGTTAGATTCTTGTCGAAGATAG |

| | |
|---|---|
| | TCGGTCAATGGGGAAATGGTGTATGTTGTCGCTGTACCCTACTTTATTGGATCGGATCCCGGGCCCGTCGACTGCAGAGG<br>CCTGCATGCAAGCTTGGCGTAATCATGGTCATAGCTGTTTCCTGTGTGAAATTGTTATCCGCTCACAATTCCACACAACATA<br>CGAGCCGGAAGCATAAAGTGTAAAGCCTGGGGTGCCTAATGAGTGAGCTAACTCACATTAATTGCGTTGCGCTCACTGCC<br>CGCTTTCCAGTCGGGAAACCTGTCGTGCCAGCTGCATTAATGAATCGGCCAACGCGCGGGGAGAGGCGGTTTGCGTATT<br>GGGCGCTCTTCCGCTTCCTCGCTCACTGACTCGCTGCGCTCGGTCGTTCGGCTGCGGCGAGCGGTATCAGCTCACTCAAAG<br>GCGGTAATACGGTTATCCACAGAATCAGGGGATAACGCAGGAAAGAACATGTGAGCAAAAGGCCAGCAAAAGGCCAGG<br>AACCGTAAAAAGGCCGCGTTGCTGGCGTTTTTCCATAGGCTCCGCCCCCCTGACGAGCATCACAAAAATCGACGCTCAAG<br>TCAGAGGTGGCGAAACCCGACAGGACTATAAAGATACCAGGCGTTTCCCCCTGGAAGCTCCCTCGTGCGCTCTCCTGTTCC<br>GACCCTGCCGCTTACCGGATACCTGTCCGCCTTTCTCCCTTCGGGAAGCGTGGCGCTTTCTCATAGCTCACGCTGTAGGTAT<br>CTCAGTTCGGTGTAGGTCGTTCGCTCCAAGCTGGGCTGTGTGCACGAACCCCCCGTTCAGCCCGACCGCTGCGCCTTATCC<br>GGTAACTATCGTCTTGAGTCCAACCCGGTAAGACACGACTTATCGCCACTGGCAGCAGCCACTGGTAACAGGATTAGCAG<br>AGCGAGGTATGTAGGCGGTGCTACAGAGTTCTTGAAGTGGTGGCCTAACTACGGCTACACTAGAAGAACAGTATTTGGT<br>ATCTGCGCTCTGCTGAAGCCAGTTACCTTCGGAAAAAGAGTTGGTAGCTCTTGATCCGGCAAACAAACCACCGCTGGTAG<br>CGGTGGTTTTTTTGTTTGCAAGCAGCAGATTACGCGCAGAAAAAAAGGATCTCAAGAAGATCCTTTGATCTTTTCTACGGG<br>GTCTGACGCTCAGTGGAACGAAAACTCACGTTAAGGGATTTTGGTCATGAGATTATCAAAAAGGATCTTCACCTAGATCCT<br>TTTAAATTAAAAATGAAGTTTTAAATCAATCTAAAGTATATATGAGTAAACTTGGTCTGACAGTTACCAATGCTTAATCAGT<br>GAGGCACCTATCTCAGCGATCTGTCTATTTCGTTCATCCATAGTTGCCTGACTCCCCGTCGTGTAGATAACTACGATACGG<br>GAGGGCTTACCATCTGGCCCCAGTGCTGCAATGATACCGCGAGACCCACGCTCACCGGCTCCAGATTTATCAGCAATAAA<br>CCAGCCAGCCGGAAGGGCCGAGCGCAGAAGTGGTCCTGCAACTTTATCCGCCTCCATCCAGTCTATTAATTGTTGCCGGG<br>AAGCTAGAGTAAGTAGTTCGCCAGTTAATAGTTTGCGCAACGTTGTTGCCATTGCTACAGGCATCGTGGTGTCACGCTCGT<br>CGTTTGGTATGGCTTCATTCAGCTCCGGTTCCCAACGATCAAGGCGAGTTACATGATCCCCCATGTTGTGCAAAAAAGCGG<br>TTAGCTCCTTCGGTCCTCCGATCGTTGTCAGAAGTAAGTTGGCCGCAGTGTTATCACTCATGGTTATGGCAGCACTGCATA<br>ATTCTCTTACTGTCATGCCATCCGTAAGATGCTTTTCTGTGACTGGTGAGTACTCAACCAAGTCATTCTGAGAATAGTGTAT<br>GCGGCGACCGAGTTGCTCTTGCCCGGCGTCAATACGGGATAATACCGCGCCACATAGCAGAACTTTAAAAGTGCTCATCA<br>TTGGAAAACGTTCTTCGGGGCGAAAACTCTCAAGGATCTTACCGCTGTTGAGATCCAGTTCGATGTAACCCACTCGTGCAC<br>CCAACTGATCTTCAGCATCTTTTACTTTCACCAGCGTTTCTGGGTGAGCAAAAACAGGAAGGCAAAATGCCGCAAAAAAG<br>GGAATAAGGGCGACACGGAAATGTTGAATACTCATACTCTTCCTTTTTCAATATTATTGAAGCATTTATCAGGGTTATTGTC<br>TCATGAGCGGATACATATTTGAATGTATTTAGAAAAATAAACAAATAGGGGTTCCGCGCACATTTCCCCGAAAAGTGCCA<br>CCTGACGTCTAAGAAACCATTATTATCATGACATTAACCTATAAAAATAGGCGTATCACGAGGCCCTTTCGTCTCGCGCGTT<br>TCGGTGATGACGGTGAAAACCTCTGACACATGCAGCTCCCGGAGACGGTCACAGCTTGTCTGTAAGCGGATGCCGGGAG<br>CAGACAAGCCCGTCAGGGCGCGTCAGCGGGTGTTGGCGGGTGTCGGGGCTGGCTTAACTATGCGGCATCAGAGCAGATT<br>GTACTGAGAGTGCACCATATGCGGTGTGAAATACCGCACAGATGCGTAAGGAGAAAAT |
| pUC_<br>OriLR_<br>gfp | GTCACGACGTTGTAAAACGACGGCCAGTGAATTCGAGCTCGGTACCTCGCGAATGCATCTAGATCCATAAAGTAAGCCCC<br>CACCCTCACATGATACCATTCTCCTAATATCGACATAATCCGTCGATCCTCGGCATACCATGATCAGGGAGGGAAACTACT<br>ACTTAATATATCAATCTATAGACCTACTAGATAGGTTTGTCAATGAACAACATAAAACGACACAGAATCCCACGTTTTAGC<br>GCTTCGTCTGTGTCGCAGGGCCCGTCGACTGCTAATACGACTCACTATAGGGAGACCACAACGGTTTCCCTCTAGAAATAA<br>TTTTGTTTAACTTTAAGAAGGAGATATACATATGGAGCTTTTCACTGGCGTTGTTCCCATCCTGGTCGAGCTGGACGGCGA<br>CGTAAACGGCCACAAGTTCAGCGTGTCCGGCGAGGGCGAGGGCGATGCCACCTACGGCAAGCTGACCCTGAAGTTCATC<br>TGCACCACCGGCAAGCTGCCCGTGCCCTGGCCCACCCTCGTGACCACCCTGACCTACGGCGTGCAGTGCTTCAGCCGCTAC<br>CCCGACCACATGAAGCAGCACGACTTCTTCAAGTCCGCCATGCCCGAAGGCTACGTCCAGGAGCGCACCATCTTCTTCAAG<br>GACGACGGCAACTACAAGACCCGCGCCGAGGTGAAGTTCGAGGGCGACACCCTGGTGAACCGCATCGAGCTGAAGGGC<br>ATCGACTTCAAGGAGGACGGCAACATCCTGGGGCACAAGCTGGAGTACAACTACAACAGCCACAACGTCTATATCATGGC<br>CGACAAGCAGAAGAACGGCATCAAGGTGAACTTCAAGATCCGCCACAACATCGAGGACGGCAGCGTGCAGCTCGCCGAC<br>CACTACCAGCAGAACACCCCCATCGGCGACGGCCCCGTGCTGCTGCCCGACAACCACTACCTGAGCACCCAGTCCGCCCT<br>GAGCAAAGACCCCAACGAGAAGCGCGATCACATGGTCCTGCTGGAGTTCGTGACCGCCGCCGGGATCTAAGATCTGTCG<br>CTGAAAGGCTTCTAATAGCATAACCCCTTGGGGCCTCTAAACGGGTCTTGAGGGGTTTTTTGCCTCCTATGATTGGTTGTC<br>TTATTACCTTACTTCTATTATAGTATAACATGTTAAACGATAGTTTGTCTACCCTTTTCGACAAATTGATGATAATAAATAGT<br>ATAGGTATATAGTCGTGATTTAGTTGTTAGATTCTTGTCGAAGATAGTCGGTCAATGGGGAAATGGTGTATGTTGTCGCTG<br>TACCCTACTTTATTGGATCGGATCCCGGGCCCGTCGACTGCAGAGGCCTGCATGCAAGCTTGGCGTAATCATGGTCATAG<br>CTGTTTCCTGTGTGAAATTGTTATCCGCTCACAATTCCACACAACATACGAGCCGGAAGCATAAAGTGTAAAGCCTGGGGT<br>GCCTAATGAGTGAGCTAACTCACATTAATTGCGTTGCGCTCACTGCCCGCTTTCCAGTCGGGAAACCTGTCGTGCCAGCTG<br>CATTAATGAATCGGCCAACGCGCGGGGAGAGGCGGTTTGCGTATTGGGCGCTCTTCCGCTTCCTCGCTCACTGACTCGCT<br>GCGCTCGGTCGTTCGGCTGCGGCGAGCGGTATCAGCTCACTCAAAGGCGGTAATACGGTTATCCACAGAATCAGGGGAT<br>AACGCAGGAAAGAACATGTGAGCAAAAGGCCAGCAAAAGGCCAGGAACCGTAAAAAGGCCGCGTTGCTGGCGTTTTTCC<br>ATAGGCTCCGCCCCCCTGACGAGCATCACAAAAATCGACGCTCAAGTCAGAGGTGGCGAAACCCGACAGGACTATAAAG<br>ATACCAGGCGTTTCCCCCTGGAAGCTCCCTCGTGCGCTCTCCTGTTCCGACCCTGCCGCTTACCGGATACCTGTCCGCCTTT<br>CTCCCTTCGGGAAGCGTGGCGCTTTCTCATAGCTCACGCTGTAGGTATCTCAGTTCGGTGTAGGTCGTTCGCTCCAAGCTG<br>GGCTGTGTGCACGAACCCCCCGTTCAGCCCGACCGCTGCGCCTTATCCGGTAACTATCGTCTTGAGTCCAACCCGGTAAGA<br>CACGACTTATCGCCACTGGCAGCAGCCACTGGTAACAGGATTAGCAGAGCGAGGTATGTAGGCGGTGCTACAGAGTTCTT<br>GAAGTGGTGGCCTAACTACGGCTACACTAGAAGAACAGTATTTGGTATCTGCGCTCTGCTGAAGCCAGTTACCTTCGGAA<br>AAAGAGTTGGTAGCTCTTGATCCGGCAAACAAACCACCGCTGGTAGCGGTGGTTTTTTTGTTTGCAAGCAGCAGATTACG<br>CGCAGAAAAAAAGGATCTCAAGAAGATCCTTTGATCTTTTCTACGGGGTCTGACGCTCAGTGGAACGAAAACTCACGTTA<br>AGGGATTTTGGTCATGAGATTATCAAAAAGGATCTTCACCTAGATCCTTTTAAATTAAAAATGAAGTTTTAAATCAATCTAA |

| | |
|---|---|
| | AGTATATATGAGTAAACTTGGTCTGACAGTTACCAATGCTTAATCAGTGAGGCACCTATCTCAGCGATCTGTCTATTTCGTT<br>CATCCATAGTTGCCTGACTCCCCGTCGTGTAGATAACTACGATACGGGAGGGCTTACCATCTGGCCCCAGTGCTGCAATGA<br>TACCGCGAGACCCACGCTCACCGGCTCCAGATTTATCAGCAATAAACCAGCCAGCCGGAAGGGCCGAGCGCAGAAGTGG<br>TCCTGCAACTTTATCCGCCTCCATCCAGTCTATTAATTGTTGCCGGGAAGCTAGAGTAAGTAGTTCGCCAGTTAATAGTTTG<br>CGCAACGTTGTTGCCATTGCTACAGGCATCGTGGTGTCACGCTCGTCGTTTGGTATGGCTTCATTCAGCTCCGGTTCCCAA<br>CGATCAAGGCGAGTTACATGATCCCCCATGTTGTGCAAAAAAGCGGTTAGCTCCTTCGGTCCTCCGATCGTTGTCAGAAGT<br>AAGTTGGCCGCAGTGTTATCACTCATGGTTATGGCAGCACTGCATAATTCTCTTACTGTCATGCCATCCGTAAGATGCTTTT<br>CTGTGACTGGTGAGTACTCAACCAAGTCATTCTGAGAATAGTGTATGCGGCGACCGAGTTGCTCTTGCCCGGCGTCAATA<br>CGGGATAATACCGCGCCACATAGCAGAACTTTAAAAGTGCTCATCATTGGAAAACGTTCTTCGGGGCGAAAACTCTCAAG<br>GATCTTACCGCTGTTGAGATCCAGTTCGATGTAACCCACTCGTGCACCCAACTGATCTTCAGCATCTTTTACTTTCACCAGC<br>GTTTCTGGGTGAGCAAAAACAGGAAGGCAAAATGCCGCAAAAAAGGGAATAAGGGCGACACGGAAATGTTGAATACTC<br>ATACTCTTCCTTTTTCAATATTATTGAAGCATTTATCAGGGTTATTGTCTCATGAGCGGATACATATTTGAATGTATTTAGAA<br>AAATAAACAAATAGGGGTTCCGCGCACATTTCCCCGAAAAGTGCCACCTGACGTCTAAGAAACCATTATTATCATGACATT<br>AACCTATAAAAATAGGCGTATCACGAGGCCCTTTCGTCTCGCGCGTTTCGGTGATGACGGTGAAAACCTCTGACACATGC<br>AGCTCCCGGAGACGGTCACAGCTTGTCTGTAAGCGGATGCCGGGAGCAGACAAGCCCGTCAGGGCGCGTCAGCGGGTG<br>TTGGCGGGTGTCGGGGCTGGCTTAACTATGCGGCATCAGAGCAGATTGTACTGAGAGTGCACCATATGCGGTGTGAAAT<br>ACCGCACAGATGCGTAAGGAGAAAATACCGCATCAGGCGCCATTCGCCATTCAGGCTGCGCAACTGTTGGGAAGGGCGA<br>TCGGTGCGGGCCTCTTCGCTATTACGCCAGCTGGCGAAAGGGGGATGTGCTGCAAGGCGATTAAGTTGGGTAACGCCAG<br>GGTTTTCCCA |
| X_T7p<br>romot<br>er | ATGGCGTCTCGCAGAATCACTGACGCGATTGTAATACGACTCACTATAGGgataatACAACGGTTtccctctagaaataattTTGT<br>TTAACTCGAGACGATGG |
| gBlock<br>T7_RB<br>S-<br>AT_dn<br>k_wt | CAGATAATACGACTCACTATAGGGATAATACAACGGTTTCCCTCTAGAAATAATTTTGTTTAACTTTAAGAAGGAGATATA<br>CATATGTCAGTATTAGTCGGTTTACATGGTGAAGCTGGTTCGGGAAAAGATACCGTTGCAAAATTAATTATCGATTGGTGT<br>AATGACACGTATCCAACGTGTTTATCCCGTCGTTACAGTTTTGCTAAGCCCGTTTATGAACTTGCATCCGTAATCCTCGGCG<br>TAACTCCAGAGTTTCTAGGAGAGCGCAGGGGGAAAGAGATTGACCAATGGTTTACGGTCACACAATCTCAATTGGAGCG<br>AGCTCGTGACGTATGGTTCAAATACGGTATCGATAAGTTTGAGGACTTCTCGTACGTTTGGCCGATTTTTGAGGAAAAATA<br>TCTTAATCCTCAACAACTTATCTCAGAGAATAAAGAAGACGGACTTTATAGTTTATTTATTTCTCCGAGAAAAATGTTACAG<br>CTCGTAGGAACGGAGCTAGGAAGGCAGCTGGTACATGAACGCATTTGGCTCATAATTCTGGAGCAATCCATCGCTAAAGA<br>CGACCCAGATGTCGCCGTAATAACAGATGTGAGATTCCCCAATGAAGGAGAGTTACTCCGTGAAACGAACCATTTAGATA<br>TGGATTCTTTACTAGTAAATGTAGTGCCCGCTGAGCAGAAGTTCACTATTAAATCAGATCATCCATCTGAAAGTGGTATTC<br>CTGCAAAATATATCACTCACGAATTAGTTAATAAATTCGATGGTATCAATAACCTTAAACTAGAAGTGTATAACTTCTGTGA<br>CTTAGAGCTAGAACCACTAGTTGGATAAATAGTCAATCTTCGGAAATCCCATGCCGTATGAGTAGAGATTTGTAAGAGCT<br>GTTAGTTAGC |
| pEX-<br>A128-<br>pAnob<br>ar | acctgcttttgctcgcttggatccgaattcaaaggtgaaattgttatccgctcacaattccacacaacatacgagccggaagcataaagtgtaaagcctggggtg<br>cctaatgagtgagctaactcacattaattgcgttgcgctcactgcccgctttccagtcgggaaacctgtcgtgccagctgcattaatgaatcggccaacgcgcgg<br>ggagaggcggtttgcgtattgggcgctcttccgcttcctcgctcactgactcgctgcgctcggtcgttcggctgcggcgagcggtatcagctcactcaaaggcggt<br>aatacggttatccacagaatcaggggataacgcaggaaagaacatgtgagcaaaaggccagcaaaaggccaggaaccgtaaaaaggccgcgttgctggcgt<br>ttttccataggctccgcccccctgacgagcatcacaaaaatcgacgctcaagtcagaggtggcgaaacccgacaggactataaagataccaggcgtttccccct<br>ggaagctccctcgtgcgctctcctgttccgaccctgccgcttaccggatacctgtccgcctttctcccttcgggaagcgtggcgctttctcatagctcacgctgtagg<br>tatctcagttcggtgtaggtcgttcgctccaagctgggctgtgtgcacgaaccccccgttcagcccgaccgctgcgccttatccggtaactatcgtcttgagtccaa<br>cccggtaagacacgacttatcgccactggcagcagccactggtaacaggattagcagagcgaggtatgtaggcggtgctacagagttcttgaagtggtggcct<br>aactacggctacactagaagaacagtatttggtatctgcgctctgctgaagccagttaccttcggaaaaagagttggtagctcttgatccggcaaacaaaccac<br>cgctggtagcggtggtttttttgtttgcaagcagcagattacgcgcagaaaaaaaggatctcaagaagatcctttgatcttttctacggggtctgacgctcagtgg<br>aacgaaaactcacgttaagggattttggtcatgagattatcaaaaaggatcttcacctagatccttttaaattaaaaatgaagttttaaatcaatctaaagtatat<br>atgagtaaacttggtctgacagttaccaatgcttaatcagtgaggcacctatctcagcgatctgtctatttcgttcatccatagttgcctgactccccgtcgtgtaga<br>taactacgatacgggagggcttaccatctggccccagtgctgcaatgataccgcgactcccacgctcaccggctccagatttatcagcaataaaccagccagcc<br>ggaagggccgagcgcagaagtggtcctgcaactttatccgcctccatccagtctattaattgttgccgggaagctagagtaagtagttcgccagttaatagtttg<br>cgcaacgttgttgccattgctacaggcatcgtggtgtcacgctcgtcgtttggtatggcttcattcagctccggttcccaacgatcaaggcgagttacatgatcccc<br>catgttgtgcaaaaaagcggttagctccttcggtcctccgatcgttgtcagaagtaagttggccgcagtgttatcactcatggttatggcagcactgcataattctc<br>ttactgtcatgccatccgtaagatgcttttctgtgactggtgagtactcaaccaagtcattctgagaatagtgtatgcggcgaccgagttgctcttgcccggcgtca<br>atacgggataataccgcgccacatagcagaactttaaaagtgctcatcattggaaaacgttcttcggggcgaaaactctcaaggatcttaccgctgttgagatcc<br>agttcgatgtaacccactcgtgcacccaactgatcttcagcatcttttactttcaccagcgtttctgggtgagcaaaaacaggaaggcaaaatgccgcaaaaaa<br>gggaataagggcgacacggaaatgttgaatactcatactcttcctttttcaatattattgaagcatttatcagggttattgtctcatgagcggatacatatttgaat<br>gtatttagaaaaataaacaaataggggttccgcgcacatttccccgaaaagtgccacctgacgtctaagaaaccattattatcatgacattaacctataaaaat<br>aggcgtatcacgaggccctttcgtctcgcgcgtttcggtgatgacggtgaaaacctctgacacatgcagctcccggagacggtcacagcttgtctgtaagcggat<br>gccgggagcagacaagcccgtcagggcgcgtcagcgggtgttggcgggtgtcggggctggcttaactatgcggcatcagagcagattgtactgagagaaagg<br>caattgggtaccgagctcgcggccgcaagcGCGGCCGCGTCTGATCAACCTTCACACAGATCTAGAATCAAAAGCAGTGATCTCCCG<br>GGTGCGAAATAAAAATACTAGGTAACTAGAGGGACTGCGACGTTCCCTCCTATGATTGGTTGTCTTATTACCTTACTTCTAT<br>TATAGTATAACATGTTAAACGATAGTTTGTCTACCCTTTTCGACAAATTGATGATAATAAATAGTATAGGTATATAGTCGTG<br>ATTTAGTTGTTAGATTCTTGTCGAAGATAGTCGGTCAATGGGGAAATGGTGTATGTTGTCGCTGTACCCTACTTTAAAGTA<br>AGCCCCCACCCTCACATGATACCATTCTCCTAATATCGACATAATCCGTCGATCCTCGGCATACCATGATCAGGGAGGGAA<br>ACTACTACTTAATATATCAATCTATAGACCTACTAGATAGGTTTGTCAATGAACAACATAAAACGACACAGAATCCCACGTT |

| | |
|---|---|
| | TTAGCGCTTCGTCTGTGTCGCAGAAGCAGGCACGGTAAGACACGGTCTAGCTGACTGTCTATCGGCTAGGTCAAATAGGG AGCTTTGATATCTGCGTGTCCAGCCTTAGAATTCAGTTCAGCGCGGCCGC |
| pEX-A125-pBnobar | CAAAAACAGGAAGGCAAAATGCCGCAAAAAAGGGAATAAGGGCGACACGGAAATGTTGAATACTCATACTTCTCCCTTTT TCAATATTATTGAAGCATTTATCAGGGTTATTGTCTCATGAGCGGATACATATTTGAATGTATTTAGAAAAATAAACAAATA GGGGTTCCGCGCACATTTCCCCGAAAAGTGCCACCTGACGTCTAAGAAACCATTATTATCATGACATTAACCTATAAAAAT AGGCGTATCACGAGGCCCTTTCGTCTCGCGCGTTTCGGTGATGACGGTGAAAACCTCTGACACATGCAGCTCCCGGAGAC GGTCACAGCTTGTCTGTAAGCGGATGCCGGGAGCAGACAAGCCCGTCAGGGCGCGTCAGCGGGTGTTGGCGGGTGTCG GGGCTGGCTTAACTATGCGGCATCAGAGCAGATTGTACTGAGAGTGCACCAATTGGGTACCGAGCTCGCGGCCGCAAGC GCGGCCGCGAACATCCACACTTTAGTGAATCGAAGCGCGGCTTCAGAATACCGTTTTGGCTACCTGATACAAAGCCCATC GTGGTCCTCAGATATCGTGCACGTAGAGCCTCCTATGATTGGTTGTCTTATTACCTTACTTCTATTATAGTATAACATGTTA AACGATAGTTTGTCTACCCTTTTCGACAAATTGATGATAATAAATAGTATAGGTATATAGTCGTGATTTAGTTGTTAGATTC TTGTCGAAGATAGTCGGTCAATGGGGAAATGGTGTATGTTGTCGCTGTACCCTACTTTAAAGTAAGCCCCCACCCTCACAT GATACCATTCTCCTAATATCGACATAATCCGTCGATCCTCGGCATACCATGATCAGGGAGGGAAACTACTACTTAATATAT CAATCTATAGACCTACTAGATAGGTTTGTCAATGAACAACATAAAACGACACAGAATCCCACGTTTTAGCGCTTCGTCTGT GTCGCATTGCACCGCACGCATGTGGAATTAGTGGCGAAGTACGATTCCAAGACCGACGTACGATACAACTATGCGGATGT GACGAGCTTCTTTTATATGCTTCGCCGCGGCCGCACCTGCTTTTGCTCGCTTGGATCCGAATTCCTGTGTGAAATTGTTATC CGCTCACAATTCCACACAACATACGAGCCGGAAGCATAAAGTGTAAAGCCTGGGGTGCCTAATGAGTGAGCTAACTCACA TTAATTGCGTTGCGCTCACTGCCCGCTTTCCAGTCGGGAAACCTGTCGTGCCAGCTGCATTAATGAATCGGCCAACGCGCG GGGAGAGGCGGTTTGCGTATTGGGCGCTCTTCCGCTTCCTCGCTCACTGACTCGCTGCGCTCGGTCGTTCGGCTGCGGCG AGCGGTATCAGCTCACTCAAAGGCGGTAATACGGTTATCCACAGAATCAGGGGATAACGCAGGAAAAGAACATGTGAGC AAAAGGCCAGCAAAAGGCCAGGAACCGTAAAAAGGCCGCGTTGCTGGCGTTTTTCCATAG |

| Libraries example names | sequences |
|---|---|
| OriLR_epPCR_DNK_WT | aaagtaagcccccaccctcacatgataccattctcctaatatcgacataatccgtcgatcctcggcataccatgatcagggagggaaactactactt aatatatcaatctatagacctactagataggtttgtcaatgaacaacataaaacgacacagaatcccacgttttagcgcttcgtctgtgtcgcaggg cccgtcgactgctaatacgactcactatagggagaccacaacggtttccctctagaaataattttgtttaactttaagaaggagatataccatgtca gtattagtcggtttacatggtgaagctggttcgggaaaagataccgttgcaaaattaattatcgattggtgtaatgacacgtatccaacgtgtttatc ccgtcgttacagttttgctaagcccgtttatgaacttgcatccgtaatcctcggcgtaactccagagtttctaggagagcgcaggggggaaagagatt gaccaatggtttacggtcacacaatctcaattggagcgagctagagacgtatggtttaaatacggtatcgataagtttgaagacttctcgtacgtttg gccgatttttgaggaaaaatatcttaatcctcaacaacttatctcagagaataaagaagacggactttatagtttatttatttctccgagaaaaatgt tacagctcgtaggaacggagctaggaaggcagctggtacatgaacgcatttggctcataattctggagcaatccatcgctaaagacgacccagat gtcgccgtaataacagatgtgagattccccaatgaaggagagttactcagagagacgaaccatttagatatggattctttactagtaaatgtagtg cccgctgagcagaagttcactattaaatcagatcatccatctgaaagtggtattcctgcaaaatatatcactcacgaattagttaataaattcgatg gtatcaataaccttaaactagaagtgtataacttctgtgacttagagctagaaccactagttggataaatagtcaatcttcggaaatcccaaatcta ctgggactcaaaatnnnnnnnnnnnnnnnnnnnnnnnnnnnnnntgactagttaggatctgtcgctgaaaggcttctaatagcataaccc cttggggcctctaaacgggtcttgaggggtttttttgcctcctatgattggttgtcttattaccttacttctattatagtataacatgttaaacgatagttt gtctacccttttcgacaaattgatgataataaatagtataggtatatagtcgtgatttagttgttagattcttgtcgaagatagtcggtcaatggggaa atggtgtatgttgtcgctgtaccctacttt |
| Y_OriLR_12_RBS-1_WT | AAAGTAAGCCCCCACCCTCACATGATACCATTCTCCTAATATCGACATAATCCGTCGATCCTCGGCATACCATGAT CAGGGAGGGAAACTACTACTTAATATATCAATCTATAGACCTACTAGATAGGTTTGTCAATGAACAACATAAAA CGACACAGAATCCCACGTTTTAGCGCTTCGTCTGTGTCGCATTGCACCGCACGCATGTGGAATTAGTGGCGAAG TACGATTCCAAGACCGACGTACGATACAACTATGCGGATGTGACGAGCTTCTTTTATATGCTTCGCCGTCTCGGC AANNWNNNNNWNNGCAAGCTCTTCGATGTGGAGGTTCAGAATCACTGACGCGATTGTAATACGACTCACTAT AGGgataatACAACGGTTtccctctagaaataattTTGTTTAACTTAGGGCCCGTTAGGTTCCGATGgttagtaaaggtgaag aggataacatggccatcatcaaggagttcatgcgcttcaaggtgcacatggagggctccgtgaacggccacgagttcgagatcgagggcgaggg cgagggccgcccctacgagggcacccagaccgccaagctgaaggtgaccaagggtggccccctgcccttcgcctgggacatcctgtcccctcagtt catgtacggctccaaggcctacgtgaagcaccccgccgacatccccgactacttgaagctgtccttccccgagggcttcaagtgggagcgcgtgat gaacttcgaggacggcggcgtggtgaccgtgacccaggactcctccctgcaggacggcgagttcatctacaaggtgaagctgcgcggcaccaact tcccctccgacggccccgtaatgcagaagaagaccatgggctgggaggcctcctccgagcggatgtaccccgaggacggcgccctgaagggcga gatcaagcagaggctgaagctgaaggacggcggccactacgacgctgaggtcaagaccacctacaaggccaagaagcccgtgcagctgcccgg cgcctacaacgtcaacatcaagttggacatcacctcccacaacgaggactacaccatcgtggaacagtacgaacgcgccgagggccgccactcc accggcggcatggacgagctgtacaagGGTGGCGGTGGCAGTTTAGTGCCACGTGGTTCAGGCGGTGGCGGTTCTATGT CAGTATTAGTCGGTTTACATGGTGAAGCTGGTTCGGGAAAAGATACCGTTGCAAAATTAATTATCGATTGGTGT AATGACACGTATCCAACGTGTTTATCCCGTCGTTACAGTTTTGCTAAGCCCGTTTATGAACTTGCATCCGTAATCC TCGGCGTAACTCCAGAGTTTCTAGGAGAGCGCAGGGGGAAAGAGATTGACCAATGGTTTACGGTCACACAATC TCAATTGGAGCGAGCTCGTGACGTATGGTTCAAATACGGTATCGATAAGTTTGAGGACTTCTCGTACGTTTGGC CGATTTTTGAGGAAAAATATCTTAATCCTCAACAACTTATCTCAGAGAATAAAGAAGACGGACTTTATAGTTTAT TTATTTCTCCGAGAAAAATGTTACAGCTCGTAGGAACGGAGCTAGGAAGGCAGCTGGTACATGAACGCATTTG GCTCATAATTCTGGAGCAATCCATCGCTAAAGACGACCCAGATGTCGCCGTAATAACAGATGTGAGATTCCCCA ATGAAGGAGAGTTACTCCGTGAAACGAACCATTTAGATATGGATTCTTTACTAGTAAATGTAGTGCCCGCTGAG CAGAAGTTCACTATTAAATCAGATCATCCATCTGAAAGTGGTATTCCTGCAAAATATATCACTCACGAATTAGTT AATAAATTCGATGGTATCAATAACCTTAAACTAGAAGTGTATAACTTCTGTGACTTAGAGCTAGAACCACTAGTT |

| | |
|---|---|
| | GGATAAATAGTCAatcttcggaaatcccATGCCTCTAATACTACACACTTAGCATAACCCCTTGGGGCCTCTAAACGG<br>GTCTTGAGGGGTTTTTTGTACTGTAGTAATCATGTTACCACAGTGAATGCGAAGAGCTGCCNNWNNNNNWNN<br>TGCCCGAGACGGAACATCCACACTTTAGTGAATCGAAGCGCGGCTTCAGAATACCGTTTTGGCTACCTGATACA<br>AAGCCCATCGTGGTCCTCAGATATCGTGCACGTAGAGCCTCCTATGATTGGTTGTCTTATTACCTTACTTCTATTA<br>TAGTATAACATGTTAAACGATAGTTTGTCTACCCTTTTCGACAAATTGATGATAATAAATAGTATAGGTATATAG<br>TCGTGATTTAGTTGTTAGATTCTTGTCGAAGATAGTCGGTCAATGGGGAAATGGTGTATGTTGTCGCTGTACCCT<br>ACTTT |

**Supplementary note 1.** Estimation of dNTP production in early phase

According to Mikoulinskalia (Mikoulinskaia et al., 2013) the activity of wild type DNK towards phosphorylation of dNMP in *in vitro* conditions if 41.24 ± 1.77 U/mg. Considering DNK molecular weight is 28700 Da and one unit is defined as the amount of the enzyme catalyzing the transformation of 1 µmol monophosphate in 1 min at 25°C, one can calculate that one enzyme activity is $2.10^{-16}$U, or one enzyme transforms $2.10^{-16}$ µmol monophosphate per minute. Droplet volume being around $2.10^{-12}$ L, one enzyme increases the dNDP concentration (in initial conditions) of $1.10^{-4}$ µM per minute, or $6.10^{-3}$ µM per hour. It would take 166 molecules to produce 1 µM of dNTP in one hour. Since dNTP is necessary for replication, with typical polymerase having a $K_M$ in the low micromolar range (1-50 µM), and that the DNK enzymes need to be produced from a discrete number of gene copies, we expect a significant delay before the actual replication can kick-start.

**Supplementary note 2.** Decrease of IVTTR expression and replication capabilities over incubation time

The replication proteins DNAP and TP are produced by the PURE system during incubation. As the accumulation of dNTP resources is also gradually increasing due to DNK activity, we deemed necessary to observe the effect of incubation time on replication capacity of the system.

We designed a <*gfp*> replicator assay to observe the effect of delay of dNTP addition on replication, in bulk conditions. The <*gfp*> replicator was obtained by amplifying the gene encoding for GFP flanked by origins of replication in pUC_OriLR_gfp with phosphorylated primers T22/T23 and Q5 polymerase using program **PCR 24**. We mimicked the delayed appearance of deoxynucleotide triphosphate by spiking dNTP at different time point. We then recorded GFP fluorescence at plateau, and quantified <*gfp*> replicator concentration by qPCR, shown in Supplementary Figure 15. All conditions share these same components: PUREfrex 1.0 diluted as in manufacturer's protocol, $AmSO_4$ 20 mM, P5 0.375 mg/mL, P6 0.105 mg/mL, <*gfp*> 12 pM, and different concentrations of pUC_OriLR_p2p3 of 50 pM, 100 pM and 200 pM indicated on the figure. The spike-in times were 0, 30 min, 1 h, 2 h, and 3 h. 4 µL of reaction mixture without dNTP was aliquoted in an optical PCR microtube (white, low profile, BioRad) and 1 µL of dNTP mixture of different composition was added to a final concentration of 0.3 mM at the indicated time. Reactions were incubated in a real-time thermocycler (CFX Touch BioRad) at 33 °C for 15 h while monitoring fluorescence in all channels. After incubation, the sample was diluted $1/1000^{th}$ in milliQ water and <*gfp*> was quantified by qPCR using primers T129/T130.

Whereas the expression capacity at T0 decreases with *p2p3* concentration (*p2p3* expression burden). It also decreases with time for every *p2p3* concentration, but we clearly see that after 1 h, only 50 pM *p2p3* condition can still produce high amount of GFP after dNTP addition. This result suggests replication capacity.

We can observe that if the dNTP spike-in is added at T0, the yield is 100 nM to 200 nM and at 30 min the yield is around 70-80 nM. At 60 min the biggest discrepancy is visible: *gfp* concentration is 58.5 nM with 50 pM *p2p3*, 6.78 nM with 100 pM *p2p3* and 2.54 nM with 200 pM *p2p3*. This shows that P2 and P3 accumulation is not useful for replication if it happens too late: it is better to synthesize P2 and P3 regularly. This seems to indicate that these proteins are not stable during incubation without replication.

Without pUC_OriLR_p2p3, GFP expression capacity slowly decreases with time, reaching half of the starting capacity at about 4.5 h for PUREfrex 1.0 and between 3 h and 4.5 h for PUREfrex 2.0. The expression capacity in presence of 200 nM of pUC_OriLR_p2p3 (expression burden) decreases faster, being half of the starting capacity at about 30 min for PUREfrex 1.0 and less than 30 min for PUREfrex 2.0.

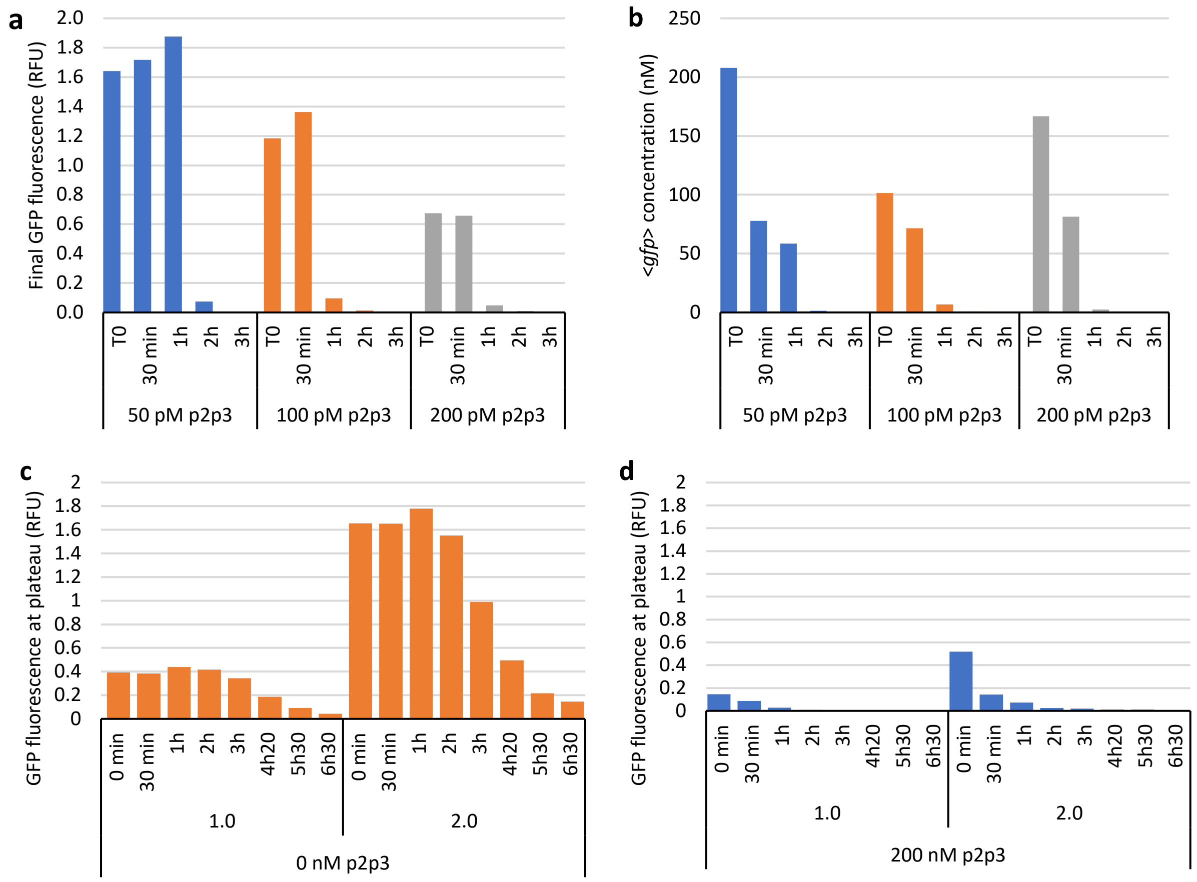


**Supplementary Figure 15 a,b.** Endpoint IVTTR results with PUREfrex 1.0 after timed addition of 0.3 mM dNTP (time written under the bars) for different concentrations of DNA pUC_OriLR_p2p3. a. GFP fluorescence at plateau (RFU). b. qPCR quantification of <*gfp*> gene concentration at plateau. **c**,**d**. GFP Fluorescence at plateau after addition of 2 nM of non-replicative pUC_OriLR_gfp in IVTTR system preincubated at 33 °C for a given time, in absence (orange) or presence (blue) of 200 nM pUC_OriLR_p2p3, creating a burden on PURE machinery.

**Supplementary note 3.** Determination of DNA copy number per droplet**.**

Sample preparation and processing is important to quantitatively analyze the UMI distribution. One microliter of emulsion contains 0.7 µL of aqueous solution considering a droplet packing of 70% (between random and optimal (C. Song et al., 2008; Wu et al., 2003)), translating to $2.4 \times 10^5$ initial DNA molecules for 2 pL droplets and a lambda of 0.7.

Here, one microliter of the emulsion was taken and diluted in 16 µL milliQ water then dialyzed, giving around 27 µL after swelling due to dialysis.

The total DNA concentration in this 27 µL sample was measured by qPCR at 2.86 pM, corresponding to $4.7 \times 10^7$ molecules. One microliter of this pool is used for PCR and the PCR product is sequenced, giving $2.6 \times 10^5$ reads. Thus, the sequencing was performed on $4.7 \times 10^7 \times 1/27 = 1.7 \times 10^6$ molecules. Overall, the relationship between a cluster of size S >> 1 and the total number of molecules from the initial 27 µL (and so from the droplet) is:

$$\frac{S}{2.4 \times 10^5} \times 4.7 \times 10^7$$

A cluster of size 20 thus corresponds to 130 molecules in the subpool, or 3600 copies in its droplet of origin.

**Supplementary note 4.** Effect of PUREfrex 2.1 DTT or GSH on selection or replication

Because changes in redox chemistry can affect protein folding and activity in the PURE system, we tested both conditions in parallel. The control reaction containing all four dNTP was performed with GSH as a reducing agent instead of DTT. The selection conditions, in which dTTP was replaced by dTTP, were tested in the presence of GSH or DTT. In the GSH-selection emulsion no fluorescence was observed, indirectly indicating no DNA replication. As fluorescence was observed with the positive control, the GSH is not toxic for replication machinery nor mCherry protein. Reducing power may be important for the structure and function of DNK enzyme. Indeed, DNK possesses 3 cysteine residues: C26 and C31 are oriented toward each other may form a disulfide bridge, while C241 is buried in the structure and is a conserved residue according to the error prone library analysis.

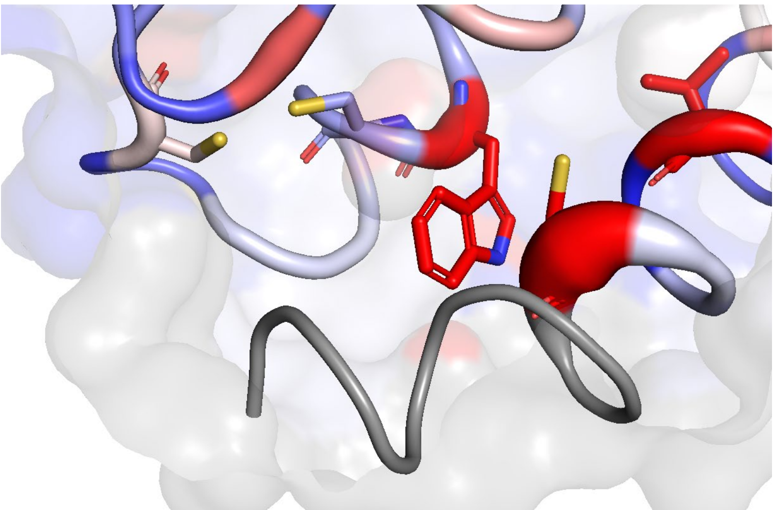

**Supplementary Figure 16.** Cysteine residues on the AlphaFold3 structure of DNK, in PyMOL putty cartoon representation with tube radius and color change according to the relative entropy of the residue (blue: 0; red: 0.02). From left to right, C33, C26 and C241. C26 and C33 possibly form a disulfide bridge while C241 is a conserved residue.
